# Dispersal-behavioral plasticity within an insect-host system undergoing human-induced rapid environmental change (HIREC)

**DOI:** 10.1101/2022.06.19.496739

**Authors:** Anastasia Bernat, Meredith Cenzer

**Affiliations:** University of Chicago, Department of Ecology & Evolution, Chicago, IL 60637

**Keywords:** flight dispersal, phenotypic plasticity, sexual dimorphism, host adaption, morphology, HIREC

## Abstract

As environments change, often drastically, due to human activities, dispersal-behavioral plasticity can become a key mediator of metapopulation connectivity and the interactions between an individual and its altered environment. Our goal was to investigate the traits and ecological processes that affect plastic dispersal responses within an insect-host system undergoing human-induced rapid evolutionary change (HIREC). Since the 1960s, populations of the red-shouldered soapberry bug from Florida, USA, originally feeding on the seeds of *Cardiospermum corindum* on the islands, quickly adapted to feeding on an invasive, ornamental tree, *Koelreuteria elegans*, on the mainland, which led to host-specific wing polyphenism. Here, we measured the morphology of >3,500 soapberry bugs field collected from 2013 to 2020 and the flight performance of 378 total soapberry bugs tested in a flight mill during Fall 2019 and Winter 2020. Flight tests showed females and mainland natives exhibited variable flight responses mediated by mass, while males were consistent, wing-dependent flyers. However, historical specimens showed annual rises in flightless morphs for males and dwindling wing-to-body sizes for island natives since 2013. Despite uncertain future fitness consequences, plasticity could help predict mobility character and agent dispersal behavior and ultimately help identify whether recent trends signal adjustment or maladaptation to HIREC.

## 1 Introduction

In the past decade, several ecologists have called to action the need to translate adaptive plasticity into explicit science not only to better understand how organisms make decisions in response to their environment but also to redefine behavioral ecology as a discipline [1,2,3,4,5]. The call was ushered in response to a rise in consciousness since the 1990s [6,7,8,9] about humans as an evolutionary force and its irrefutable, accelerating discord with the field’s approach to studying animal behavior within the environmental conditions that animals originally evolved in [10]. As shown, human-induced rapid environmental change (HIREC) has stemmed from rapid discontinuities with our environmental and evolutionary past, as direct and indirect results of human activities have been inundating organisms with novel conditions, leading biota to either sink [11,12] or adapt [13,14] but mostly sink [15]. In HIREC, these novel conditions facilitated by humans cause great or significant changes in evolutionary potential, from directional selection to genetic or plastic expression, and they facilitate these changes at speeds faster than organisms’ evolutionary past. In many cases, organisms are not doing poorly with HIREC because they cannot respond fast enough but because they show maladaptive behavioral responses, such as those associated with ecological or evolutionary traps [16]. From insects attracted to laying their eggs on artificial surfaces (e.g. oil or roads) resulting in their death [17,18] to ‘silent’ insect mass extinctions [19], there seems to be an approaching homogeneous species bottleneck, and yet there is a growing consensus that plasticity could act as both a means for organisms to weather and resist HIREC [20] and as a blueprint for humans to reconstruct ethology.

When defined conservatively, plasticity is the expression of multiple phenotypes from one genotype when exposed to different environments [21,22]. Plasticity can affect all levels of ecological organization, but its most pertinent influence has been its link to environmental change and evolutionary response. A meta-analyses comparing the rates of phenotypic change across 68 animal systems suggested that, for systems undergoing HIREC, plasticity contributed up to twice as much to phenotypic change as genetic effects [23]. Additionally, among plants and invertebrates, increases or decreases in plasticity following anthropogenic disturbances were shown to be directed by trait type and taxon [24]. Although assessing the degree in which plasticity is expressed is important, what is largely missing in the HIREC literature are patterns of behavioral plasticity and their trait dependencies within a species [25]. Already, researchers have demonstrated how generalists and diet-expansive species can raise their chances of coping with extinction [26]. Furthermore, it has been shown how behavioral plasticity can be used to help predict the adaptability of a species undergoing habitat degradation due to reductions in spatial complexity [27] or urbanization resulting in spatial fragmentation [28]. Others have demonstrated how behavioral plasticity can be confounded by body size [29] and habitat breadth [30]. In turn, the need to test individual plastic behaviors in response to HIREC has become more prevalent alongside the demand to demonstrate how plastic-behavioral traits link to other, slower-evolving processes, like adaptions in life history or morphology. For insects and their behavioral responses involving diapause or threshold traits like polyphenisms and polymorphisms, anthropogenic environmental sources of variation in transgenerational and maternal effects have been more documented [25,31]. This has opened avenues to study the plasticity of more allusive and intractable forms of behavior like dispersal.

However, plastic expressions of various acts of dispersal can be challenging to study by behavioral ecologists, despite dispersal occurring ubiquitously across many biological systems. Dispersal ability, the potential for movement by animals between locations [32], is a fundamental trait for determining geographic range and gene flow within and between populations [24] and it can be dampened [33,34] or induced by persistent deleterious changes in habitat conditions, particularly for philopatric organisms [35]. Individuals also display various dispersal characteristics, such as endurance, periodicity, and speed, and these traits have been linked to morphological traits like sex [36], body size [37,38], and wing size [39] as well as trade-offs with reproduction[40]. However, capturing movement, especially plastic behaviors in movement, requires reliable tracking methods or machinery, which has led to an explosion of flight mill contraptions in the past six decades for insect flight dispersal quantification [41,42,43]. For ecologists that have taken the approach to empirically capture dispersal, the advantage of studying dispersal-behavioral plasticity arises not only when we consider why or how a species disperses but also ask how fixed or plastic acts of dispersal have evolved over time. For species undergoing HIREC, we can further assess how variability in dispersal can facilitate adaptive or maladaptive strategies in response to HIREC and identify who the main contributing dispersal agents in a population are. Especially considering that acts of dispersal can be directly initiated by individuals, measuring agent dispersal behavior, as opposed to population genetics, can reveal mobility characteristics that have not necessarily been genetically assimilated yet [44]. This could ultimately better help us understand how environmentally-induced phenotypic variation links to a population’s evolutionary history or guides evolutionary trajectories of adaption or maladaption in response to HIREC. Thus, evidenced patterns of dispersal-behavioral plasticity can help us better infer the future connectivity and survival of populations.

In this paper, we examined how, under an insect-host system undergoing HIREC, plasticity in dispersal behavior could help predict mobility character and agent behavior as it depends on an organism’s evolutionary history. As such, we studied the soapberry bug collected from Florida and its host plants (*Cardiospermum corindum*, native; *Koelreuteria elegans*, invasive) as a promising system for assessing the connections between HIREC and dispersal-behavioral plasticity. As observed from field collections, the soapberry bug is a seed-eating and sexually dimorphic insect that has adapted to an invasive, ornamental tree within the past half-century [45,46]. This rapid evolutionary change has led to host-specific wing polyphenism [47,48]; adults will either develop long forewings with flight muscle (“long-wing” volant morph), long forewings without flight muscle or with histolyzed flight muscle (“cryptic” flightless morph), or a truncated wing without flight muscle (“short-wing” flightless morph). These flightless morphs have been found at higher frequencies on *K. elegans*, a host plant that synchronously produces ample volumes of seeds starting in the Fall season, compared to *C. corindum* that variably produces fewer seeds throughout the year [47]. Furthermore, each host is spatially separated between the mainland (invasive) and the islands (native) of Florida, which peripherally introduces questions about range-edge dynamics [49,50]. With such spatial diversity, including observed latitudinal clines in beak length [51], ocean barriers, and mainland urbanization, we considered how populations deeper in the mainland as opposed to those at the sympatric zone (i.e. the geographic area where two host plants overlap) or in the islands were dispersing, and how plastic flight behaviors could inform slower-evolving traits like wing morphology and wing morph frequencies. Furthermore, considering the interconnected evolution of sexual dimorphism within the species [52], we assessed how sexual asymmetry in morphology leads to sex-biased dispersal [53,54] and how its ecological consequences will depend on our ability to determine its prevalence and plastic variation in natural populations. Thus, these patterns of plasticity in a behavior as morphological, mechanistic, metabolic, and spatial like insect flight movement [55,56,57,58,59] can also provide insights into how behavioral traits are confounded with weight, sex, or local geographic variation.

In our study, we asked which traits and ecological processes affect plastic dispersal responses and which agents in a population are driving those responses? We also asked whether plasticity could help predict mobility character and frame how insects are coping or not coping with HIREC? In turn, we measured the morphology of >3,500 soapberry bugs field collected from 2013 to 2020 and recorded the flight behavior of 378 total soapberry bugs using a flight mill during Fall 2019 and Winter 2020. We found novel sex-specific and host-specific plastic flight responses, repeatedly mediated by mass changes. Flight tests showed high variability in flight response between trials for females and a latitudinal cline in flight response for mainland natives from *K. elegans*, suggesting trade-offs among flight potential, reproduction, and host plant. Conversely, males and island natives from *C. corindum* were consistent, homogeneous flyers; males were 2.5x more likely to fly than females and were 1.3x more likely to be long-winged. However, historical specimens showed annual rises in flightless morphs for males since 2013. Soapberry bugs collected from *C. corindum* since 2013 also showed dwindling wing-to-body sizes annually. These dependencies on sex, host plant, and mass suggest that a species ability to cope well with future environmental variation will depend on differing or biased dispersal agent responses in a population. Furthermore, our findings on the plastic effects of latitudinal distance from the sympatric zone, provide insight into how degrees of insularity or confluence direct whether a trait exhibits a more fixed or variable effect on dispersal abilities. Little is known about the traits and spatio-temporal processes that affect plastic dispersal responses, but we offer a complex dispersal behavioral system that exhibits sex-specific and host-specific plastic responses as well as potential maladaptive consequences. As we learn more about the human influences that have inundated environmental changes at scales larger and frequencies faster than recent pre-human phenomena have, we will come to better understand and predict the eco-evolutionary behaviors and plastic responses engendered by HIREC.

## 2 Methods

### 2.1 Study Species 2013-2020 Field Collection and Morphology

Soapberry bugs were collected in 1 to 8 locations in Florida at least once a year from April 2013 to February 2020 (see Appendix A Section 2.5). Each seasonal bug collection contained a mix of soapberry bugs selected from their native host plant, *Cardiospermum corindum*, and nonnative host plant, *Koelreuteria elegans* (Figure 1A). After shipment, collected bugs were either preserved immediately in ethanol in 50 mL falcon conical centrifuge tubes grouped by population or preserved individually, with a unique identification number, in ethanol in 2.0 mL microcentrifuge tubes following laboratory experiments. We took morphological measurements for each soapberry bug collected, using Mitutoyo digital calipers and a Zeiss Stemi 1000 Microscope 7x – 35x. Measurements of interest included the distance from the rostrum to the tip of the wing (body length) and forewing length (see Appendix A Section 2.4 for sample microscope images).

**Figure 1:**
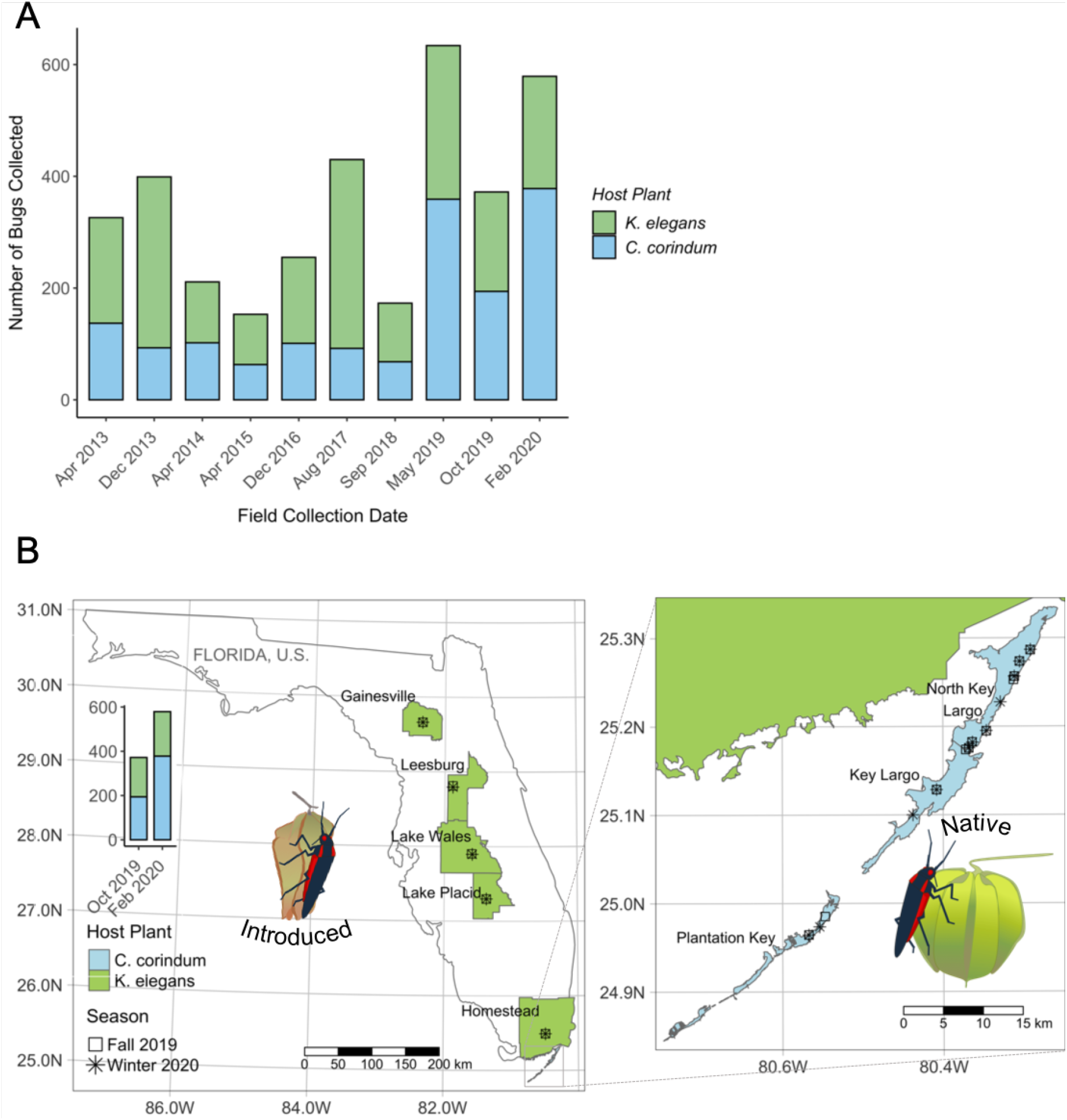
Soapberry bug collection by host plant and season across Florida U.S. from 2013 to 2020. **A.** Stacked barplot of the number of soapberry bugs collected by season and host plant from April 2013 to February 2020. Bars shaded in green represent bugs collected from *K. elegans* and bars shaded in blue represent bugs collected from *C. corindum*. **B**. Map of Fall 2019 and Winter 2020 soapberry bug collections tested for flight trials. Collection sites are marked by season where squares represent Fall 2019 collection sites and stars represent Winter 2020 collection sites. Population names are displayed and named after the nearest city to the collection site(s). Counties shaded in green on the mainland represent bug collections from *K. elegans* and counties shaded in blue in the keys represent bug collections from *C. corindum*. The stacked barplot on the left of Florida graphs the number of soapberry bugs collected and tested for Fall 2019 (*n* = 203) and Winter 2020 (*n* = 332) among each host plant. Images of a soapberry bug feeding on the seeds inside the pods of *K. elegans* (left) and *C. corindum* (right) were illustrated by AVB.

### 2.2 Study Species 2019-2020 Field Collection and Flight Trials

#### 2.2.1 Field Collection

For flight trials, soapberry bugs were collected during the Fall 2019 and Winter 2020 season in eight locations in Florida and shipped to Chicago, IL. In the three archipelago sites, bugs were collected from *C. corindum*; in the five mainland sites, bugs were collected from *K. elegans* (Figure 1B). To see exact collection site names and counts for each season, refer to the tables in Appendix A Section 2.5.

#### 2.2.2 Flight Trial Initial Preparation and Care

Of the soapberry bugs collected during Fall 2019 and Winter 2020, 207 and 476 bugs survived shipment, respectively. All except 147 short-winged bugs collected in Winter 2020 were placed into assembled bug homes with an assigned unique identification number (see Appendix A section 2.4 comparing a short-winged adult to a long-winged adult). Bug homes were assembled using a plastic soufflé cup (4 oz) that was sealed with a mesh lid and lined with filter paper at the base. Each cup contained a 2.0 mL microcentrifuge tube filled with deionized (DI) water and stoppered with cotton. Each bug was provided initially with two different seeds, one from *K. paniculata* (Sheffield’s Seed Company) and one from *C. halicacabum* (Outsidepride.com, Inc), two congeners of the field host plants that are commercially available. The upper rim of each plastic souffle cup was lined with PTFE Fluoropolymer (“Fluon”), creating a slippery surface to keep bugs primarily in the bottom of the cup (Figure 2B). Identified soapberry bugs in bug homes were then randomly assorted into boxes and placed into an incubator. The incubator was illuminated with Philips 17 W, 24 in florescent lighting and was set at 28 C/27.5 C (day/night), 70% relative humidity, and a 24 hour light/0 hour dark cycle.

**Figure 2:**
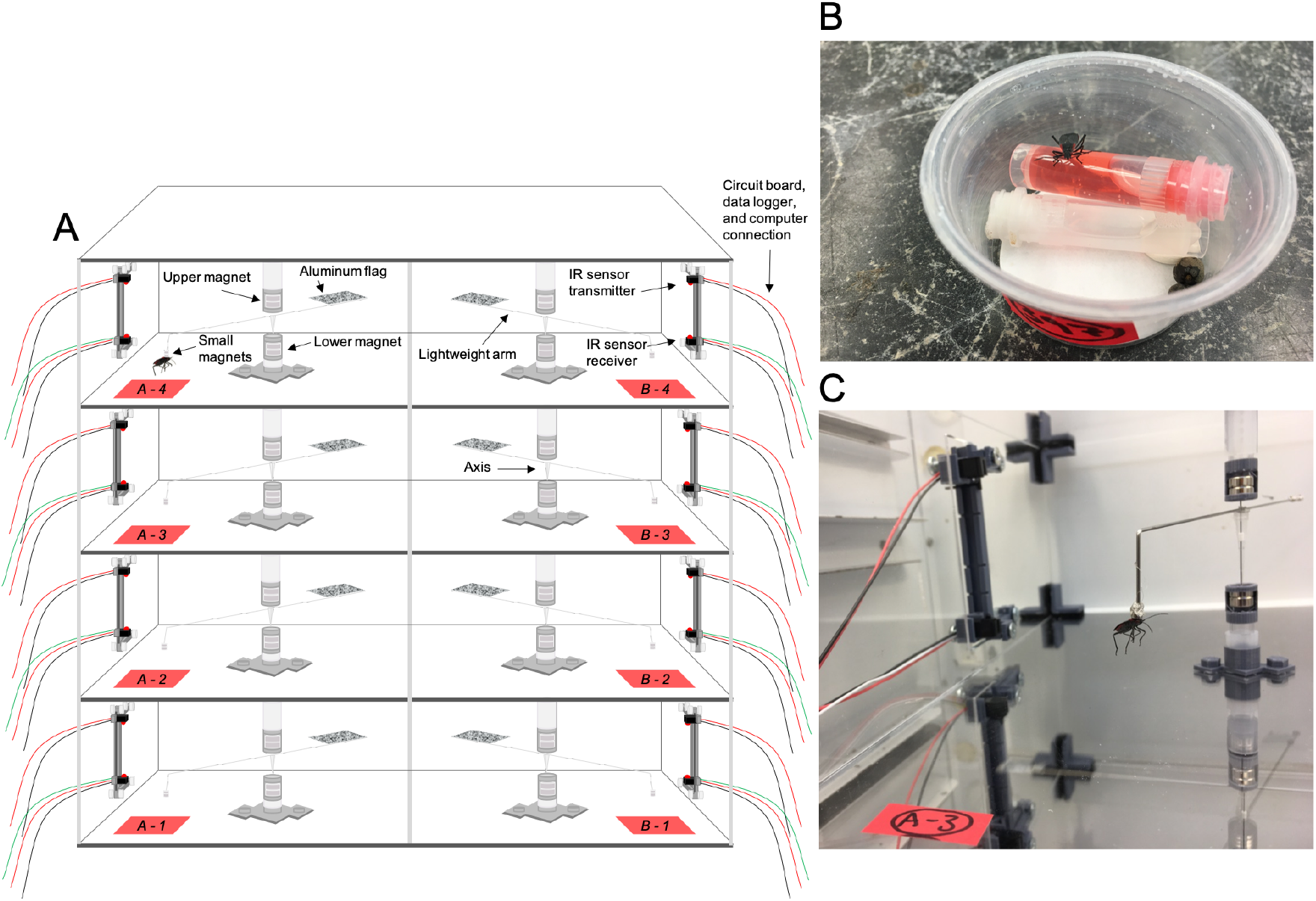
Flight mill machine and assembled bug home. **A**. Diagram of the flight mill, **B**. photograph of assembled and prepped bug home, and **C**. photograph of a stationary mounted soapberry bug in chamber A-3. Illustration made by and photos captured by AVB.

#### 2.2.3 Repeated Bug Preparation and Care

Daily, dead soapberry bugs were pulled from their assorted homes and preserved in ethanol; then, morphological measurements were taken (n =683) that followed the aforementioned methods in Section 3.1. Weekly, adult soapberry bugs received general care: their filter paper was replaced, microcentrifuge tubes were refilled with DI water and capped with a new piece of cotton, and two fresh seeds (one from each host plant) were added to each bug home. In addition, once a week, eggs laid by female bugs were collected and discarded. Eggs were only counted during Winter 2020 trials.

#### 2.2.4 Flight Mill Trials

Fall flight trials were conducted over a 27 day period in October and November of 2019. After confidently confirming that no short-wing bugs fly following different flight trial lengths, we removed all short-wing bugs from the testing pool and tested the remaining long-winged bugs for an unlimited trial length from November 5 to November 8. An unlimited trial length is defined as a trial where bugs remained on the flight mill until they stopped flying of their own volition. The unlimited trial model most successfully captured the full scope of soapberry bug flight potential, distance, and speed. The unlimited trial experimental setup was repeated for Winter flight trials.

Winter flight trials were conducted over a 22 day period in February and March of 2020. We ran two sets of trials, the first from February 17 to February 28 on 332 bugs, the second from March 3 to March 10 on the surviving 282 bugs. Refer to Table 1 for a summary of each season’s experimental setup. Only unlimited trials are described below.

1. Flight Trial Preparation For both seasons, we randomly ordered bugs by their identification number for each set of flight trials. The day before their trial, soapberry bugs received sugar water in a 2.0 mL microcentrifuge tube made from 7 volumes of Fruit Punch Gatorade and 3 volumes of DI water and capped with cotton. We also painted soapberry bugs’ pronotums with Sophisticated Finishes^*™*^ Iron Metallic Surfacer paint and set the painted bugs to dry in their bug homes overnight.
2. Running Flight Trials The day of their flight trial, we assigned a flight mill chamber to each soapberry bug based on their randomized trial order. The flight mill supported eight individual plexiglass chambers (Figure 2A). Within each chamber, we magnetically tethered a painted bug to the small N42 neodymium magnets of a magnetically suspended pivot arm. When the insect flew, the flight mill recorded its flight revolutions using a DATAQ Model DI-1100 data logger and the WinDAQ Acquisition Waveform Recording and Playback Software. After trials, we processed the recording files into flight distance and instantaneous speeds (see Bernat 2021 for details on flight mill construction, functionality, and programming). The flight mill was located in an incubator set to the same conditions as the rearing incubator except it ran a 14 hour light/10 hour dark cycle (sunrise at 8 AM and sunset at 10 PM). Before flight trials, Fall bugs (*n* = 207, number of mass measurements = 337) and Winter bugs were massed (*n* = 332, number of mass measurements = 611) with an analytical balance (0.1±mg). For each trial, flight behavior for bugs was recorded every hundredth of a second (sample rate) by the WinDAQ Software for up to 24 hours. We manually recorded flight response, which was defined as whether a bug flew (demonstrated rapid wing movement and self-propelled forward motion) or not. If the bug flew, we recorded the type of flight it displayed. We split flight type into two categories: ‘bursting’ and ‘continuous’ flight behavior. An individual was ‘bursting’ when it flew but did not maintain flight behavior for at least 10 minutes while longer flight was labelled as ‘continuous’. The majority of bursting flight was for very short intervals (< 30 seconds). During the trial period, individuals could display both bursting and continuous flight. We recorded on paper and via Event Markers in the WinDAQ Software the start and stop time for all bugs. The procedure for unlimited recording was such:

a. We began flight trials between 8am and 9am each day. To initiate flight trials, we magnetically tethered massed bugs onto each of the 8 flight mill arms. For each channel, we entered Event Markers with the bug’s ID into the flight recording. We would then blow on each individual to motivate flight.
b. We made two additional motivational attempts at 10 minutes and 20 minutes into each individual’s flight trial. If the bugs were not exhibiting continuous flight by the 30 minute mark, we pulled those bugs off the mill and returned them to their individual bug homes.
c. Bugs who exhibited continuous flight by the 30 minute mark were left on the flight mill and checked on every 30 minutes. If a bug stopped flying, its stop time was recorded and it was pulled from the flight mill and returned to its bug home.
d. Vacant chamber(s) were filled by the next bug(s) that followed in their randomized trial order. We continued adding new bugs and entering their accompanying Event Marker until 5 PM each flight day. Bugs that were still flying after 5 PM were left on the flight mill until the following morning.

**Table 1:** Experimental setup of Fall 2019 and Winter 2020 flight trails.

| Season | Year | Dates | Experimental Setup | N Tested | Wing Morph |
| --- | --- | --- | --- | --- | --- |
| Fall | 2019 | October 15 to October 23 | 30 min | 182 | short and long |
| Fall | 2019 | October 23 to October 29 | 60 min | 130 | short and long |
| Fall | 2019 | October 30 to November 4 | 90 min | 126 | short and long |
| Fall | 2019 | November 5 to November 8 | unlimited | 66 | long |
| Winter | 2020 | February 17 to February 28 | unlimited | 332 | long |
| Winter | 2020 | March 3 to March 10 | unlimited | 282 | long |

### 2.3 Statistics

All statistical analyses were conducted in R (R Core Team, 2020). Analyses included multiple R scripts that conducted regressions for bug morphology from 2013-2020 and for flight response across and between flight trials. Model selection was based on Akaike information criterion (AIC) and an analysis of variation (ANOVA) test where p >0.05 favored the simpler model. Our models included four predictor variables at most that were based on a priori hypotheses; therefore, we assumed low family-wise error. See appendices and publicly available scripts (https://github.com/mlcenzer/SBB-dispersal/tree/master/avbernat).

#### 2.3.1 Morphological Analyses From 2013-2020 Collections

Our morphological traits of interest were primarily concerned with the wing of the soapberry bug. Because we expected all morphology measurements to be correlated with the overall size of the soapberry bug, we calculated the ratio of wing length to body length (wing-to-body).

Because wing-to-body ratios and long-wing morph frequencies were recorded by month and year, we first tested for stationarity using the augmented Dickey-Fuller (ADF) test. After finding no significant temporal dependencies, we concluded that the values from each collection date were independent of each other (i.e. AR(0) process) for both wing-to-body ratio and long-wing morph frequency. We then performed multiple regression analyses to identify which predictors (sex, host plant, month, year, and all possible pairwise interactions) best predicted these two morphological traits, following the aforementioned model selection process. Finally, for visual purposes, we fit a local polynomial regression to each data set, using the lowess and geom_smooth functions in R. We applied smoothers with increasing weights until the residuals appeared to have constant variance. In turn, all local polynomial regressions were fit with a span of 0.4 (*α*, the smoothing parameter), a degree of zero (λ), and 95% confidence intervals. The locally-weighted scatterplot smoothing (LOESS) helped depict the non-linear fluctuations in wing-to-body ratio and long-wing morph frequency across time. See Appendix A for a tidy version of the analyses.

#### 2.3.2 Across-Trial and Between-Trial Flight Response From Fall 2019 and Winter 2020 Collections

To prepare the data for analyses, the recording files were manually converted and processed via three main python scripts (see Bernat 2021). We then ensured that all flight recordings matched our handwritten records of whether a bug flew or not during its trial.

Across flight trials, we performed multiple regression analyses to test whether mass, host plant, wing-to-body ratio, or latitudinal distance from the sympatric zone was associated with either sex’s flight probability (Appendix B section 2.5). Because soapberry bugs exhibit sexual dimorphism, we hypothesized that the sexes may exhibit differing effects on flight potential, so we analyzed the sexes separately.We also applied single-variate regressions to test for any experimental effects, such as days elapsed since the start of trials (Appendix B section 2.4).

Between flight trials, we conducted multinomial logistic regression models to predict the probabilities of different possible flight outcomes as a function of sex, wing-to-body-ratio, and mass percent change (Appendix B section 3.5). There were four flight outcomes possible between trials, which we termed as ‘flight cases’. Each flight case describes a soapberry bug’s change in flight response between trial 1 (T1) and trial 2 (T2): a bug either flew in both trials, only in T2, in neither trial, or only in T1 (Appendix B section 3.4). We also ran multinomial logit models for a subset of the data with females only in order to assess the impact of egg production as a predictor (Appendix B section 3.6). For females, we predicted various flight cases using mass percent change, wing-to-body ratio, and ‘egg case’. Egg case denotes the change in egg laying activity between trials: either a female laid eggs in both trials, only in T2, in neither trial, or only in T1 (Appendix B section 3.6.1). For all flight trial datasets, we used maximum likelihood to estimate multinomial logit models through the nnet library in R. Model selection followed the same aforementioned process. Each best fit model allowed us to then calculate the predicted probability of a bug’s flight case as well as the odds (OR) ratio between flight cases. The OR signifies the ratio of the probability that a flight case will happen compared to the probability that another flight case (i.e. the ‘baseline’) will happen. Refer to Appendix B section 3.5 to see how the OR is calculated from the prediction equations of a multinomial logistic model. Finally, we used each best fit model to predict flight case probabilities for bugs flight tested in Fall 2019 in order to assess the accuracy and overall performance of each best fit model. See Appendix B for a tidy version of the analyses.

## 3 Results

### 3.1 Wing Morphology of Field Bugs Over Time

#### 3.1.1 Long Wing Morph Frequency

The best fit model for wing morph included sex, host plant, month, and year as single factors and interactions (*n* = 3532; Appendix A section 3.1.1). Male soapberry bugs were 30% more likely to be long-winged than females (*β* = −0.26, *SE* = 0.09, *P* = 0.004). Soapberry bugs collected from *K. elegans* were 208% more likely to be long-winged than those collected from *C. corindum* (*β* = 1.13, *SE* = 0.11, *P* <<0.05). Within a given year, long wing morphs were approximately 11% more likely to appear for each month passed (*β* = 0.10, *SE* = 0.03, *P* << 0.05; Figure 3A, B). Long wing morph frequency also increased by 1% for each year passed, but this was a relatively weak effect (*β* = 0.01, *SE* = 0.003, P <<0.05; Figure 3C). Yearly and monthly interactions were also relatively weak but nuanced. Long wing morphs were increasingly more likely to appear in later months than earlier months before 2019 but less likely to appear in later months than earlier months after 2019 (*β* = −0.002, *SE* = 0.0005, *P* = 0.001; Appendix A section 3.1.1)

**Figure 3:**
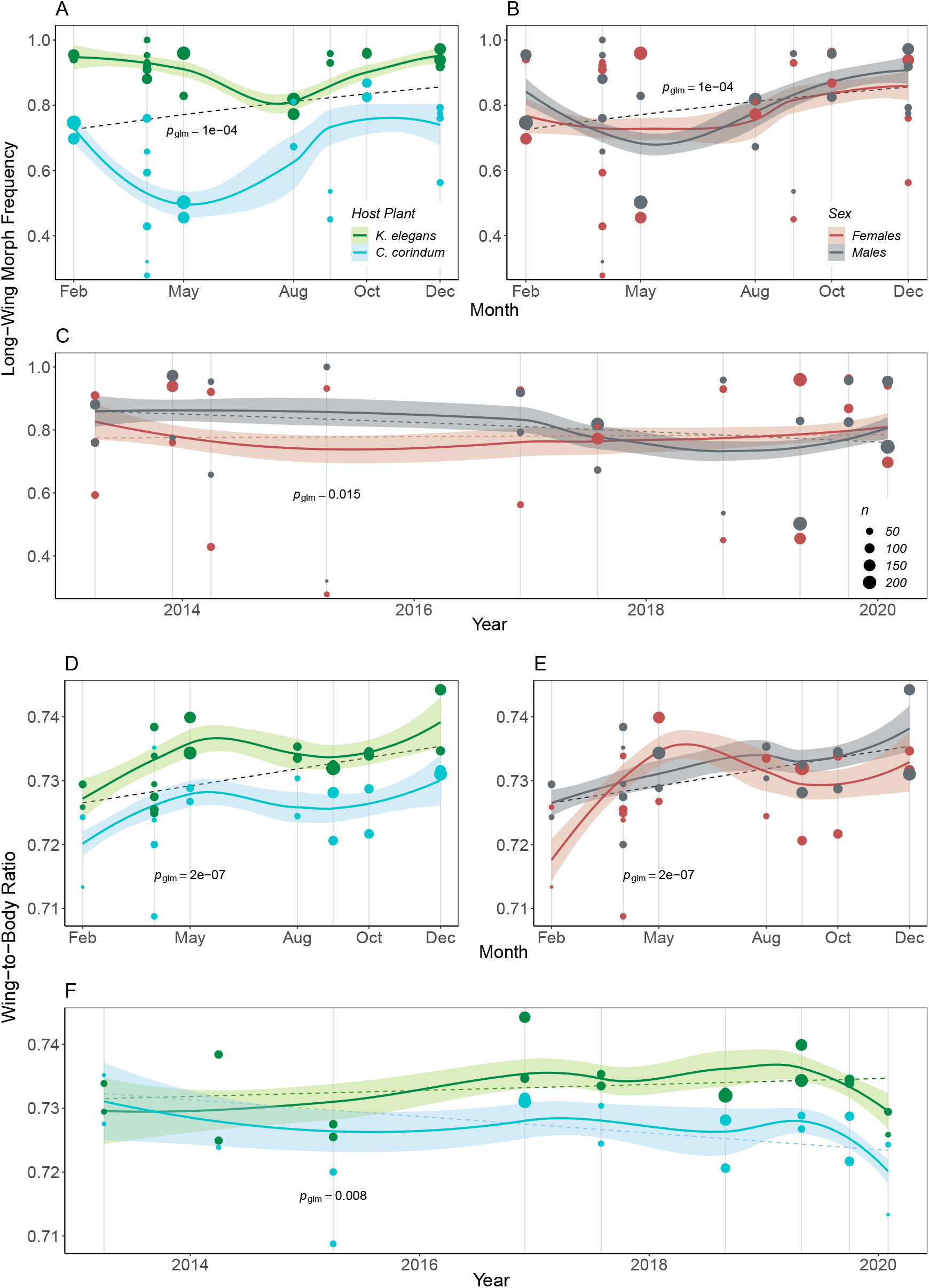
Evaluation of long-wing morph frequency and wing-to-body ratio of soapberry bugs averaged across month and year from April 2013 to February 2020. For each point, the mean long-wing morph frequency or wing-to-body ratio of each month and year is plotted with LOESS smooth lines (solid lines) and 95% confidence intervals (shading) and linear regression line(s) (dashed line(s)). LOESS plots are only for visual purposes. For LOESS regressions, the smoothing parameter (*α*) was 0.4 and the degree (*λ*) was 0. For linear regressions, each P-value was extracted from their corresponding best fit model [**A-C** formula=wing morph ~ sex * host + sex * year + host * month + month * year or **D-F** formula=wing2body ~ sex * host + host * year + month]. **A, D, F**. Each point, line, or shading colored green corresponds to soapberry bugs collected from *K. elegans*, while those colored blue correspond to soapberry bugs collected from *C. corindum*. **B, C, E**. Each point, line, or shading colored black corresponds to male soapberry bugs while those colored red correspond to female soapberry bugs.

The best fit model included three additional interaction terms, as follows: sex and host plant, sex and year, host plant and month. Female soapberry bugs from *K. elegans* or males from *C. corindum* were 4.2% more likely to be long-winged compared to their host plant counterparts within their sex group (*β* = 0.10, *SE* = 0.05, *P* = 0.05; Appendix A section 3.1.1). Female soapberry bugs also were 0.4% more likely to be long-winged each proceeding year since the first collection unlike their male counterparts which are becoming less likely to be long-winged. However, the effect of this interaction is relatively small (*β* = 0.004, *SE* = 0.002, *P* = 0.02; Figure 3C). Finally, soapberry bugs collected from *C. corindum* were 3.7% more likely to be long-winged for each month passed, whereas individuals collected from *K. elegans* decreased by 3.9% each month (Appendix A section 3.1.1 & 4.1.3).

#### 3.1.2 Wing-to-Body Ratio

Bugs that were short-winged or had torn wings were then filtered from the dataset. From this subset of the data, the best fit model for wing-to-body ratio similarly included sex, host plant, month, and year as single factors and interactions (*n* = 1903; Appendix A section 3.3.1). Males had larger wing-to-body ratios (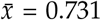, *SE* = 6e-04) than females (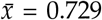, *SE* = 7e-04) by 0.002 units, on average (*β* = −0.002, *SE* = 0.0005, P <<0.05; Figure 3E). Soapberry bugs collected from *K. elegans* had larger wing-to-body ratios by approximately 0.004 units, on average, than those collected from *C. corindum* (*β* = 0.004, *SE* = 0.0005, *P* <<0.05; Figure 3). However, there was a significant interaction between sex and host plant, with female soapberry bugs collected from *K. elegans* having the larger wing-to-body ratios, on average, than their male or *C. corindum* counterparts (*β* = 0.002, *SE* = 0.0005, *P* <<0.05; Appendix A section 3.3.1).

There was no effect of year (*P* = 0.43), but there was a significant year and host plant interaction where soapberry bugs collected from *K. elegans* have larger wing-to-body ratios each year since 2013 by 6e-05 units, on average (*β* = 6e-05, *SE* = 2e-05, *P* = 0.007; Figure 3F; Appendix A section 3.3.1). Larger wing-to-body ratios were also increasingly likely later in the year (*β* = 0.0007, *SE* = 0.0001, *P* << 0.05) where each month wing-to-body ratios increased by 0.0007 units, on average (Figure 3D, E).

### 3.2 Winter 2020 Across-Trial and Between-Trial Flight Response

#### 3.2.1 Across Trials

One factor from our experimental design affected flight response across trials: days since the start of trials. On average, soapberry bugs tested on day 20 compared to day 1 of trials had a 15% drop in flight probability (*β* = −0.008, *SE* = 0.002, *P* = 0.017; Appendix B section 2.4), suggesting a possible age affect [xxxx]. Individuals were randomized across date and start times; however, to control for the impact of days since the start of trials on flight response, we calculated and used each individual’s mean trial date in our models (Appendix B section 2.5.1).

The best fit model for flight response across trials included sex as a significant single factor as well as significant interactions of mass with host plant and mass with latitudinal distance from the sympatric zone (n = 333; Appendix B section 2.5.3). Male soapberry bugs were 59% more likely to fly than females (*β* = −0.46, *SE* = 0.17, *P* = 0.006). Soapberry bugs from *K. elegans*, which resides in the mainland of Florida, were more likely to fly if they were heavier; individuals 0.05 g heavier than average were 50% more likely to fly than their lighter counterparts (*β* = 1.86, *SE* = 0.59, *P* = 0.002; Figure 4A). The opposite was true for soapberry bugs from *C. corindum*, which dominates the islands of Florida; here, bugs that were 0.05 g heavier than average were 31% less likely to fly than their lighter counterparts (Figure 4B). Additionally, the effect of mass depended on latitudinal distance from the sympatric zone: for bugs two latitudinal degrees further, individuals were 25% less likely to fly than bugs one degree closer (*β* = −1.41, *SE* = 0.69, *P* = 0.04; Figure 4 and Appendix B section 2.5.3).

**Figure 4:**
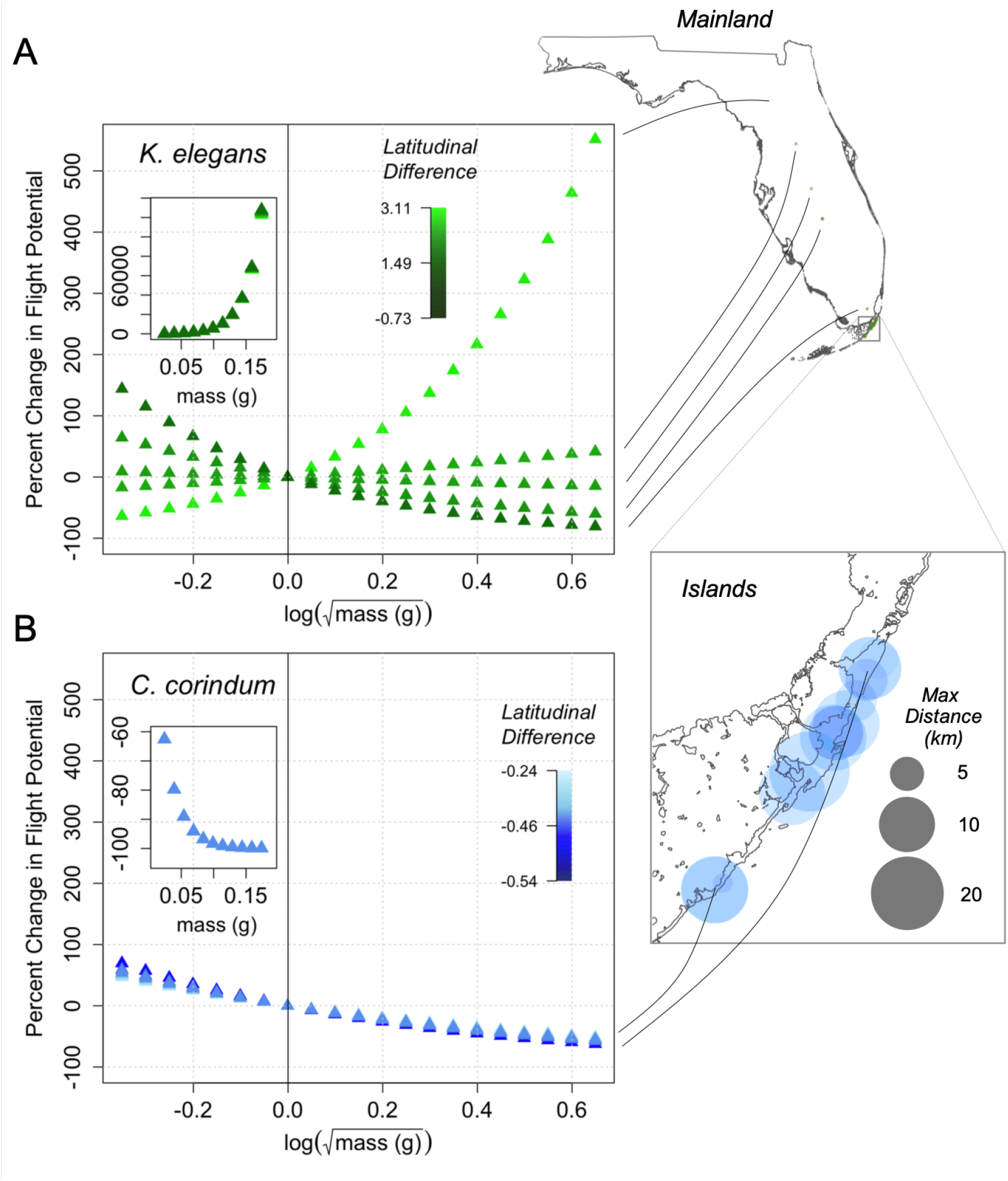
Estimated percent changes of flight potential as a function of soapberry bug mass averaged across trials. Color gradients in the plots represent latitudinal differences from the sympatric zone (Homestead, FL; latitudinal distance = −0.73), where lighter shades indicate a location far from Homestead and darker shades indicate a location close to Homestead. Latitudinal differences are standardized. Percent mass changes in flight response and the average mass for tested soapberry bugs (*n* = 333) were fitted using multiple regression modeling. From the best fit model, selected interaction effects were extracted [formula=flight potential ~ latitudinal difference * mass + host * mass].

Both the transformed and back transformed (top-left, smaller graph) coefficients are plotted. The maps of Florida on the right-hand side display the latitudinal locations of most sites and their corresponding fitted line. The map of the islands of Florida emphasizes the maximum flight distance (to scale) reached by a soapberry bug collected from each island site, which helps visualize the bugs’ ability to traverse between the island and mainland. **A** Only soapberry bugs collected from *K. elegans* are plotted in green triangles, showing relatively more variation in flight potential between collection sites as well as more variable relationships between flight potential and mass between collection sites. **B** Only soapberry bugs collected from *C. corindum* are plotted in blue triangles, showing little variation in flight potential between collection sites.

When models were separated by sex, we found wing-to-body ratio primarily drove male flight and mass drove female flight. The best fit multiple regression model for females included average mass, host plant, wing-to-body ratio, and average days since the start of trials (*n* = 120; Appendix B section 2.5.4). We found that a female tested a day later on average would be 12% more likely to fly (*β* = 0.12, *SE* = 0.05, *P* = 0.015), but later testing dates could be masking changes in female egg laying throughout the trials. We also found that, for every 0.05 unit increase in wing-to-body ratio, females experienced a 31% increase in flight potential (*β* = −5.37, *SE* = 2.66, *P* = 0.044). As with the full dataset, we found a mass by host plant interaction: a female that was 0.06 g heavier than average and from *K. elegans* would be 83% more likely to fly than those from *C. corindum (β* = 3.02, *SE* = 1.40, *P* = 0.031).

The best fit multiple regression model for males included wing-to-body ratio, host plant, and distance from the sympatric zone (*n* = 213; Appendix B section 2.5.4), but notably did not include mass. For males, every 0.05 unit increase in wing-to-body ratio corresponded to a 53% increase in flight potential (a higher rate than females; *β* = −15.21, *SE* = 5.22, *P* = 0.004). Additionally, wing-to-body ratio interacted with host plant: a 0.05 unit longer-than-average winged male from *C. corindum* would be 38% more likely to fly than those from *K. elegans* (*β* = −9.46, *SE* = 4.28, *P* = 0.027) and 37% more likely to fly if it was 1 latitudinal degree farther from the sympatric zone (*β* = 6.31, *SE* = 3.03, *P* = 0.037).

#### 3.2.2 Between Trials

The best fit multinomial logit model used to explain a soapberry bug’s observed flight case included mass percent change, sex, and wing-to-body ratio. Host plant was not significant. Results of the multinomial regression analysis are presented in Table 2; also, see Appendix B sections 3.5.4 and 3.5.5 for each prediction equation and for a heatmap adaptation of the results.

**Table 2:** Results of multinomial logistic (ML) regression for predicting flight case for all soapberry bugs. The odds of a soapberry bug exhibiting a particular flight case rather than another flight case (i.e. the base category) are presented below each column. T1 denotes trial 1 and T2 denotes trial 2. Statistically significant results (*p* < 0.05) are in boldface and 95% confidence intervals are adjacent to the odds. Only ML prediction equations with at least one significant main effect are shown; however, ML equations comparing whether a soapberry bug flew in T2 to another flight case are not shown despite sex being significant. This is because there were no flight cases where females only flew in T2 during Winter 2020 trials.

| Variables in the Model | Flew twice rather than flew in T1 | Flew twice rather than did not fly | Flew in T1 rather than did not fly |
| --- | --- | --- | --- |
| Mass % Change * | **0.67 (0.66, 0.68)** | **1.49 (1.46, 1.52)** | **2.23 (2.18, 2.27)** |
| Sex (female = 1) | 0.83 (0.56, 1.22) | **0.47 (0.34, 0.65)** | **0.57 (0.37, 0.85)** |
| (male = -1) | 1.21 (0.82, 1.79) | **2.14 (1.53, 2.98)** | **1.77 (1.17, 2.67)** |
| Wing-to-Body * | 1.04 (0.84, 1.29) | **1.32 (1.09, 1.6)** | **1.27 (1, 1.61)** |

\* The mass percent change estimates and wing-to-body estimates were multiplied by 20 and 0.01, respectively, before calculating the odds. These transformations better represent experimental observations and offer a more realistic odds (e.g. 20% mass increase and a 0.01 unit increase in wing-to-body ratio).

We found that if a soapberry bug had a 20% increase in mass, the odds of flying in T1 rather than twice (*β* = 0.02, *SE* = 0.01, *P* = 0.032) as well as the odds of flying twice rather than not flying (*β* = 0.04, *SE* = 0.01, *P* << 0.05) were both 30% lower compared to a bug that flies only in T1 rather than not flying (*β* = −0.02, *SE* = 0.01, *P* = 0.005). In other words, between flying twice, flying once in T1, and not flying at all, a bug had the highest odds of flying in T1 when a bug gained 20% of its original mass.

Sex and wing-to-body ratio were also significant for prediction equations where a bug could either fly once or twice rather than not fly at all. For bugs that had a wing-to-body ratio 0.01 units higher than average, the odds of flying twice (*β*= 0.28, SE = 0.10, *P* = 0.004) were 4% higher compared to the odds of flying in T1 (*β* = 0.24, *SE* = 0.12, *P* = 0.049). Males had a 78% higher odds of flying twice (*β* = −0.76, *SE* = 0.17, *P* << 0.05) and a 68% higher odds of flying in T1 only (*β* = −0.57, *SE* = 0.21, *P* = 0.007) compared to females. Likewise, the odds of flying twice compared to flying in T1 was 17% higher for males but approximately 17% lower for females. In summary, females not only had a lower odds of flying twice compared to males, but females also had the highest odds of not flying at all whereas males had the highest odds of flying twice.

Predicted probabilities showing the likelihood of a soapberry bug exhibiting one flight case over another can be seen in Figure 5. Figure 5B depicts males’ propensity to fly repeatedly and their narrower mass range compared to females’ larger mass ranges and propensity to fly either only in T1 or not at all. Figures 5A and 5B also display the stochasticity in flight case probability introduced by wing-to-body ratio where, across the sexes, larger wing-to-body ratios led to greater probabilities that bugs would fly twice or fly in T1 rather than not fly at all. Additionally, probability thresholds can be extracted from Figure 5. For example, if a female were to gain more than 41% of her original body mass, then she would be most likely to fly in T1. Such results reinforce previous analyses that female flight potential is consistently more dependent on mass, whereas male flight potential is strongly influenced by wing morphology.

**Figure 5:**
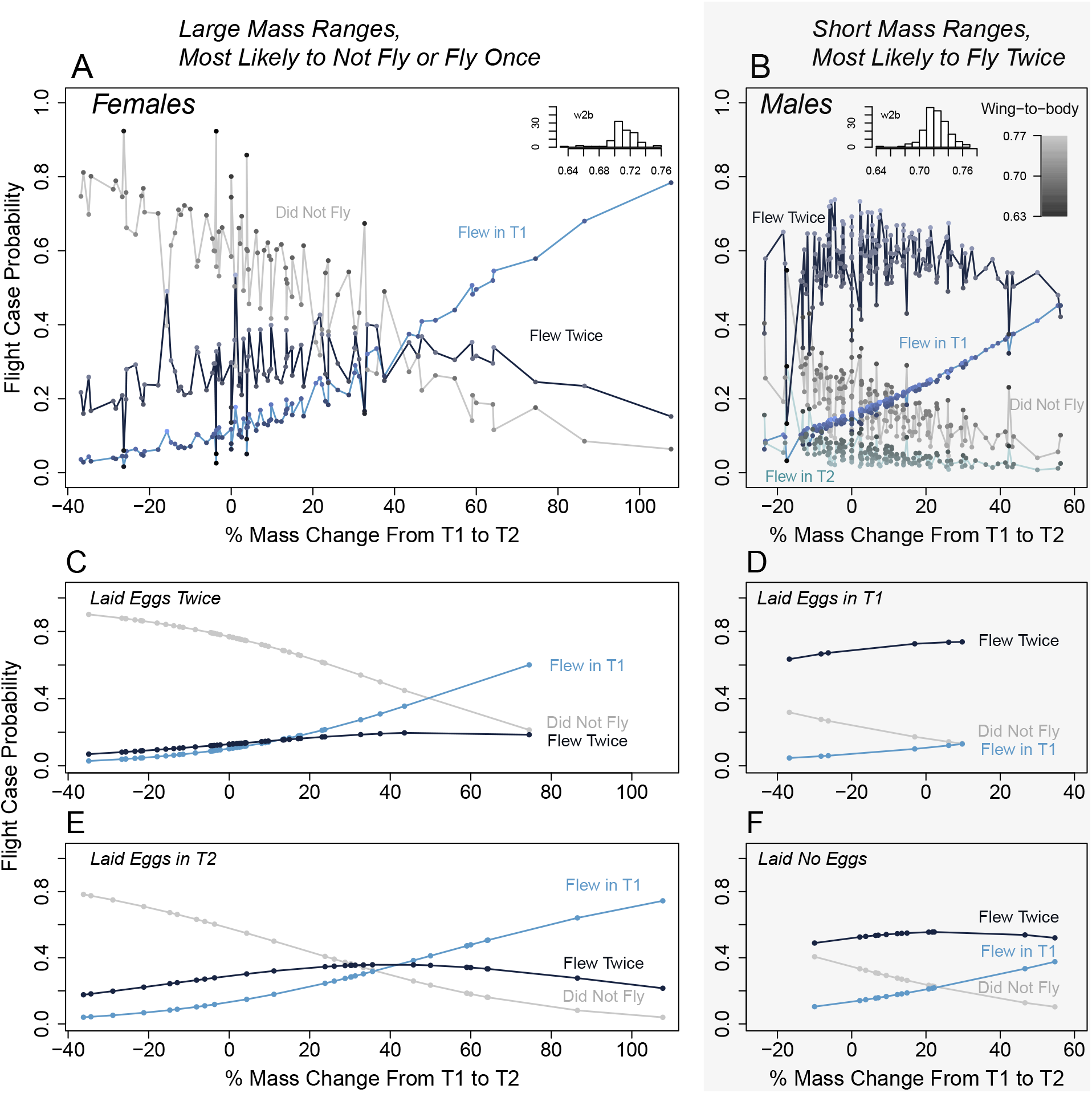
Estimated probabilities of different flight cases as a function of soapberry bug percent mass change between trial 1 (T1) and trial 2 (T2). The flight case probabilities were fitted using multinomial logit modeling. Solid lines represent estimated probabilities and each line color represents a flight case. Plot shading highlights similarities between short or large mass ranges across sexes or reproductive status. The flight case probabilities of **A**. females (*n* = 93) and **B**. males (*n* = 185) are individually plotted with the histogram of wing-to-body ratio (w2b) in the top-right corner of each plot. Each sex shares the same best fit multinomial logit model [formula=flight case ~ sex + wing-to-body ratio + mass percent change between trials]. The stochasticity in the plots represent the affect wing-to-body ratio has on flight case probability where lighter shaded circles represent larger wing-to-body ratios and darker shaded circles represent smaller wing-to-body ratios. **C-F**. Estimated probabilities of different flight cases plotted for only female soapberry bugs (n = 93) and their corresponding best fit model [formula=flight case ~ egg case + mass percent change between trials]. Each plot represents a different egg laying case for a female soapberry bug where **C**. (*n* = 45), **D**. (*n* = 6), **E**. (*n* = 28), and **F**. (*n* = 14).

For females, the timing of egg laying also impacted certain flight case outcomes. In the best fit multinomial logit model for females only, predictor variables included mass percent change and egg case. Results of the multinomial regression analysis are presented in Table 3; also, see Appendix B section 3.6.5 and 3.6.6 for each prediction equation and for a heatmap adaptation of the results.

**Table 3:** Results of multinomial logistic (ML) regression for predicting flight case for female soapberry bugs. The odds of a soapberry bug exhibiting a particular flight case rather than another flight case (i.e. the base category) are presented below each column. T1 denotes trial 1. Statistically significant results (*p* <0.05) are in boldface and 95% confidence intervals are adjacent to the odds. Only ML prediction equations with at least one significant main effect are shown.

| Variables in the Model | Flew twice rather than did not fly | Flew in T1 rather than did not fly |
| --- | --- | --- |
| Mass % Change * | **1.49 (1.46, 1.52)** | **2.23 (2.18, 2.27)** |
| Egg Case (laid eggs twice = 2) | **0.11 (0.06, 0.2)** | 0.35 (0.16, 0.73) |
| (laid eggs in T2 = 1) | **0.33 (0.18, 0.6)** | 0.59 (0.28, 1.24) |
| (did not lay eggs = 0) | **1 (0.56, 1.8)** | 1 (0.47, 2.11) |
| (laid eggs in T1 = -1) | **3 (1.67, 5.41)** | 1.7 (0.81, 3.58) |

\* The mass percent change estimates were multiplied by 20 before calculating the odds. This transformation better represents experimental observations and offers a more realistic odds (e.g. 20% mass increase).

We found that if a female had a 20% increase in mass and she had the chance of flying either once or twice rather than not flying at all, she was 33% less likely to fly twice (*β* = 0.02, *SE* = 0.01, *P* = 0.04) compared to the odds of flying only in T1 (*β* = 0.04, *SE* = 0.01, *P* = 0.001). Additionally, the odds of flying twice were 67% lower for a female that had laid no eggs, 89% lower for a female that laid eggs only in T2, and 96% lower for a female that laid eggs twice compared with those that had only laid eggs in T1 (*β* = −1.10, *SE* = 0.30, *P* << 0.05). It appears that for females who have completed oviposition or are not close to oviposition, they will most likely fly twice, irrespective of changes in mass (Figure 5D and 5F); their flight behavior also mimics the flight behavior and mass changes of males (Figure 5B). For females who are in the middle of oviposition or nearing the beginning of oviposition, they appear to be most likely to not fly if not enough mass is gained or mass is lost (mass range: −30%, ~30%; Figure 5C and 5E). However, if enough mass is gained (mass range: 40%, 110%), females have higher chances of at least flying once Figure 5C. Refer to Appendix B section 3.6.7 for an alternative display and reading of Figure 5.

### 3.3 Predicting Fall 2019 Flight Potential

To test the performance of our models, we calculated flight case accuracy for the Fall 2019 flight tested bugs (Appendix B Section 4). For the best fit model for all Winter 2020 soapberry bugs, the overall prediction accuracy was 0.60 (*n* =45), with female prediction and male prediction accuracy at 0.38 and 0.69, respectively. Because the model favored repeated flight events and not all male bugs always flew twice, this model over-estimates flight capability. It over predicts when bugs fly twice (73.3%) and misses events when they only fly once (0%). For the best fit model for Winter 2020 female bugs, the overall prediction accuracy was 0.46 (*n* = 13). Females in the Fall mostly laid eggs twice (*n* = 10), flew twice, and had a narrower mass range than females tested in the Winter (mass range: −20%, 30%; Appendix B section 3.6.7). The model greatly overestimates flight capability by overpredicting that female will only fly twice (100%) even though females in the Fall flew twice approximately half of the time (46.2%). This suggests that the narrower mass changes could be advantageous for females to fly repeatedly despite being in the middle of oviposition.

## 4 Discussion

In this study, we examined patterns of dispersal behavior that demonstrated plastic flight responses as well as recent trends in wing morphology in soapberry bugs from Florida, USA. We found plastic vs. fixed dispersal behavior to be sex-biased and wing morph-biased. We also found a latitudinal cline and host plant differences in flight response and demonstrated plastic flight responses largely mediated by mass. In particular, we identified macropterous, island-native males as the dispersal unit of this soapberry bug species as well as identified its complementing plastic dispersal unit: macropterous, mainland-native females. Furthermore, we modeled mobility characteristics that reveal links between a population’s evolutionary history and its environmentally-induced phenotypic variation [28,60]. Finally, given how plasticity impacts the ability of animals not only to adapt but physically maneuver through degraded habitats or invasive species [27,61], we provide a step towards empirically predicting mobility for organisms undergoing HIREC.

### 4.1 Sex Differences in Flight Potential

Our results unequivocally showed sex differences in dispersal-behavioral plasticity. Males were consistent flyers and their flight response was positively driven by their wing-to-body ratio. They invested more in flight morphology and they were more likely to fly and fly again irrespective of changes in mass. In contrast, females were more variable, mass-dependent flyers because of trade-offs with egg production. Male-biased dispersal and trade-offs between investments in flight structures and reproduction have been demonstrated in various wing-polymorphic species [54, 62]. For instance, short-wing morphs of the cricket *Gryllus rubens* more efficiently assimilate nutrients into ovarian mass and overall biomass, compared to their long-winged counterparts who invest in flight muscle mass [63]. Bush crickets, as measured via microsatellite DNA analyses, also largely favored long-winged males as the dispersal unit of their species, which were more frequently found at their range margins compared to longer-established populations in their range core [54]. Our observations that males are more dispersive and exhibit more fixed, as opposed to plastic, dispersal behavior suggest males similarly can be held accountable for the range expansion of their species. In turn, metapopulation connectivity, colonization, and spatial sorting [49] are likely to be driven primarily by males.

Female flight potential, on the other hand, was strongly associated with mass changes, host plant, and reproductive state. Generally, female soapberry bugs exhibited a negative mass effect that differed by host; heavy females from the invasive host were far more likely to fly than those from the native host. However, reproductive state could reverse negative mass relationships, making females more closely resemble their male counterparts (Figure 5). Females that were at the end of oviposition were most likely to fly repeatedly and they sustained short mass changes. Conversely, females in the middle or beginning oviposition had large mass ranges and were either prone to fly once or not fly at all.

These findings raise one possible interpretation of female dispersal. Female flight response, evidently plastic, could be characterized as more strategic and time-sensitive as females coordinate their flight behavior based on their reproductive activity. Time-sensitive dispersal behavior has been seen for monarch butterflies in Texas and northern Mexico that consistently increase their lipid levels before Fall migrations [64]; however, none to our knowledge have connected insect flight behavior to mass and egg laying changes observed during a two week span of testing. This observation demonstrates that brief trade-offs between dispersal and egg-laying would demand optimizing the right time to be relatively mass stable in order make a strong one-time flight attempt before laying eggs. Likewise, changes in mass – a fluctuating metric throughout an insect’s life – could temporarily induce trade-offs with dispersal but also provide the necessary physiological advantage before takeoff.

From these sex-biased plastic dispersal patterns, we can favorably characterize males as the dispersal unit of this soapberry bug species and females as the plastic dispersal unit. This characterization can relate to how soapberry bugs traverse and interact with their environment. We found that males and females raised on on *C. corindum* and males collected from the southern end of the Floridian islands had higher tendencies to fly. With the Atlantic Ocean acting as a spatial barrier, island-native males or females who have ended or not begun oviposit would favor flying towards the mainland or northern islands, taking advantage of range edges as a means to drive phenotypic change where “space per se is an evolutionary agent” [49]. Likewise, females who leave their host to oviposit on the invasive host can facilitate rapid directional gene flow from *C. corindum* to *K. elegans* [51] or, in theory, they can induce rapid gene flow reversals [65], making populations between the islands and mainland more insular in response to HIREC.

However, it is uncertain whether fast-dispersing soapberry bugs at the edges of their dispersing fronts will breed and lead to successive generations that evolve to be even faster than the last. It is also unclear whether spatial sorting or adaptive radiation leads to ecological opportunities [66], especially under a rapidly changing environment. This is relevant considering that our Winter 2020 model’s ability to predict Fall 2019 female response was always below 50% as opposed to 69% for male flight response. Thus, sex-biased evolutionary-dispersal mechanisms behind soapberry bug gene flow warrant further study accompanied by a phylogeographic analysis [67,68] or observations on additional traits (e.g. age, thorax muscle mass, thorax muscle histolysis, or diapausing status) [47,69].

### 4.2 Host Plant Differences and Latitudinal Cline in Flight Potential

To then better understand how host plant or latitudinal distance from the sympatric zone affected soapberry bug dispersal, we included environmental factors in our models and found another case of dispersal-behavioral plasticity dependent on mass. Specifically, dispersal was more dependent on mass and latitudinal distance for mainland-native bugs collected from *K. elegans* than for island-native bugs from *C. corindum*. The deeper a bug from *K. elegans* was into the mainland, then the more likely it will fly if it was heaver but if it was closer to the islands, then the more likely it will fly if it was lighter. Such a relationship was substantially variable compared to island-native bugs collected from *C. corindum* where, instead, heavier bugs would be less likely to fly, regardless of where it was located.

Previous studies have suggested that, in locations where dispersal between the divergent environments is the highest, local adaptation might not be apparent because of high migration and gene flow [30, 70]. In response, plasticity can evolve to allow these populations to display locally adaptive phenotypes in the absence of genetic differentiation. This was observed in a study tracking brain mass variability and dispersal potential in African cichlid fish in the oxygen-diverse rivers, lakes, and swamps of East Africa [30]. However, we observe a more complex scenario. Even though heavy soapberry bugs from the mainland exhibited the highest flight potentials, low-weight bugs from the islands still consistently exhibited high flight potentials; and yet, only mainland bugs exhibited a plastic latitudinal cline. This discrepancy could be due to whether soapberry bugs across Florida deferentially recognize adaptive vs. maladaptive responses to rapid shifts in host plant utilization. In this case, maladaptive responses would suggest soapberry bugs are dampening their abilities to disperse for resources and mates; highly variable flight could then be less of a phenotypic display and more of an evolutionary trap [12] because mainland bugs don’t readily recognize the potential for island resources.

Additionally, given that the ability to fly is an act that is highly mechanistic, metabolic, and spatial [55,56,57,58,59], it may be that the energetics and physics of dispersal cannot be readily divorced from the larger narrative of plastic potential and its costs. As such, a narrower, more fixed range of flight potential in the islands could be necessary if the costs of failing to traverse to the mainland, which results in death, outweighs the benefit of being plastic flyers. Meanwhile, mainland-native bugs without such irreversible costs may show less consistent investments in flight performance. To better evaluate the costs of dispersal-behavioral plasticity, it then becomes important to measure additional insect flight performance metrics (e.g. distance or duration) [unpublished data collected by AVB and MLC], especially for insects undergoing HIREC who may adapt or sink in response to human influences in unexpected ways.

### 4.3 Wing Morphology Through Time

Our analyses of soapberry bug wing morph and wing-to-body ratio from 2013 to 2020 identified annual and seasonal trends in soapberry bug wing morphology. Such trends show how dispersal could change in the future, as this insect-host system continues to undergo evolutionary responses to HIREC. First, we found strong, positive within-year trends in long-wing morph frequency and wing-to-body ratio. Second, we found that males, which significantly depend on larger wing-to-body ratios for higher flight potentials, are gradually becoming more short-winged over time. Likewise, soapberry bugs collected from the native host are gradually exhibiting smaller wing-to-body ratios. Even though males are consistent flyers and would most likely be the first in a population to traverse between the islands and mainland, these trends suggest that the traits favoring frequent flight are being selected against. Possibly, this pattern implicates another case of maladaptive plasticity induced by host plant on the system [50] where flightlessness is impeding some dispersal agents from traversing between the mainland and islands for resources and mates. This could lead to narrower windows of high dispersal for males and for soapberry bugs from *C. corindum*. Conversely, it would appear that female and mainland dispersal agents are either generally unaffected or actively resisting evolutionary responses to HIREC in the last decade due to their plastic dispersal behaviors; specifically, their plasticity may be dampening natural selection for less flight-capable traits [71]. As similarly observed by studies on birds, high phenotypic plasticity, behavioral flexibility, and/or greater within-species variation in behavioral tendencies allow a species to cope well with environmental variation [72] and extinction [26], and the same could be occurring in Floridian soapberry bugs.

### 4.4 Conclusions

Dispersal-behavioral plasticity determines the ability of animals to physically maneuver and cope with new or unusual challenges such as rapid host invasions or extreme climate events. Dispersal-behavioral plasticity thus has the potential to reduce species vulnerability and enhance population fitness following environmental changes. However, in light of the speed and force of human culture evolution, we have put into question how biota will weather and response to these rapid changes, how does dispersal link plasticity to HIREC, and how we can better predict who will adapt or maladapt? Additionally, as ethology reconciles losing the contexts in which organisms originally evolved in, it may become increasingly necessary to not speak too broadly about environmental changes but instead become abundantly more specific about an organisms’ evolutionary history, especially if it has been significantly mediated by human activities. This specialization may seem counterintuitive to a discipline defined by broad patterns in behavioral ecology, and it may be tempting to advise that studies like ours simply be assimilated into ethology; however, it may be worthwhile to instead consider the evolutionary force of humans as imperatively self-identifying because of its scope. To consolidate this, empirical studies would need to continue to quantify and demonstrate the links and patterns formed between plasticity and HIREC as well as invest in tracking direct phenotypic plastic changes over time.

## 6 Acknowledgements

We would like to thank Ana Silberg for her assistance in taking flight trial measurements and John Zdenek for overseeing incubator setup and maintenance at the University of Chicago. We also thank Micah Freedman for helpful and detailed comments on various manuscript drafts. Finally, we thank the Florida Division of Recreation and Parks for permission to collect Jadera haematoloma and Cardiospermum corindum seeds within Florida state parks. All funding was provided by the Department of Ecology & Evolution at the University of Chicago.

## 7 Author Contributions

MLC conceived and designed the study; AVB enhanced the design of the study, performed the data collection of flight trials, and assisted the data collection of morphology; MLC performed the data collection of morphology and assisted the data collection of flight trials; AVB and MLC drafted code; AVB analyzed the data and MLC code reviewed; AVB drew or generated the figures; AVB wrote the paper and appendices. All authors contributed to the final draft.

## 9 Data Availability Statement

See Dryad (https://doi.org/10.5061/dryad.XXXX.) for a full description of each experimental setup and its accompanying data.

