## Appendix A for "Dispersal-behavioral plasticity within an insect-host system undergoing human-induced rapid environmental change (HIREC)"

Bernat, AV, Cenzer, ML

### 1 Details of the Analyses

This document was generated by R Markdown on 2022-06-19 using R version 4.0.5 (2021-03-31). The document provides the step-by-step analytical methods used in the manuscript by Anastasia Bernat (AVB) and Meredith Cenzer (MLC). Multiple draft scripts were written by AVB between 2021-03-01 and 2021-07-26 until being distilled and complied by AVB and code reviewed by MLC at the University of Chicago into this comprehensive script. All draft scripts can be viewed in the GitHub repository, SBB-dispersal (https://github.com/mlcenzer/SBB-dispersal), within the directory **avbernat > All_Morphology > stats**.

All code and output from the statistical analyses are shown. Code for data cleaning and the generation of plots is not displayed but can be viewed in the **appendix_A-wing_summary.Rmd** file and its accompanying sourced scripts. To repeat analyses and the generation of plots, all data files and sourced scripts should follow the directory structure presented in the SBB-dispersal repository.

#### 1.1 Description of the Data

This document analyzes two main datasets, `raw_data` and `data_long`. The raw_data set provides morphology measurements for each soapberry bug, *Jadera haematoloma*, collected and measured between the April 2013 and February 2020. There are four morphology measurements: beak length, thorax width, wing length, and body length. The sex, wing morph (long-winged, shot-winged, or ambiguously-winged), and host plant the bug was collected from as well as the month and year each bug was collected in was recorded. The data_long set provides the same recordings as the raw_data set, but it has been filtered for only long-winged soapberry bugs.

#### 1.2 Abbreviations Used in the Data and Code

- **SBB** - soapberry bug, *Jadera haematoloma*
- **S** - short-winged morph
- **L** - long-winged morph
- **LS** or **SL** - ambiguous wing morph
- **pophost** - the host plant soapberry bugs were collected from, which was either *Koelreuteria elegans* or *Cardiospermum corindum*, occasionally called (and abbreviated) as goldenrain tree (GRT) or balloon vine (BV), respectively
- **months_since_start** - proxy for year where the first collection occurred on April 2013
- **month_of_year** - proxy for season where collections occurred only in months February, April, May, August, September, and October
- **wing2body** - a computed and unitless value calculated from the wing length divided by the body length of a soapberry bug
- **wing2thorax** - a computed and unitless value calculated from the wing length divided by the thorax width of a soapberry bug
- **sd** - standard deviation
- **se** - standard error
- `w_` - a column name that starts with `w_` is shortened from "wing" (e.g. w_morph is "wing morph")
- `_c` - a column name that ends in `_c` is a column that has been centered. Example columns: wing2body_c, month_of_year_c, and months_since_start_c
- `_b` - a column name that ends in `_b` is a column that has been recodified into binary data (0's and 1's). Example columns: sex_b, pophost_b, and wing_morph_b

### 2 Data Cleaning And Exploration

#### 2.1 Read Libraries

The occurrence of long-wing morphology and the wing-to-body ratio of *J. haematoloma* were analyzed using multivariate, generalized linear modeling (GLM) as implemented in the R packages `lme4` and binom. The dplyr package helped pipeline data manipulation processes by grouping data quickly. All plots, except the histograms, were generated using ggplot libraries and helper functions found in R packages ggformula and cowplot.

Additional R packages not shown below, but embedded in the sourced scripts are zoo and lubridate, which aid in data manipulation and datetime manipulation, respectively.

```
library(lme4)       # fit regressions
library(dplyr)      # data manipulation
library(ggformula) # ggplot plotting
library(cowplot)    # ggplot helper functions to arrange multi-panel figures
library(binom)      # binomial confidence intervals
```

#### 2.2 Read Source Files

Each sourced script below aides in either data cleaning (read_morph_data(), remove_torn_wings()) or multivariate GLM (model_comparisonsAIC(), get_model_probs()). Additionally, the function model_comparisonsAIC() takes in the path of a generic multi-factor script with a specified, hard-coded GLM family and link function needed to build the predictive models. All aforementioned sourced scripts are located in the **Rscr** folder.

```
source_path = paste0(dir,"/Rsrc/")

script_names = c("clean_morph_data.R",    # 1 function: read_morph_data()
                 "remove_torn_wings.R",  # 1 function: remove_torn_wings()
                 "compare_models.R",     # 1 function: model_comparisonsAIC()
                 "get_Akaike_weights.R") # 1 function: get_model_probs()

for (script in script_names) {
  path = paste0(source_path, script)
  source(path)
}
```

#### 2.3 Read the Data

The morphology data were started in 2013-04-28 and last updated on 2021-05-18. The read_morph_data() function standardizes population names, host plant names, and month and year inputs. Month and year inputs are also converted into datetimes. Variables of interest like wing-to-body ratio and wing-to-thorax ratio are also calculated and centered. The full dataset, raw_data (n=3532), and a long-winged bug dataset, data_long (n=2096), are returned.

```
datapath = paste0(dir,"All_Morphology/stats/data/allmorphology05.18.21.csv")
data_list = read_morph_data(datapath)
```

```
## morph types: L S  NA LS SL
##    recoding missing morph types...
##    S if wing2thorax <=2.2, L if wing2thorax >=2.5
##
## ambiguous wing morph bug count:  48
```

```
raw_data = data_list[[1]]
data_long = data_list[[2]] # long-wing bugs with wing-to-body value only

data_long = remove_torn_wings(data_long)
```

```
##
## number of bugs with torn wings: 193
```

Bugs marked as having torn wings during measurements were only filtered out of the `data_long` dataset (n=1903). That was because `data_long` is used only to analyze the wing-to-body ratio, which was computed for long-winged bugs since no short-winged bugs can fly. `raw_data` is only used to analyze long-wing morph frequency.

#### 2.4 Histograms of Wing Morph Data

Soapberry bugs have two noticeable wing morphs: a long-wing (left) and short-wing morph (right).

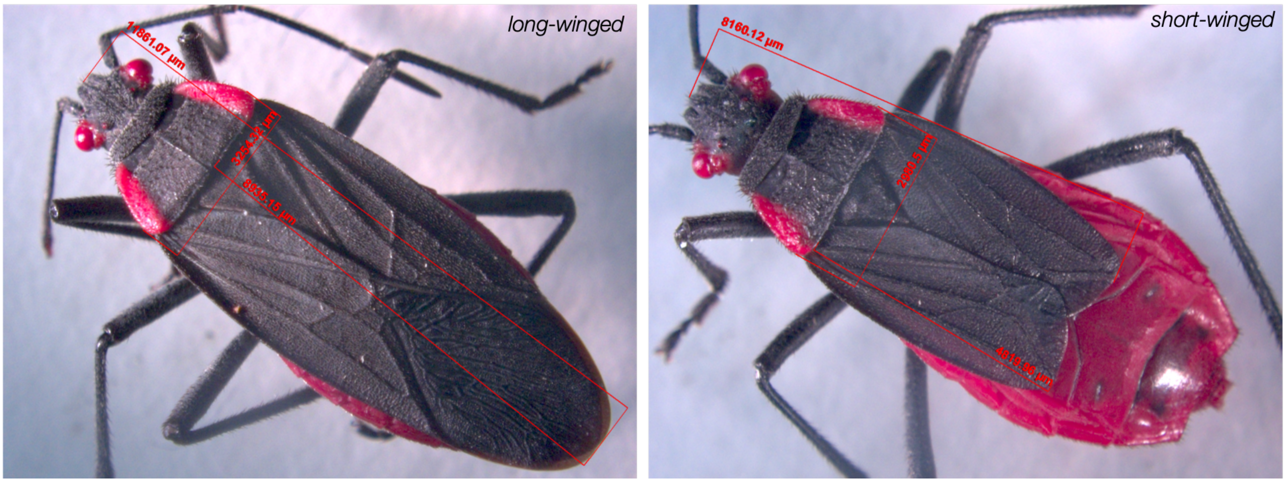

To better visualize how wing morph relates to a SBB allometric measurement, wing-to-thorax ratio, the following histograms were plotted:

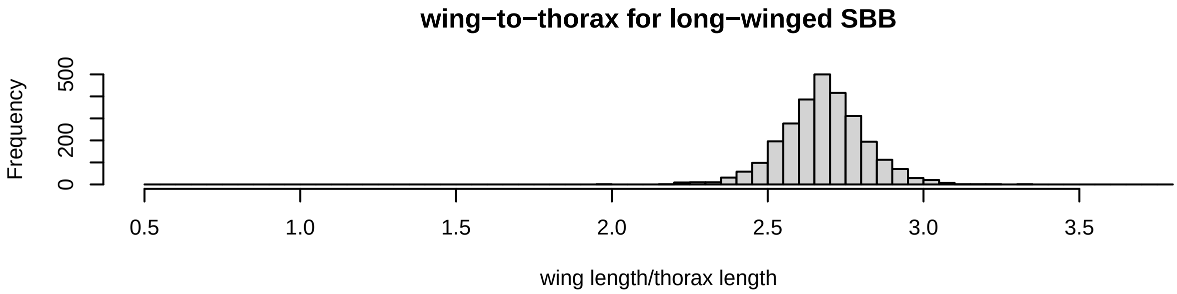

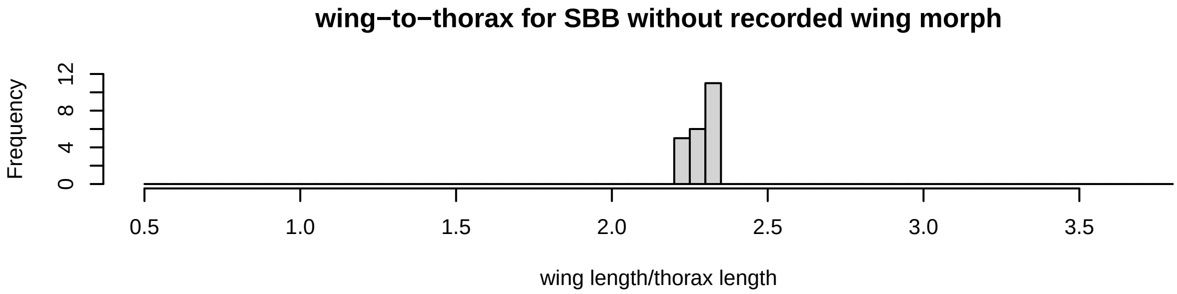

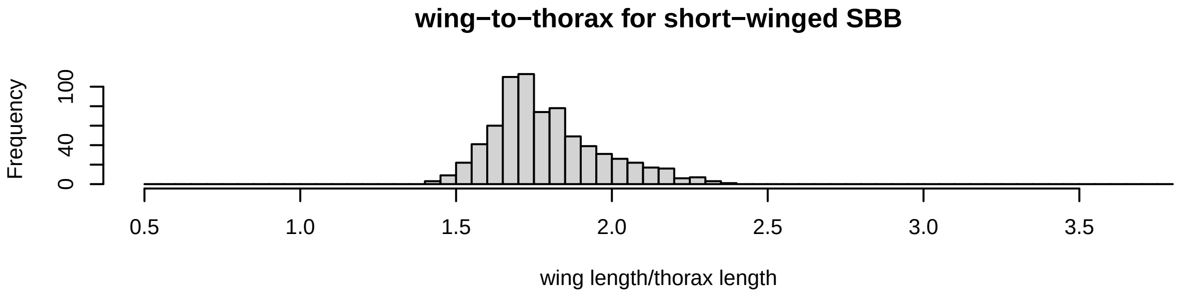

From the histograms, the relationship between wing morph and wing-to-thorax ratio is bimodal. Long-winged bugs have larger wing-to-thorax ratios with a frequency peak around 2.75, whereas short-winged bugs have much smaller wing-to-thorax ratios with a frequency peak around 1.75. It is then noticeable that there are 22 bugs who had not been identified as either S or L during measurements, but cannot be categorized into S or L because their wing-to-thorax values reside in-between the two modes.

##### 2.5 Field Collection Numbers

Bugs were collected from the field during different years and months. They were shipped to Davis, CA [2013-2016]; Tallahassee, FL [2017]; Chapel Hill, NC [2018]; or Chicago, IL [2019-2020]. The barplots below display the bugs collected per **population**, **host plant**, and **sex** across the years and months:

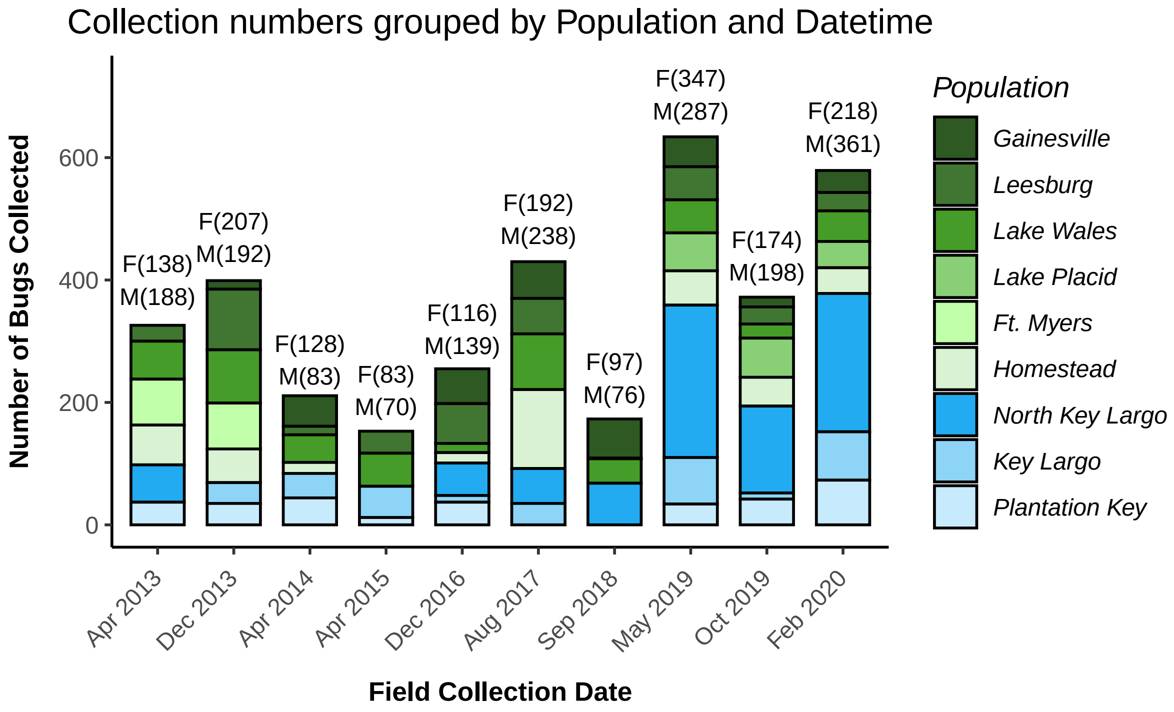

Stacked barplots are grouped and colored by population. Populations labeled in the legend are ordered by latitude from highest to lowest across Florida. Populations in shades of green indicate populations from the mainland of Florida while those in shades of blue indicate populations from the islands. Numbers above each bar represent the count between male (M) and female (F) soapberry bugs in each collection season. Population collections across datetimes are noticeably heterogeneous. However, collection numbers by sex and host plant were relatively homogeneous, as seen below.

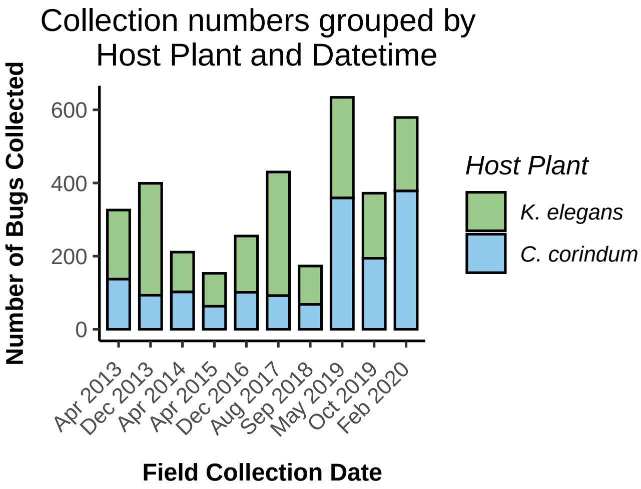

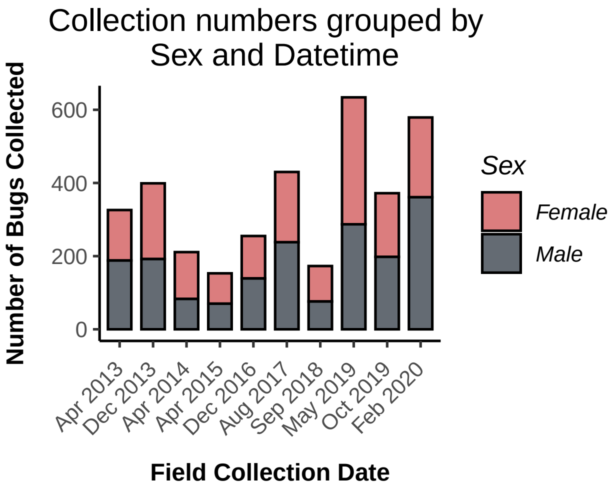

For more details on the Fall 2019 and Winter 2020 soapberry bugs collected, see the table below. These bugs were also tested for flight trials (see Appendix B). Flight tested bugs are grouped by collection site and ordered by descending latitude. Collection numbers across each collection site ($n_{site}$) and each population ($n_{pop}$) are also recorded under Sample Sizes.

| Fall 2019 | | | | | | |
|---|---|---|---|---|---|---|
| | | | Coordinates | | Sample Sizes | |
| host plant | population | site | latitude | longitude | n (site) | n (pop) |
| K. elegans | Gainesville | NW 10th Ave & 18th St | 29.66182 | -82.34721 | 14 | 14 |
| K. elegans | Leesburg | Veteran's Memorial Park | 28.81298 | -81.87789 | 13 | 13 |
| K. elegans | Lake Wales | Polk | 27.90335 | -81.58946 | 11 | 11 |
| K. elegans | Lake Placid | Barber shop | 27.29866 | -81.36612 | 43 | 43 |
| K. elegans | Homestead | SW 296th St & 182nd Ave | 25.49197 | -80.48562 | 34 | 34 |
| C. corindum | North Key Largo | DD | 25.28637 | -80.29077 | 26 | 65 |
| C. corindum | North Key Largo | DD front | 25.27352 | -80.30428 | 2 | 65 |
| C. corindum | North Key Largo | Carysfort Cr | 25.25648 | -80.31069 | 8 | 65 |
| C. corindum | North Key Largo | DD -inter | 25.25284 | -80.31183 | 2 | 65 |
| C. corindum | North Key Largo | Charlemagne | 25.19484 | -80.34590 | 9 | 65 |
| C. corindum | North Key Largo | N. Dagny | 25.18229 | -80.36370 | 5 | 65 |
| C. corindum | North Key Largo | Dagny Trellis | 25.17550 | -80.36782 | 8 | 65 |
| C. corindum | North Key Largo | Dagny 1/2 Loop | 25.17369 | -80.37153 | 5 | 65 |
| C. corindum | Key Largo | JP Grove | 25.12846 | -80.40809 | 14 | 14 |
| C. corindum | Plantation Key | Founder's #2 | 24.98519 | -80.54710 | 1 | 13 |
| C. corindum | Plantation Key | Founder's #1 | 24.96448 | -80.56739 | 12 | 13 |

| Winter 2020 | | | | | | |
|---|---|---|---|---|---|---|
| | | | Coordinates | | Sample Sizes | |
| host plant | population | site | latitude | longitude | n (site) | n (pop) |
| K. elegans | Gainesville | 23rd & 8th | 29.66200 | -82.34731 | 17 | 17 |
| K. elegans | Leesburg | Mount & 8th | 28.79634 | -81.87788 | 12 | 12 |
| K. elegans | Lake Wales | Polk Ave | 27.89674 | -81.58047 | 11 | 11 |
| K. elegans | Lake Placid | 110N Main | 27.29863 | -81.36615 | 24 | 24 |
| K. elegans | Homestead | SW 296th St | 25.49136 | -80.48582 | 27 | 27 |
| C. corindum | North Key Largo | Dynamite Docks | 25.28675 | -80.29059 | 7 | 143 |
| C. corindum | North Key Largo | DD front | 25.27349 | -80.30410 | 32 | 143 |
| C. corindum | North Key Largo | Carysfort | 25.25656 | -80.31080 | 18 | 143 |
| C. corindum | North Key Largo | MM165 | 25.22752 | -80.32838 | 17 | 143 |
| C. corindum | North Key Largo | Charlemagne | 25.19515 | -80.34592 | 21 | 143 |
| C. corindum | North Key Largo | N. Dagny | 25.18209 | -80.36319 | 30 | 143 |
| C. corindum | North Key Largo | Dagny Trellis | 25.17557 | -80.36780 | 18 | 143 |
| C. corindum | Key Largo | JP | 25.12846 | -80.40809 | 22 | 45 |
| C. corindum | Key Largo | KLMRL | 25.10002 | -80.43752 | 23 | 45 |
| C. corindum | Plantation Key | Aregood Ln | 24.97357 | -80.55392 | 18 | 53 |
| C. corindum | Plantation Key | Founder's | 24.96424 | -80.56733 | 35 | 53 |

Fall 2019 field and test collections were a mix of short wing and long wing soapberry bugs. Winter 2020 field collections were a mix of short wing and long wing bugs, but only long wing soapberry bugs were tested.

### 3 Regression Modeling

Multivariate, GLM was performed using the `glm()` function in the `lme4` package. Models were compared using Akaike Information Criterion (AIC) and model selection was determined using Akaike weights. Model fit was further evaluated between two models using the `anova()` function.

#### 3.1 Long-Wing Morph Frequency

We tested how sex, host plant, month, and/or year effected whether a soapberry bug is long-winged (`wing_morph_b=1`) or short-winged (`wing_morph_b=0`).

```
data = data.frame(R=raw_data$wing_morph_b,
                  A=raw_data$sex_b,
                  B=raw_data$pophost_b,
                  C=raw_data$month_of_year,
                  D=raw_data$months_since_start)

model_script = paste0(source_path,"generic models-binomial glm 4-FF.R")
model_comparisonsAIC(model_script)
```

```
##        [,1]      [,2]      [,3]      [,4]       [,5]
## AICs   3145.306  3146.842  3147.157  3147.201   3148.521
## models 98        110       84        107        105
## probs  0.2529382 0.1173602 0.1002697 0.09808583 0.05068685
##
## m98  glm(formula = R ~ A * B + A * D + B * C + C * D, family = binomial,
##     data = data)
## m110     glm(formula = R ~ A * B + A * D + B * C + B * D + C * D, family = binomial,
##     data = data)
## m84  glm(formula = R ~ A * D + B * C + C * D, family = binomial, data = data)
## m107     glm(formula = R ~ A * B + A * C + A * D + B * C + C * D, family = binomial,
##     data = data)
## m105     glm(formula = R ~ A * D + B * C + B * D + C * D, family = binomial,
##     data = data)
```

The R output above can be read as follows: Models exhibiting an Akaike weight greater than 0.05 are selected and displayed on the top table. The table is ordered by decreasing Akaike weight (or increasing AIC) where, for example, model `m98` had the highest Akaike weigh and lowest AIC. These Akaike weighs would demonstrate the relative likelihood of each model and they can be interpreted as the probabilities that a given model is the best approximating model.

Following the table, the formula of each model is pasted in order to make the models easy to refer to during the upcoming model comparisons using `anova()`.

```
anova(m98, m110, test="Chisq") # adding B*D does not improve fit
anova(m84, m98, test="Chisq")  # adding A*B improves fit
anova(m63, m84, test="Chisq")  # Adding C*D improves fit
anova(m51, m63, test="Chisq")  # Adding B improves fit
```

```
## Analysis of Deviance Table
##
## Model 1: R ~ A * B + A * D + B * C + C * D
## Model 2: R ~ A * B + A * D + B * C + B * D + C * D
##   Resid. Df Resid. Dev Df Deviance Pr(>Chi)
## 1      3461     3127.3
## 2      3460     3126.8  1  0.46421   0.4957
## Analysis of Deviance Table
```

```
##
## Model 1: R ~ A * D + B * C + C * D
## Model 2: R ~ A * B + A * D + B * C + C * D
##   Resid. Df Resid. Dev Df Deviance Pr(>Chi)
## 1      3462     3131.2
## 2      3461     3127.3  1   3.8506  0.04973 *
## ---
## Signif. codes:  0 '***' 0.001 '**' 0.01 '*' 0.05 '.' 0.1 ' ' 1
## Analysis of Deviance Table
##
## Model 1: R ~ A * D + C * D + B
## Model 2: R ~ A * D + B * C + C * D
##   Resid. Df Resid. Dev Df Deviance Pr(>Chi)
## 1      3463     3137.3
## 2      3462     3131.2  1   6.1886  0.01286 *
## ---
## Signif. codes:  0 '***' 0.001 '**' 0.01 '*' 0.05 '.' 0.1 ' ' 1
## Analysis of Deviance Table
##
## Model 1: R ~ A * D + C * D
## Model 2: R ~ A * D + C * D + B
##   Resid. Df Resid. Dev Df Deviance  Pr(>Chi)
## 1      3464     3497.3
## 2      3463     3137.3  1   359.93 < 2.2e-16 ***
## ---
## Signif. codes:  0 '***' 0.001 '**' 0.01 '*' 0.05 '.' 0.1 ' ' 1
```

The best fit model is m98. That is confirmed by its minimum AIC value, maximum Akaike weight, and the addition of `A*B` (`sex_b*pophost_b`) leading to a significant improvement in model fit as detected by the ANOVA test.

###### 3.1.1 Best Fit

```r
M1 = glm(wing_morph_b ~ sex_b * pophost_b + sex_b * months_since_start +
           pophost_b * month_of_year + month_of_year * months_since_start,
         data=raw_data, family="binomial")
summary(M1)
```

```
##
## Call:
## glm(formula = wing_morph_b ~ sex_b * pophost_b + sex_b * months_since_start +
##     pophost_b * month_of_year + month_of_year * months_since_start,
##     family = "binomial", data = raw_data)
##
## Deviance Residuals:
##     Min       1Q   Median       3Q      Max
## -2.3803   0.3597   0.4321   0.8450   1.2552
##
## Coefficients:
##                                    Estimate Std. Error z value Pr(>|z|)
## (Intercept)                       0.7516501  0.1841942   4.081 4.49e-05 ***
## sex_b                            -0.2597900  0.0902673  -2.878 0.004002 **
## pophost_b                         1.1256358  0.1142931   9.849  < 2e-16 ***
## months_since_start                0.0107239  0.0029582   3.625 0.000289 ***
```

```
## month_of_year                        0.0995560  0.0255307   3.899 9.64e-05 ***
## sex_b:pophost_b                      0.0973323  0.0495811   1.963 0.049635 *
## sex_b:months_since_start             0.0037212  0.0015337   2.426 0.015254 *
## pophost_b:month_of_year             -0.0379395  0.0150617  -2.519 0.011771 *
## months_since_start:month_of_year -0.0014557  0.0004553  -3.198 0.001386 **
## ---
## Signif. codes:  0 '***' 0.001 '**' 0.01 '*' 0.05 '.' 0.1 ' ' 1
##
## (Dispersion parameter for binomial family taken to be 1)
##
##     Null deviance: 3562.3  on 3469  degrees of freedom
## Residual deviance: 3127.3  on 3461  degrees of freedom
##   (62 observations deleted due to missingness)
## AIC: 3145.3
##
## Number of Fisher Scoring iterations: 5
```

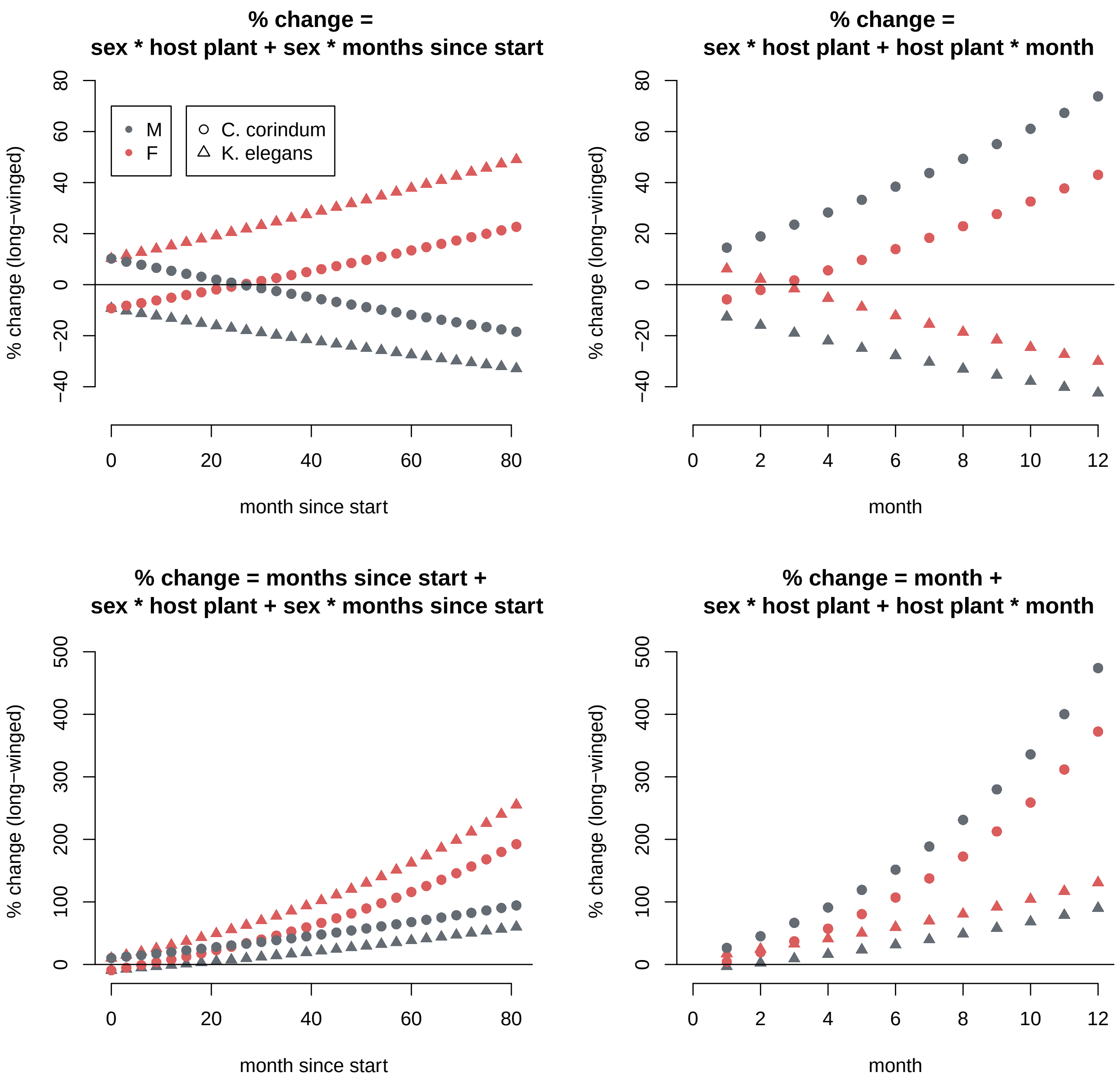

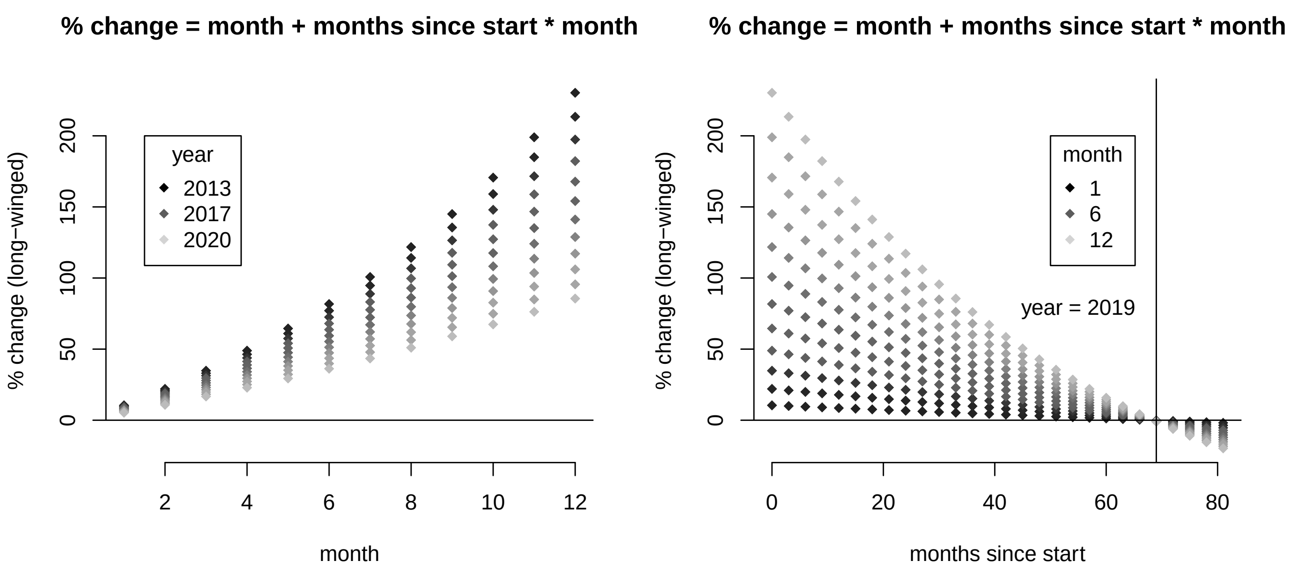

All single effects and their interactions are significant in the best fit model for predicting wing morph. It may be because of the size of the dataset that the model is more sensitive at detecting weak interactions as significant.

Also, notice how the interaction terms are weaker than the single variate effects in the best fit model. The first row of plots is showing only the interaction terms when considered independently whereas the second row of plots includes a single effect related to time (month or months since start). The strongest single variate effects, sex and host plant, are not plotted, but it is implied that they would drastically influence the possible outcome of whether a soapberry bug is long-winged or short-winged. This becomes more evident in the LOESS plots section of this appendix.

##### 3.2 Long-Wing Morph Variance

We then tested how sex, host plant, month, and/or year effected long-wing morph frequency variance.

First, the long-wing morph mean frequency was computed using `aggregate()` to group the long-wing morph recordings in `raw_data` according to sex, host plant, month, and year. The subsequent subset data created was `wmorph_table` (n=40) Then, summary statistics were applied to the data subset and variance (`sd`) was modeled.

```
wmorph_table = aggregate(wing_morph_b ~
                           sex_b*pophost_b*month_of_year*months_since_start,
                           data=raw_data, FUN=mean)

SE = function(x){sd(x)/sqrt(length(x))}

wmorph_table$sd = aggregate(wing_morph_b ~
                              sex_b*pophost_b*month_of_year*months_since_start,
                              data=raw_data,FUN=sd)$wing_morph_b
wmorph_table$se = aggregate(wing_morph_b ~
                              sex_b*pophost_b*month_of_year*months_since_start,
                              data=raw_data,FUN=SE)$wing_morph_b
wmorph_table$n = aggregate(wing_morph_b ~
                             sex_b*pophost_b*month_of_year*months_since_start,
                              data=raw_data,FUN=length)$wing_morph_b
```

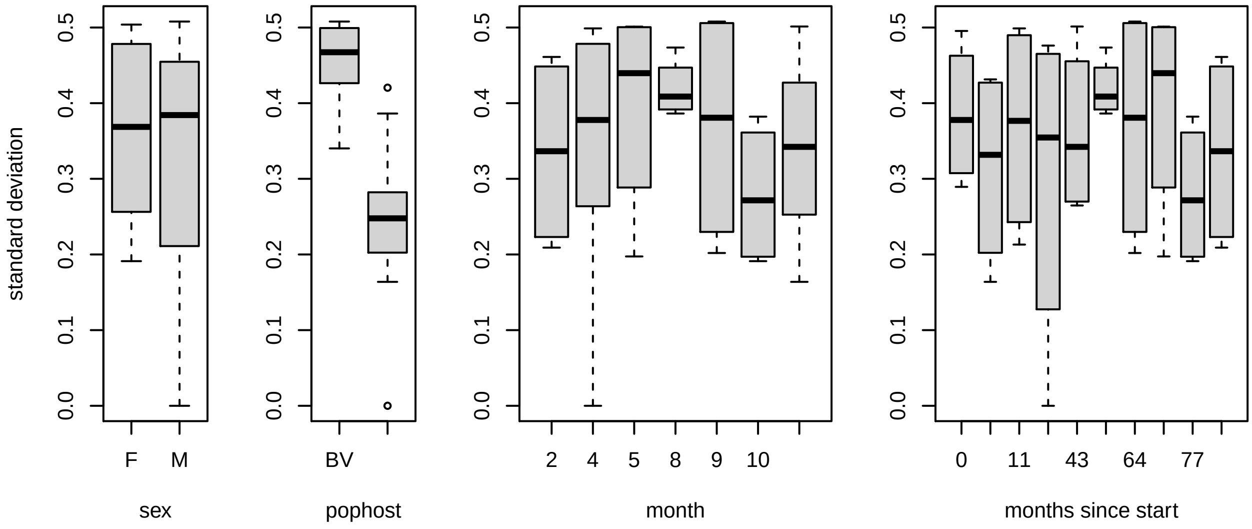

```r
data = wmorph_table
data = data.frame(R=data$sd,
                  A=data$sex_b,
                  B=data$pophost_b,
                  C=data$month_of_year,
                  D=data$months_since_start)

model_script = paste0(source_path,"generic models-gaussian glm 4-FF.R")
model_comparisonsAIC(model_script)
```

```
##        [,1]      [,2]      [,3]       [,4]
## AICs   -92.39855 -90.95292 -90.75898  -90.41465
## models 2         5         8          9
## probs  0.183788  0.0892081 0.08096352 0.06815837
##
## m2   glm(formula = R ~ B, family = gaussian, data = data)
## m5   glm(formula = R ~ A + B, family = gaussian, data = data)
## m8   glm(formula = R ~ B + C, family = gaussian, data = data)
## m9   glm(formula = R ~ B + D, family = gaussian, data = data)
```

```r
anova(m2, m5, test="Chisq") # Adding A does not improve fit
anova(m2, m8, test="Chisq") # Adding C does not improve fit
anova(m2, m9, test="Chisq") # Adding D does not improve fit
anova(m0, m2, test="Chisq") # Adding B improves fit
```

```
## Analysis of Deviance Table
##
## Model 1: R ~ B
## Model 2: R ~ A + B
##   Resid. Df Resid. Dev Df  Deviance Pr(>Chi)
## 1        38    0.20009
## 2        37    0.19734  1 0.0027541   0.4724
## Analysis of Deviance Table
##
## Model 1: R ~ B
## Model 2: R ~ B + C
##   Resid. Df Resid. Dev Df  Deviance Pr(>Chi)
## 1        38    0.20009
```

```
## 2        37    0.19830  1 0.0017949   0.5628
## Analysis of Deviance Table
##
## Model 1: R ~ B
## Model 2: R ~ B + D
##   Resid. Df Resid. Dev Df   Deviance Pr(>Chi)
## 1        38    0.20009
## 2        37    0.20001  1 8.0534e-05   0.9029
## Analysis of Deviance Table
##
## Model 1: R ~ 1
## Model 2: R ~ B
##   Resid. Df Resid. Dev Df Deviance  Pr(>Chi)
## 1        39    0.62439
## 2        38    0.20010  1  0.42429 < 2.2e-16 ***
## ---
## Signif. codes:  0 '***' 0.001 '**' 0.01 '*' 0.05 '.' 0.1 ' ' 1
```

The best fit model is m2. That is confirmed by its minimum AIC value, maximum Akaike weight, and the addition of `B` (`pophost_b`) to the null model leading to a significant improvement in model fit as detected by the ANOVA test.

##### 3.2.1 Best Fit

```
M2 = glm(sd ~ pophost_b, data=wmorph_table, family="gaussian")
summary(M2)
```

```
##
## Call:
## glm(formula = sd ~ pophost_b, family = "gaussian", data = wmorph_table)
##
## Deviance Residuals:
##       Min         1Q     Median         3Q        Max
## -0.249168  -0.041487   0.005877   0.041147   0.171269
##
## Coefficients:
##             Estimate Std. Error t value Pr(>|t|)
## (Intercept)  0.35216    0.01147  30.693  < 2e-16 ***
## pophost_b   -0.10299    0.01147  -8.976 6.28e-11 ***
## ---
## Signif. codes:  0 '***' 0.001 '**' 0.01 '*' 0.05 '.' 0.1 ' ' 1
##
## (Dispersion parameter for gaussian family taken to be 0.005265667)
##
##     Null deviance: 0.62439  on 39  degrees of freedom
## Residual deviance: 0.20010  on 38  degrees of freedom
## AIC: -92.399
##
## Number of Fisher Scoring iterations: 2
```

Host plant (`K. elegans = 1`, `C. corindum = -1`) is significant in predicting long-wing morph frequency variance. Soapberry bugs collected from *C. corindum*, balloon vine, experience more variance in long-wing morph frequency than those collected from *K. elegans*, goldenrain tree.

##### 3.3 Wing-to-Body Ratio

We tested how sex, host plant, month, and/or year effected whether the wing-to-body ratio of long-winged soapberry bugs.

```
data = data.frame(R=data_long$wing2body_c,
                  A=data_long$sex_b,
                  B=data_long$pophost_b,
                  C=data_long$month_of_year_c,
                  D=data_long$months_since_start_c)

model_script = paste0(source_path,"generic models-gaussian glm 4-FF.R")
model_comparisonsAIC(model_script)
```

```
##         [,1]      [,2]      [,3]       [,4]       [,5]       [,6]
## AICs   -9722.301 -9721.371 -9720.852  -9720.339  -9720.331  -9719.674
## models 88        99        58         92         97         76
## probs  0.1948772 0.1224324 0.09441271 0.07306166 0.07277994 0.05239229
##
## m88  glm(formula = R ~ A * B + A * D + B * D + C, family = gaussian,
##     data = data)
## m99  glm(formula = R ~ A * B + A * D + B * D + C * D, family = gaussian,
##     data = data)
## m58  glm(formula = R ~ A * B + B * D + C, family = gaussian, data = data)
## m92  glm(formula = R ~ A * B + A * C + A * D + B * D, family = gaussian,
##     data = data)
## m97  glm(formula = R ~ A * B + A * D + B * C + B * D, family = gaussian,
##     data = data)
## m76  glm(formula = R ~ A * B + B * D + C * D, family = gaussian, data = data)
```

```
anova(m88, m99, test="Chisq") # adding C*D does not improve fit
anova(m58, m88, test="Chisq") # Adding A*D marginally improves fit
anova(m58, m76, test="Chisq") # Adding C*D does not improve fit
anova(m34, m58, test="Chisq") # Adding B*D improves fit
```

```
## Analysis of Deviance Table
##
## Model 1: R ~ A * B + A * D + B * D + C
## Model 2: R ~ A * B + A * D + B * D + C * D
##   Resid. Df Resid. Dev Df   Deviance Pr(>Chi)
## 1      1895    0.66692
## 2      1894    0.66655  1 0.00037502   0.3019
## Analysis of Deviance Table
##
## Model 1: R ~ A * B + B * D + C
## Model 2: R ~ A * B + A * D + B * D + C
##   Resid. Df Resid. Dev Df Deviance Pr(>Chi)
## 1      1896    0.66813
## 2      1895    0.66692  1  0.00121  0.06371 .
## ---
## Signif. codes:  0 '***' 0.001 '**' 0.01 '*' 0.05 '.' 0.1 ' ' 1
## Analysis of Deviance Table
##
## Model 1: R ~ A * B + B * D + C
```

```
## Model 2: R ~ A * B + B * D + C * D
##   Resid. Df Resid. Dev Df  Deviance Pr(>Chi)
## 1      1896    0.66813
## 2      1895    0.66784  1 0.0002886   0.3655
## Analysis of Deviance Table
##
## Model 1: R ~ A * B + C + D
## Model 2: R ~ A * B + B * D + C
##   Resid. Df Resid. Dev Df  Deviance Pr(>Chi)
## 1      1897    0.67063
## 2      1896    0.66813  1 0.0024994  0.00774 **
## ---
## Signif. codes:  0 '***' 0.001 '**' 0.01 '*' 0.05 '.' 0.1 ' ' 1
```

The best fit model is m58. It did not have the minimum AIC value or maximum Akaike weight, but the addition of `A*D (sex_b*months_since_start_c)` was not detected as a significant improvement in model fit, according to the ANOVA test.

##### 3.3.1 Best Fit

```
M3 = glm(wing2body_c ~ sex_b*pophost_b + pophost_b*months_since_start_c
        + month_of_year_c, data=data_long, family=gaussian)
summary(M3)
```

```
##
## Call:
## glm(formula = wing2body_c ~ sex_b * pophost_b + pophost_b * months_since_start_c +
##     month_of_year_c, family = gaussian, data = data_long)
##
## Deviance Residuals:
##       Min         1Q     Median         3Q        Max
## -0.070837  -0.010794  -0.000093   0.010596   0.113993
##
## Coefficients:
##                                  Estimate Std. Error t value Pr(>|t|)
## (Intercept)                    -4.542e-04  4.601e-04  -0.987  0.32368
## sex_b                          -1.787e-03  4.467e-04  -4.001 6.55e-05 ***
## pophost_b                       4.289e-03  4.613e-04   9.297  < 2e-16 ***
## months_since_start_c           -1.727e-05  2.225e-05  -0.776  0.43763
## month_of_year_c                 7.155e-04  1.379e-04   5.188 2.35e-07 ***
## sex_b:pophost_b                 1.804e-03  4.466e-04   4.038 5.60e-05 ***
## pophost_b:months_since_start_c  5.904e-05  2.217e-05   2.663  0.00781 **
## ---
## Signif. codes:  0 '***' 0.001 '**' 0.01 '*' 0.05 '.' 0.1 ' ' 1
##
## (Dispersion parameter for gaussian family taken to be 0.0003523901)
##
##     Null deviance: 0.72538  on 1902  degrees of freedom
## Residual deviance: 0.66813  on 1896  degrees of freedom
## AIC: -9720.9
##
## Number of Fisher Scoring iterations: 2
```

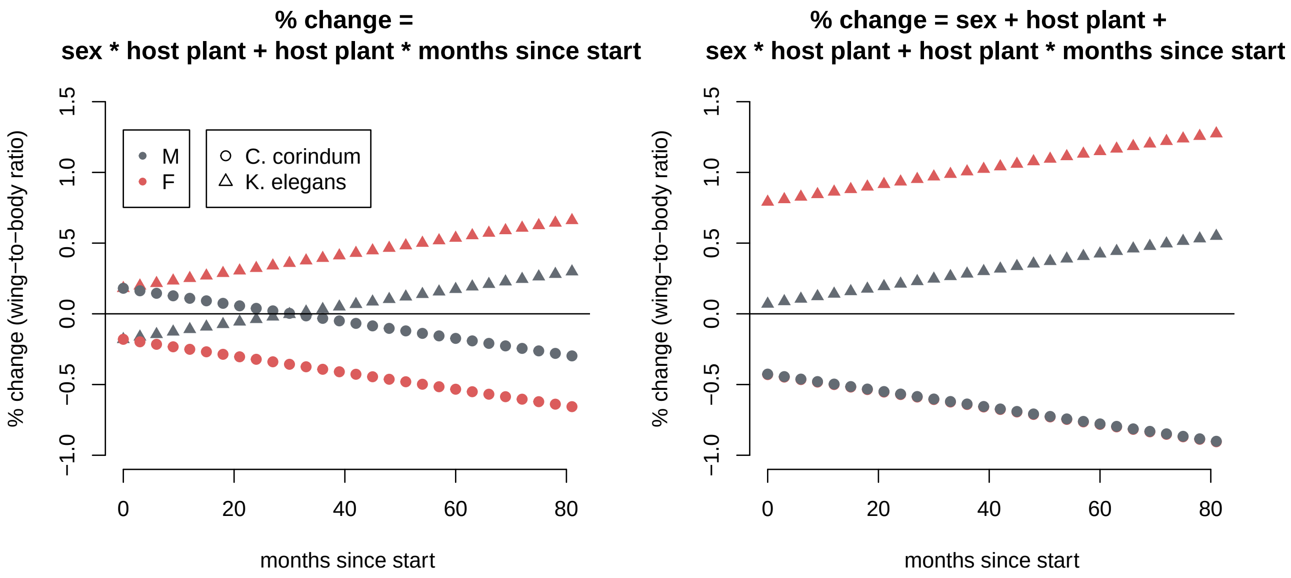

All single effects except `months_since_start` and all interactions are significant in the best fit model for predicting wing-to-body ratio. It is noticeable that month and year effect sizes are relatively small.

In general, it is noticeable that the variables in the model have weak effect sizes, especially when compared to how long-wing morph frequency changes over time. Considering single effects, does not lead to more pronounced percent changes in wing-to-body ratio across the years, but it does highlight which host plants exhibit sex differences and how host plant differences seems to be most influencing changes in wing-to-body ratio over time.

##### 3.4 Wing-to-Body Ratio Variance

We then tested how sex, host plant, month, or year effects the wing-to-body ratio variance of long-winged soapberry bugs.

First, the mean wing-to-body ratio was computed using `aggregate()` to group the wing-to-body ratio recordings in `data_long` according to sex, host plant, month, and year. The subsequent subset data created was `w2b_table` (n=36). Then, summary statistics were applied to the data subset and variance (`sd`) was modeled.

```
w2b_table = aggregate(wing2body ~
                        sex_b*pophost_b*month_of_year*months_since_start,
                        data=data_long, FUN=mean)
w2b_table$sd = aggregate(wing2body ~
                           sex_b*pophost_b*month_of_year*months_since_start,
                           data=data_long, FUN=sd)$wing2body
w2b_table$se = aggregate(wing2body ~
                           sex_b*pophost_b*month_of_year*months_since_start,
                           data=data_long, FUN=SE)$wing2body
```

```
data = w2b_table
data = data.frame(R=data$sd,
                  A=data$sex_b,
                  B=data$pophost_b,
                  C=data$month_of_year,
                  D=data$months_since_start)

model_script = paste0(source_path,"generic models-gaussian glm 4-FF.R")
model_comparisonsAIC(model_script)
```

```
##        [,1]      [,2]      [,3]
## AICs   -280.1872 -279.8713 -279.4758
## models 8         19        2
## probs  0.1198675 0.1023577 0.08398967
##
## m8   glm(formula = R ~ B + C, family = gaussian, data = data)
## m19  glm(formula = R ~ B * C, family = gaussian, data = data)
## m2   glm(formula = R ~ B, family = gaussian, data = data)
```

```
anova(m8, m19, test="Chisq") # Adding B*C does not improve fit
anova(m2, m8, test="Chisq") # Adding C does not improve fit
anova(m0, m2, test="Chisq") # Adding B improves fit
```

```
## Analysis of Deviance Table
##
## Model 1: R ~ B + C
## Model 2: R ~ B * C
##   Resid. Df Resid. Dev Df  Deviance Pr(>Chi)
## 1        33 0.00070342
## 2        32 0.00067127  1 3.215e-05   0.2157
## Analysis of Deviance Table
##
## Model 1: R ~ B
## Model 2: R ~ B + C
##   Resid. Df Resid. Dev Df   Deviance Pr(>Chi)
## 1        34 0.00075844
## 2        33 0.00070342  1 5.5025e-05   0.1081
## Analysis of Deviance Table
##
## Model 1: R ~ 1
## Model 2: R ~ B
##   Resid. Df Resid. Dev Df   Deviance Pr(>Chi)
## 1        35 0.00087733
## 2        34 0.00075844  1 0.00011888  0.02097 *
## ---
## Signif. codes:  0 '***' 0.001 '**' 0.01 '*' 0.05 '.' 0.1 ' ' 1
```

The best fit model is m2. It did not have the minimum AIC value or maximum Akaike weight, but the addition of `B` (`pophost_b`) to the null model lead to a significant improvement in model fit detected by the ANOVA test.

##### 3.4.1 Best Fit

```
M4 = glm(sd ~ pophost_b, data=w2b_table, family=gaussian)
summary(M4)
```

```
##
## Call:
## glm(formula = sd ~ pophost_b, family = gaussian, data = w2b_table)
##
## Deviance Residuals:
##        Min         1Q     Median         3Q        Max
## -0.0059374  -0.0033018  -0.0006274   0.0022332   0.0147212
```

```
##
## Coefficients:
##              Estimate Std. Error t value Pr(>|t|)
## (Intercept) 0.0165999  0.0007872  21.088   <2e-16 ***
## pophost_b   0.0018172  0.0007872   2.309   0.0272 *
## ---
## Signif. codes:  0 '***' 0.001 '**' 0.01 '*' 0.05 '.' 0.1 ' ' 1
##
## (Dispersion parameter for gaussian family taken to be 2.230719e-05)
##
##     Null deviance: 0.00087733  on 35  degrees of freedom
## Residual deviance: 0.00075844  on 34  degrees of freedom
## AIC: -279.48
##
## Number of Fisher Scoring iterations: 2
```

Host plant (`K. elegans = 1,C. corindum = -1`) is significant in predicting wing-to-body ratio variance. Soapberry bugs collected from *K. elegans*, goldenrain tree, experience more variance in wing-to-body ratio than those collected from *C. corindum*, balloon vine.

### 4 LOESS & Linear Regression Plots

Locally-weighted scatterplot smoothing (LOESS) helped display and explore the non-linear fluctuations in long-wing morph frequency and wing-to-body ratio across time. Each data set was fit with a local polynomial regression using `lowess()` to determine LOESS parameters ($\alpha$ and $\lambda$) and `geom_smooth()` for plotting more aesthetic visuals.

#### 4.1 Wing Morph Frequency

##### 4.1.1 Group significant elements

Data are aggregated according to predictors present in their respective aforementioned best fit GLM model. For predicting long-wing morph frequency (`raw_data`), the best fit model had the following predictors: sex, host plant, month, and year. We used `dates`, a datetime object, instead of `months_since_start` for cleaner plotting, but the two are interchangeable.

```
# function to calculate 95% confidence interval (CI).
CI_95 = function(x){qnorm(0.975)*sd(x)/sqrt(length(x))}
CI_95_binom_upper = function(y) {
                        binom.confint(x=sum(y, na.rm=TRUE),
                                      n=length(y[!is.na(y)]),
                                      conf.level=0.95,
                                      methods='exact')$upper}
CI_95_binom_lower = function(y) {
                        binom.confint(x=sum(y, na.rm=TRUE),
                                      n=length(y[!is.na(y)]),
                                      conf.level=0.95,
                                      methods='exact')$lower}
```

```
# aggregate the full data
w_morph_summary = aggregate(wing_morph_b ~
                            sex*pophost*month_of_year*dates,
                            data=raw_data, FUN=mean)
```

```r
# compute standard error (SE), upper and lower CI, & sample size (n)
w_morph_summary$se = aggregate(wing_morph_b ~
                               sex*pophost*month_of_year*dates,
                               data=raw_data,
                               FUN=SE)$wing_morph_b
w_morph_summary$upper = aggregate(wing_morph_b ~
                                  sex*pophost*month_of_year*dates,
                                  data=raw_data,
                                  FUN=CI_95_binom_upper)$wing_morph_b
w_morph_summary$lower = aggregate(wing_morph_b ~
                                  sex*pophost*month_of_year*dates,
                                  data=raw_data,
                                  FUN=CI_95_binom_lower)$wing_morph_b
w_morph_summary$n = aggregate(wing_morph_b ~
                              sex*pophost*month_of_year*dates,
                              data=raw_data,
                              FUN=length)$wing_morph_b

dd = w_morph_summary
```

##### 4.1.2 Check for LOESS Residuals

To determine the span ($\alpha$, the smoothing parameter) and the degree of zero ($\lambda$) of the LOESS, smoothers were applied with increasing weights until the residuals appeared to have constant variance. Only the best LOESS parameters are shown below:

```r
plot_lowess_residuals = function(lfit, x, y, color) {
  lfun = approxfun(lfit)
  fitted = lfun(x)
  resid = y-fitted
  plot(fitted,resid,col=color, pch=19)
  abline(h=0,col=8)
}
```

```r
# loess models (month and year)
lM = lowess(dd$month_of_year, dd$wing_morph_b, f=0.4) # f = alpha, the smoother span
lY = lowess(dd$dates, dd$wing_morph_b, f=0.4)

# plot loess fit and residuals
par(mfrow=c(2,2), mai=c(0.80,0.80,0.3,0.3), mgp=c(2.3,1,0))

color=alpha("black", alpha = 0.75)
plot(dd$month_of_year, dd$wing_morph_b,
     xlab="month of year", ylab="long-wing morph freq", col=color)
lines(lM, type = "l")
plot_lowess_residuals(lM, dd$month_of_year, dd$wing_morph_b, color)

plot(dd$dates, dd$wing_morph_b,
     xlab="year", ylab="long-wing morph freq", col=color)
lines(lY, type = "l", color="#BEBEBE")
plot_lowess_residuals(lY, dd$dates, dd$wing_morph_b, color)
```

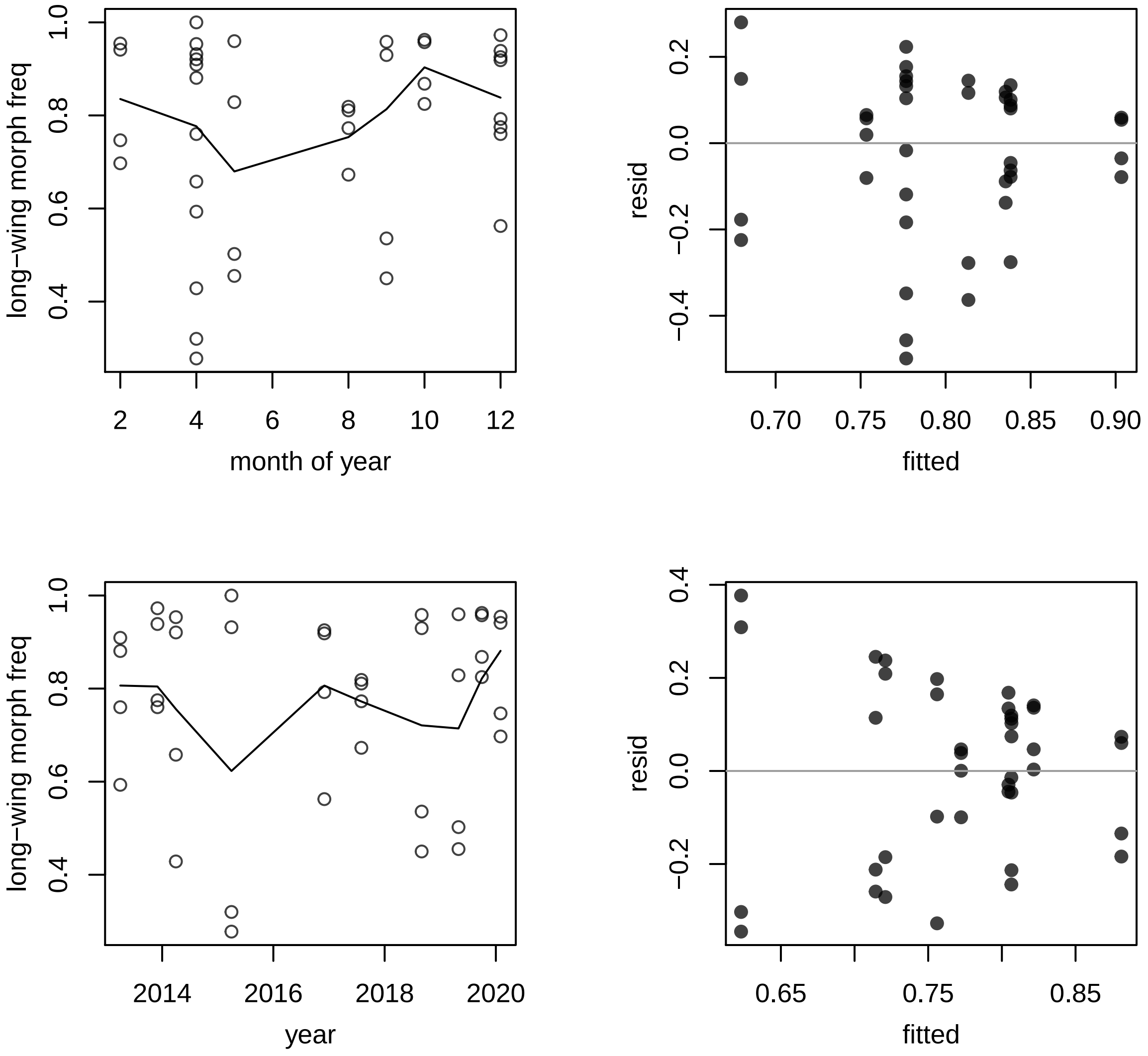

From these residual plots (right-side), we selected a $\lambda$=0 and $\alpha$=0.4. With a zero degree polynomial, LOESS acts as a weighted moving average and a span of 0.4 demonstrates independence between the residuals.

##### 4.1.3 Figure: Panels A, B, C, D (long-wing morph freq with month) & E (long-wing morph freq with year)

In addition to plotting local polynomial regression lines, the effects (slopes) of the best fit GLM models were also plotted. However, due to multiple interaction terms, we substituted the complex GLM models with single-variate or simpler models. This led to cleaner GLM line plotting, and the plots still reasonably reflected the aforementioned GLM models. Finally, all p-values displayed were extracted from the aforementioned best fit GLM model.

**Panels A and B Regression Computations:**

```
# single-variate model of month predicting wing morph
fit1 = glm(wing_morph_b ~ month_of_year, family="binomial", data=raw_data)
xmonth = seq(2,12, 0.01)
wing_probs = predict(fit1, list(month_of_year=xmonth), type="response")

# extract p-value from best fit regression model
fit_pvalue = round(summary(M1)$coeff[,"Pr(>|z|)"][5],5)
pvalue = paste0("italic(p)[glm]==", fit_pvalue)
```

**Panels C and D Regression Computations:**

```
# multiple variate model with month, sex, and host plant predicting wing morph
fit2 = glm(wing_morph_b ~ sex_b * pophost_b +
          pophost_b * month_of_year, family = "binomial", data = raw_data)

set.seed(194842)
xmon = seq(2,12, 0.01)
bsex = sample(c(-1,1), replace=TRUE, size=length(xmon))
bhost = sample(c(-1,1), replace=TRUE, size=length(xmon))
wprobs = predict(fit2, list(sex_b = bsex,
                            pophost_b = bhost,
                            month_of_year = xmon), type="response")

pred = cbind(xmon, bsex, bhost, wprobs)
pred = as.data.frame(pred)

predFK = pred[pred$bhost==1 & pred$bsex==1,]
predFC = pred[pred$bhost==-1 & pred$bsex==1,]

predMK = pred[pred$bhost==1 & pred$bsex==-1,]
predMC = pred[pred$bhost==-1 & pred$bsex==-1,]

# extract p-value from best fit regression model
fit_pvalue = round(summary(M1)$coeff[,"Pr(>|z|)"][6],4)
pvalue = paste0("italic(p)[glm]==", fit_pvalue)
```

**Panel F Regression Computations:**

```
# multiple variate model with year, sex, and host plant predicting wing morph
fit3 = glm(wing_morph_b ~ sex_b * dates, family = "binomial", data = raw_data)

set.seed(194842)
xyr = seq(sort(unique(dd$dates))[1],sort(unique(dd$dates))[10], 1)
bsex = sample(c(-1,1), replace=TRUE, size=length(xyr))
bhost = sample(c(-1,1), replace=TRUE, size=length(xyr))
wprobs = predict(fit3, list(sex_b = bsex,
                            pophost_b = bhost,
                            dates = xyr), type="response")

pred = cbind(xyr, bsex, bhost, wprobs)
pred = as.data.frame(pred)
pred$xyr = as.Date.numeric(pred$xyr)

predF = pred[pred$bsex==1,]
predM = pred[pred$bsex==-1,]

# extract p-value from best fit regression model
fit_pvalue = round(summary(M1)$coeff[,"Pr(>|z|)"][7],3)
pvalue = paste0("italic(p)[glm]==", fit_pvalue)
```

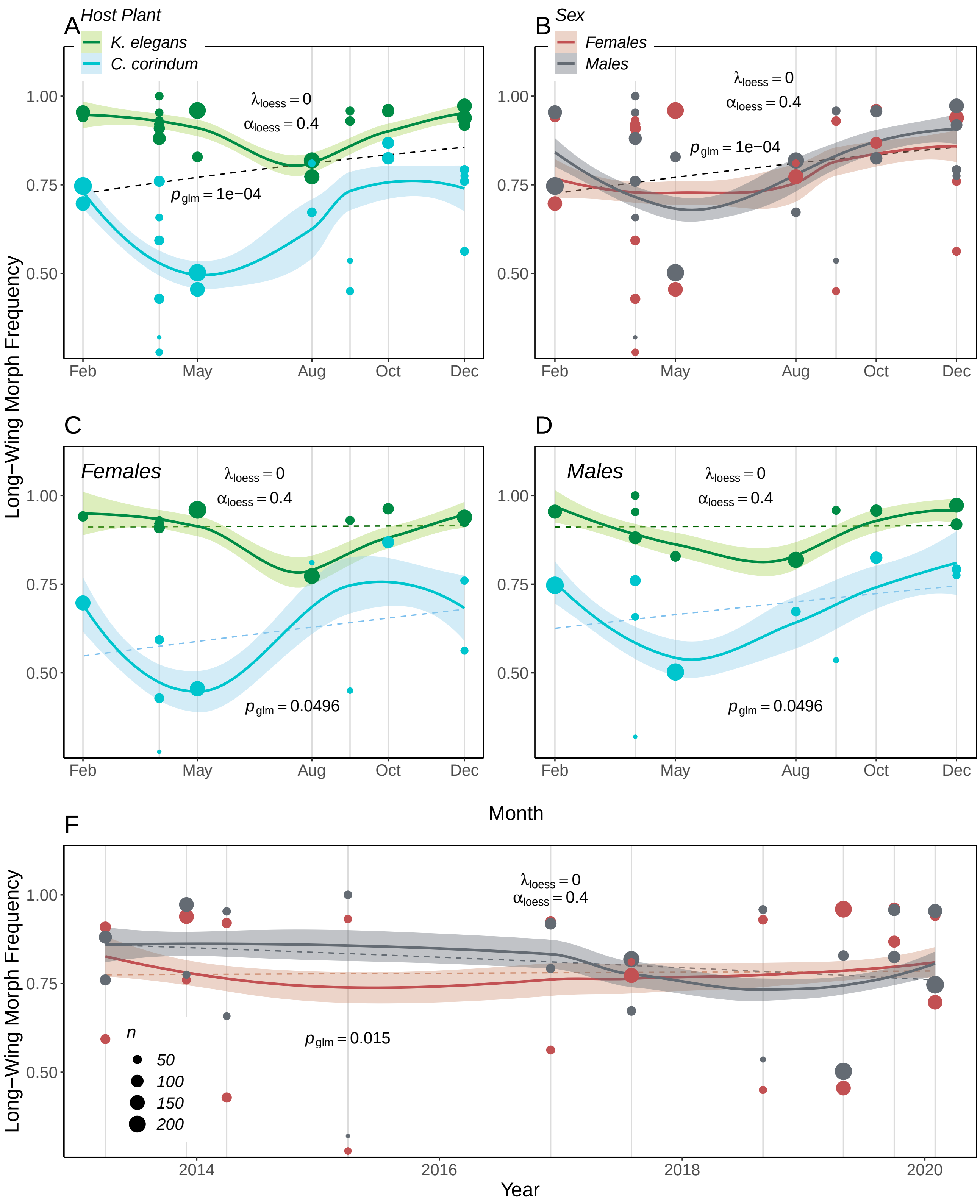

Evaluation of the frequency of long-winged morph soapberry bugs averaged across month and year from April 2013 to February 2020 using exploratory plots. For each point, the mean frequency of long-winged morphs of each month and year is plotted with LOESS smooth lines (solid lines) and 95% confidence intervals (shading) and linear regression line(s) (dashed line(s)).

##### 4.2 Wing-to-Body Ratio

###### 4.2.1 Group significant elements

Data are aggregated according to predictors present in their respective aforementioned best fit GLM model. For predicting wing-to-body ratio (`data_long`), the best fit model had the following predictors: sex, host plant, month, and year. We used `dates`, a datetime object, instead of `months_since_start` for cleaner plotting, but the two are interchangeable.

```
w2b_summary = aggregate(wing2body~sex*pophost*dates*month_of_year,
                        data=data_long, FUN=mean)
w2b_summary$se = aggregate(wing2body~sex*pophost*dates,
                           data=data_long,
                           FUN=SE)$wing2body
w2b_summary$n = aggregate(wing2body~sex*pophost*dates,
                          data=data_long,
                          FUN=length)$wing2body
d = w2b_summary
```

###### 4.2.2 Check for LOESS Residuals

To determine the span ($\alpha$, the smoothing parameter) and the degree of zero ($\lambda$) of the LOESS, smoothers were applied with increasing weights until the residuals appeared to have constant variance. Only the best LOESS parameters are shown below:

```
# loess models (month and year)
lM = lowess(d$month_of_year, d$wing2body, f=0.4) # f = alpha, the smoother span
lY = lowess(d$dates, d$wing2body, f=0.4)

# plot loess fit and residuals
par(mfrow=c(1,2), mai=c(0.80,0.80,0.3,0.3), mgp=c(2.5,1,0))

color="#00000080"
plot(d$month_of_year, d$wing2body,
     xlab="month of year", ylab="wing-to-body ratio", col=color)
lines(lM, type = "l")
plot_lowess_residuals(lM, d$month_of_year, d$wing2body, color)

plot(d$dates, d$wing2body,
     xlab="year", ylab="wing-to-body ratio", col=color)
lines(lY, type = "l")
plot_lowess_residuals(lY, d$dates, d$wing2body, color)
```

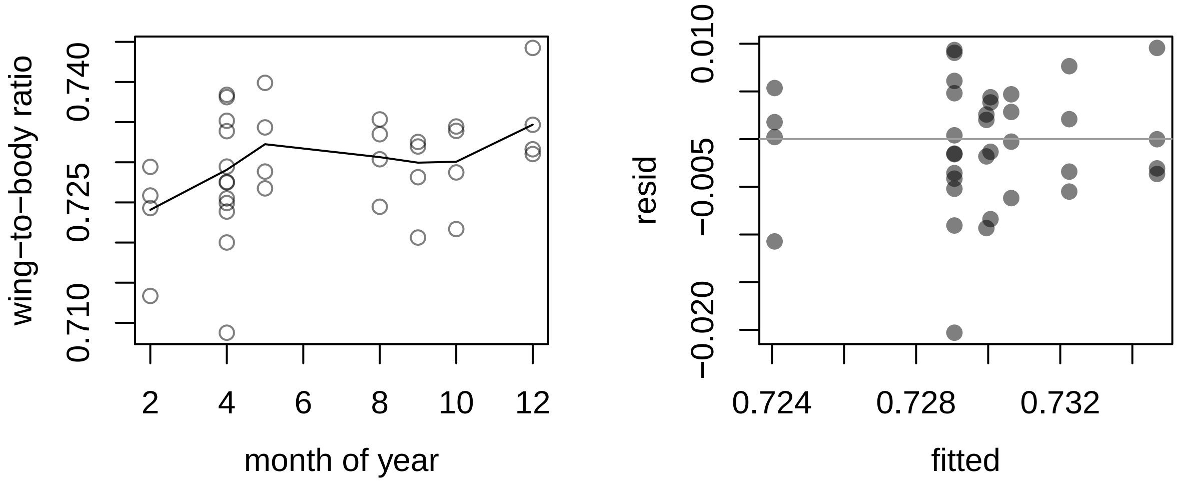

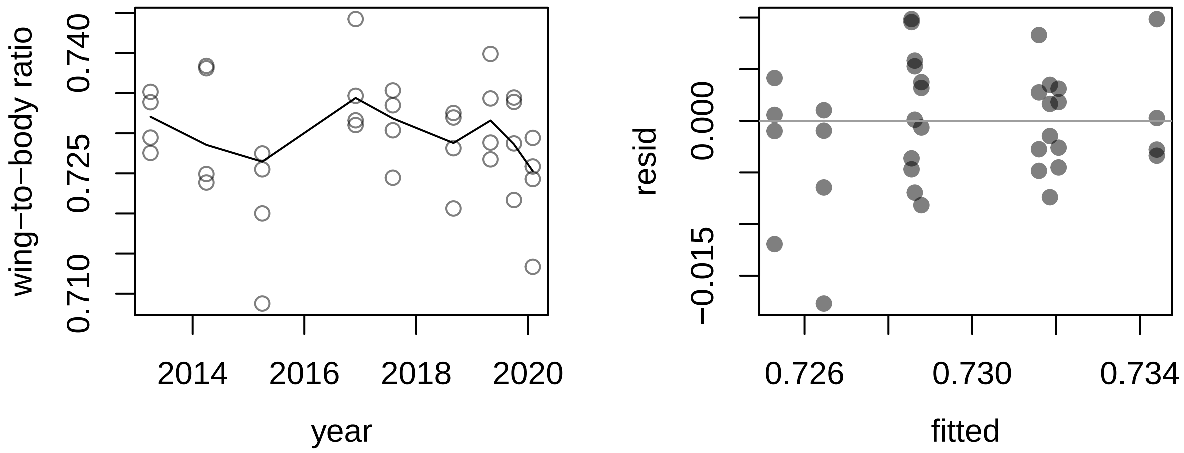

Similarly, from these residual plots (right-side), we selected a $\lambda$=0 and $\alpha$=0.4.

##### 4.2.3 Figure: Panels A, B, (wing-to-body ratio with month) & C (wing-to-body ratio with year)

In similar fashion, the local polynomial regression lines and the effects (slopes) of the best fit GLM models were plotted together. Due to multiple interaction terms, we substituted the complex GLM models with single-variate or simpler models. This led to cleaner GLM line plotting, and the plots still reasonably reflected the aforementioned GLM models. Finally, all p-values displayed were extracted from the aforementioned best fit GLM model.

**Panels A and B Regression Computations:**

Wing-to-body ratio is continuous data, unlike the wing morph data which is binary data. As a result, rather than using the `predict()` function to calculate the best fit line between wing-to-body and month, we used a single line of `ggplot` code, `geom_smooth(data=data_long, method="glm", mapping = aes(x = month_of_year, y = wing2body)...)`. This line of code can be see in the wing_summary.Rmd script.

**Panel C Regression Computations:**

```
# multiple variate model with year and host plant predicting wing2body ratio
fit4 = glm(wing2body ~ pophost_b * dates, data = data_long)

set.seed(194842)
xyr = seq(sort(unique(dd$dates))[1],sort(unique(dd$dates))[10], 1)
bhost = sample(c(-1,1), replace=TRUE, size=length(xyr))
wprobs = predict(fit4, list(pophost_b = bhost,
                                dates = xyr), type="response")

pred = cbind(xyr, bhost, wprobs)
pred = as.data.frame(pred)
pred$xyr = as.Date.numeric(pred$xyr)

predK = pred[pred$bhost==1,]
predC = pred[pred$bhost==-1,]

# extract p-value from best fit regression model
fit_pvalue = round(summary(M3)$coeff[,"Pr(>|t|)"][7],3)
pvalue = paste0("italic(p)[glm]==", fit_pvalue)
```

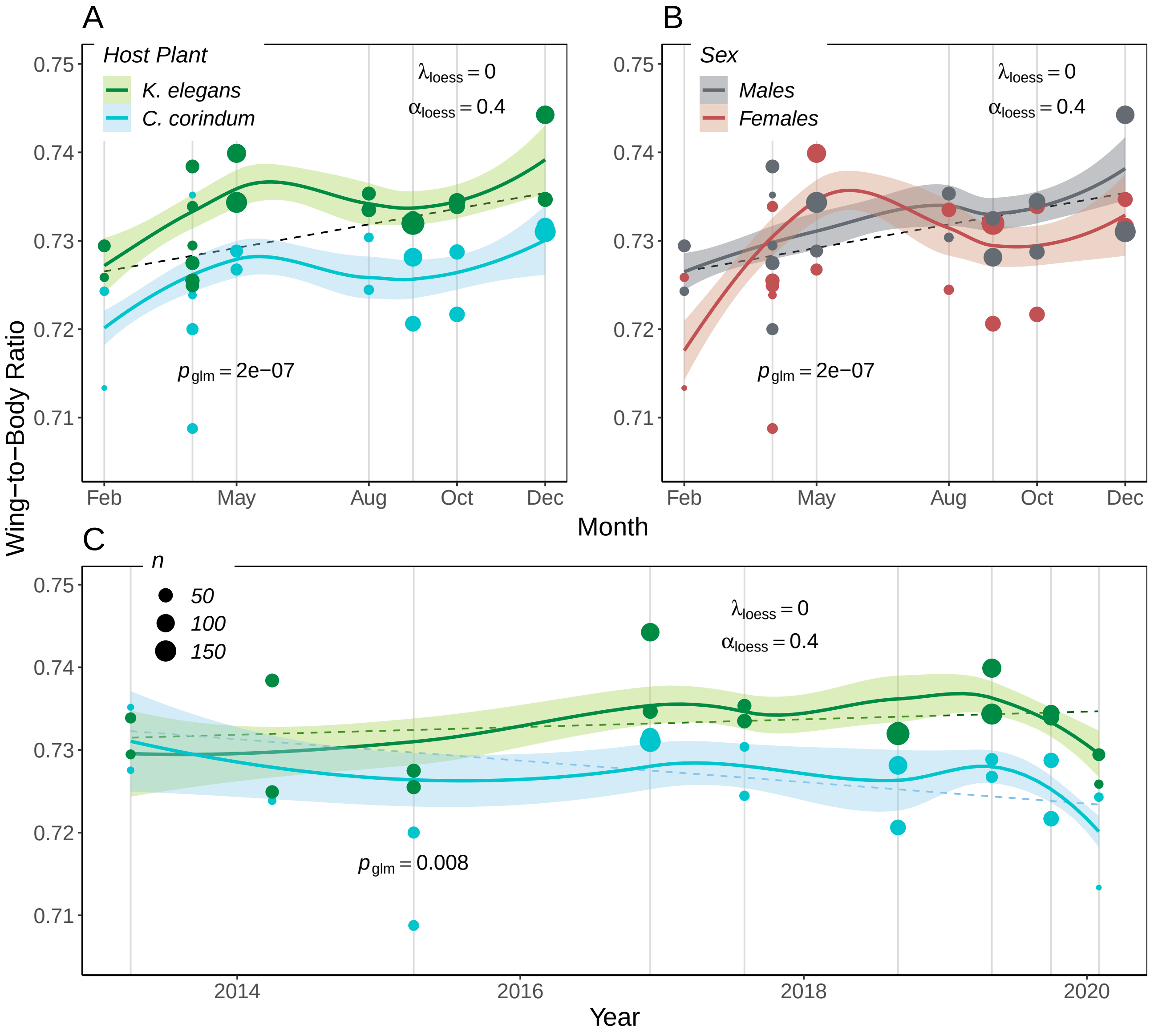

Evaluation of the wing-to-body ratio of soapberry bugs averaged across month and year from April 2013 to February 2020 using exploratory plots. For each point, the mean wing-to-body ratio of soapberry bugs collected in each month and year is plotted with LOESS smooth lines (solid lines) and 95% confidence intervals (shading) and linear regression line(s) (dashed line(s)).
