## Appendix B for "Dispersal-behavioral plasticity within an insect-host system undergoing human-induced rapid environmental change (HIREC)"

Bernat, AV, Cenzer, ML

### 1 Details of the Analyses

This document was generated by R Markdown on 2022-06-19 using R version 4.0.5 (2021-03-31). The document provides the step-by-step analytical methods used in the manuscript by Anastasia Bernat (AVB) and Meredith Cenzer (MLC). Multiple draft scripts were written by AVB and MLC between 2020-03-01 and 2021-07-26 until being distilled and complied by AVB and code reviewed by MLC at the University of Chicago into this comprehensive script. All draft scripts can be viewed in the GitHub repository, SBB-dispersal (https://github.com/mlcenzer/SBB-dispersal), within the directory **avbernat > Dispersal > Winter_2020 > stats** .

All code and output from the statistical analyses are shown. Code for data cleaning and the generation of plots is not displayed, but can be viewed in the **appendix_B-flight_summary.Rmd** file and its accompanying sourced scripts. To repeat analyses and the generation of plots, all data files and sourced scripts should follow the directory structure presented in the SBB-dispersal repository.

#### 1.1 Description of the Data

Soapberry bugs, *Jadera haematoloma*, were flight tested in the Fall 2019 (2019-10-15 to 2019-11-08) and Winter 2020 (2020-02-17 to 2020-03-10) seasons using a flight mill machine. Soapberry bugs were flight tested twice for either set time increments or multiple hours in the flight mill and observed from 8 AM to (5-8 PM) each day. For each trial, the mass, flight response, egg-laying response, distance, duration, average speed, and max speed of each soapberry bug were recorded and then processed.

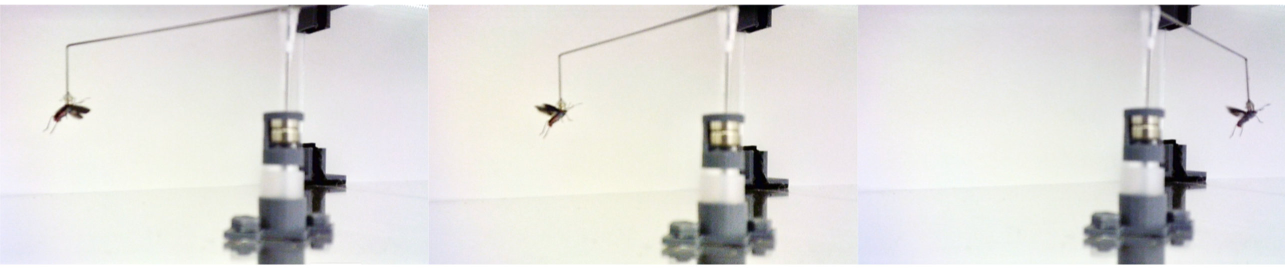

All Python scripts used to process the flight records are located in the GitHub repository within the directory **avbernat > Dispersal > Winter_2020 > windaq_processing** . After trials, morphology measurements were taken for each bug. There are four morphology measurements: beak length, thorax width, wing length, and body length. The sex, wing morph (long-winged, shot-winged, or ambiguously-winged), host plant, and population of each bug were also recorded.

As a result of the experimental design, this document analyzes two main types of datasets: a full dataset and a unique dataset. A **full dataset** is a dataset where each row has a unique bug ID and trial type combination. A **unique dataset** is a dataset where each row has a unique bug ID only because each trial has been grouped by ID. Examples are provided below. The advantage of generating a unique dataset is that changes between trials can be observed and analyzed.

#### 1.2 Abbreviations Used in the Data and Code

- **SBB** - soapberry bug, *Jadera haematoloma*
- **S** - short-winged morph
- **L** - long-winged morph
- **LS** or **SL** - ambiguous wing morph
- **T1** - trial 1 of flight testing
- **T2** - trial 2 of flight testing
- **EWM** - eggs when massed, binary response (yes or no)
- **host_** - the host plant soapberry bugs were collected from, which was either *Koelreuteria elegans* or *Cardiospermum corindum*, occasionally called (and abbreviated) as goldenrain tree (GRT) or balloon vine (BV), respectively
- **sym_dist** - distance from the local sympatric zone, which is demarked as Homestead, Florida

- **wing2body** - a computed and unitless column that calculates the wing length divided by the body length of a soapberry bug
- **sd** - standard deviation
- **se** - standard error
- **w__** - a column name that starts with `w_` is abbreviated from "wing". Example column: w_morph is "wing morph"

##### 1.3 Data Transformations

- `_b` - a column name that ends in `_b` is a column that has been recodified into binary data (0's and 1's). Example columns: flew_b, eggs_b

- `_c` - a column name that ends in `_c` is a column that has been centered. Example columns: sex_c, host_c, avg_days_c

- `_s` - a column name that ends in `_s` is a column that has been standardized. Example columns: wing2body_s, sym_dist_s, thorax_s

- `avg_` - a column name that starts in `avg_` is a column that has been averaged across trial 1 (T1) and trial 2 (T2). Example columns: avg_mass, avg_days, avg_time_start, avg_rec_dur (exception: average_speed)

- `_diff` - a column name that ends in `_diff` is a column that is the difference between T1 and T2 data values.

- `_per` - a column name that ends in `_per` is a column that is the percent change between T1 and T2 (T2-T1) data values. Formula: (T2-T1)/T1 * 100.

- `_logsqrt` - a column name that ends in `_logsqrt` is a column that has been normalized using a log-square-root transformation. Formula: `log(sqrt(<data_column>))-mean(log(sqrt(<data_column>))`. Example column(s): avg_mass_logsqrt

- `_logsqrt_i` - a column name that ends in `_logsqrt_i` is a column that has been normalized using a log-square-root transformation but its sign is the inverse of the column. Formula: `log(sqrt(0.85-column))-mean(log(sqrt(0.85-column))` where 0.85 is a number we selected that generates random errors that closely follow a normal distribution. Example column(s): wing2body_logsqrt_i

### Winter 2020 Flight Trials

### 2 Across-Trial Flight Response (T1 & T2)

#### 2.1 Read Libraries

The flight response of *J. haematoloma* was analyzed using multivariate, generalized linear modeling (GLM) as implemented in the R packages `lme4` and `binom`. Models were generated using the `glm()` function and compared using Akaike Information Criterion (AIC). Model selection was determined using Akaike weights, and model fit was further evaluated between two models using `anova()`.

All plots were generated using base R and supplemented with the `popbio` package to display logistic regressions and the `rethinking` package to display 95% confidence intervals of linear regressions. Additional R packages not show below but embedded in the sourced scripts are `lubridate`, `chron`, and `dplyr`. `lubridate` and `chron` both aid in datetime manipulation while `dplry` pipelines data grouping processes.

```r
library(lme4)       # fit regressions
library(rethinking) # Bayesian data analysis and plotting
library(popbio)     # logistic regression plotting
library(binom)      # binomial confidence intervals
```

##### 2.2 Read Source Files

Each sourced script below aides in either data cleaning (`read_flight_data()`, `center_data()`) or multivariate GLM (`model_comparisonsAIC()`, `get_model_probs()`). Additionally, the function `model_comparisonsAIC()` takes in the path of a generic multi-factor script specific to the GLM family and link function needed to build the predictive models. All aforementioned, sourced scripts are located in the **Rscr** folder.

```r
source_path = paste0(dir,"/Rsrc/")

script_names = c("center_flight_data.R",  # 1 function: center_data()
                 "clean_flight_data.R",   # 1 function: clean_flight_data()
                 "unique_flight_data.R",  # 1 function: create_delta_data()
                 "compare_models.R",      # 1 function: model_comparisonsAIC()
                 "get_Akaike_weights.R")  # 1 function: get_model_probs()

for (script in script_names) {
  path = paste0(source_path, script)
  source(path)
}
```

##### 2.3 Read the Data

The flight performance data read directly below are only from Winter 2020 flight trials. The `read_flight_data()` function standardizes data types and names of the ID, trial type, host plant, flight response, egg-laying response, sex, population, and wing morph inputs. The date, start time, and end time of trails are also converted into datetimes. Variables of interest like wing-to-body ratio are also calculated and centered. Using the `clean_flight_data()` function, all morphology, mass, and flight performance measurements are centered and/or standardized within the `read_flight_data()` function. Then, what is returned is a full dataset (n=758) that includes all bugs collected during Winter 2020 and a subset of the full dataset (n=614) that includes only bugs tested from the Winter 2020 collection.

The `create_delta_data()` function generates the unique dataset by grouping by ID. The function also computes trial differences, percent differences, and averages for variables of interest such as mass, flight response, and egg-laying response. Then, the unique data variables are centered and/or standardized.

```r
data_path = paste0(dir,"/Dispersal/Winter_2020/stats/data/all_flight_data-Winter2020.csv")

data = read_flight_data(data_path) # centers each subset of data
data_all = data[[1]]               # full dataset
data_tested = data[[2]]            # subset of data_all, contains only bugs flight tested

# create the unique dataset
d = create_delta_data(data_tested, remove_bugs_tested_once = FALSE)

# keep all bugs (even bugs only tested once), then re-center
dc = center_data(d, is_not_unique_data = FALSE)
```

Example of a **full dataset** (each row has unique ID and trial type):

```r
data_tested[c(1:2,400:401), c("ID", "trial_type")]
```

```
##      ID trial_type
## 1   114         T1
## 2   318         T1
## 400 316         T2
## 401 416         T2
```

Example of a **unique dataset** (each row has unique ID):

```r
dc[c(1:2,295:296), c("ID", "trial_type")]
```

```
## # A tibble: 4 x 2
## # Groups:   ID [4]
##   ID    trial_type
##   <fct> <list>
## 1 1     <fct [2]>
## 2 2     <fct [2]>
## 3 400   <fct [2]>
## 4 401   <fct [2]>
```

The datatype of the `trial_type` column is a list because when expanded out, it would show `list(T1, T2)`.

#### 2.4 Experimental Effects
##### trial type, days from start, trial start time

To determine how the design of the experiment affected flight response and/or performance, three design factors were modeled: trial type (T1 vs. T2), days from start, and trial start time.

```r
# computed how many times flew yes or no per trial
binary_counts = table(data_tested$flew_b, data_tested$trial_type)[,2:3]

# aggregated by days since trials began and flight response to determine flight prob
dd = aggregate(flew_b ~ days_from_start, data=data_tested, FUN=mean)
```

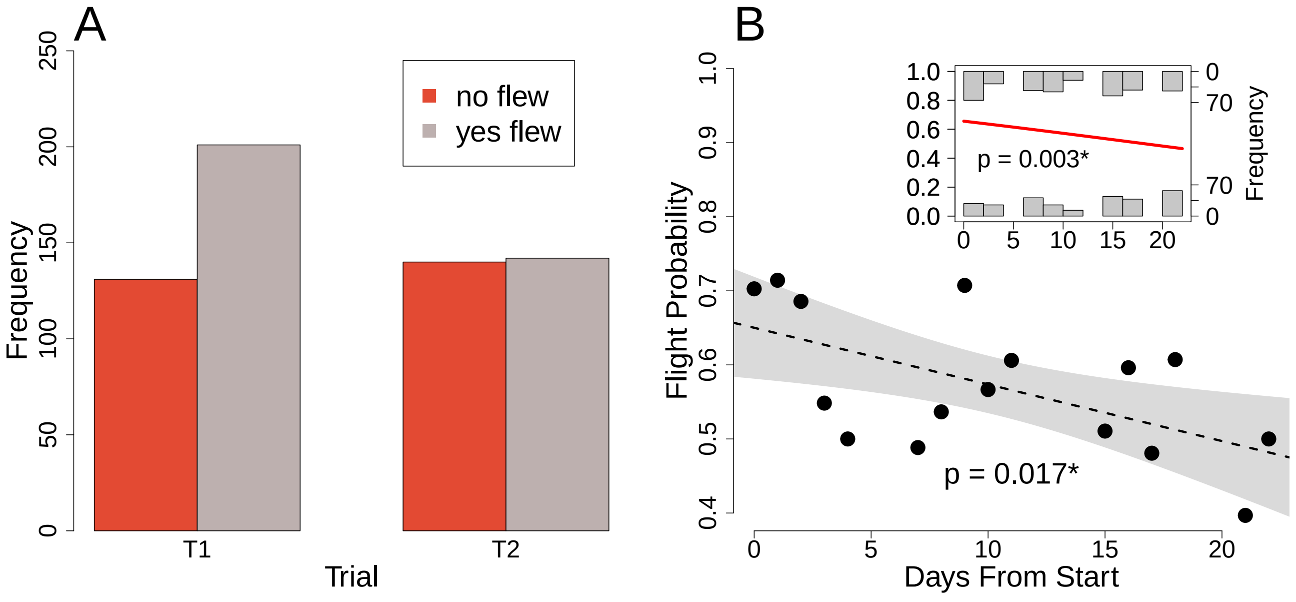

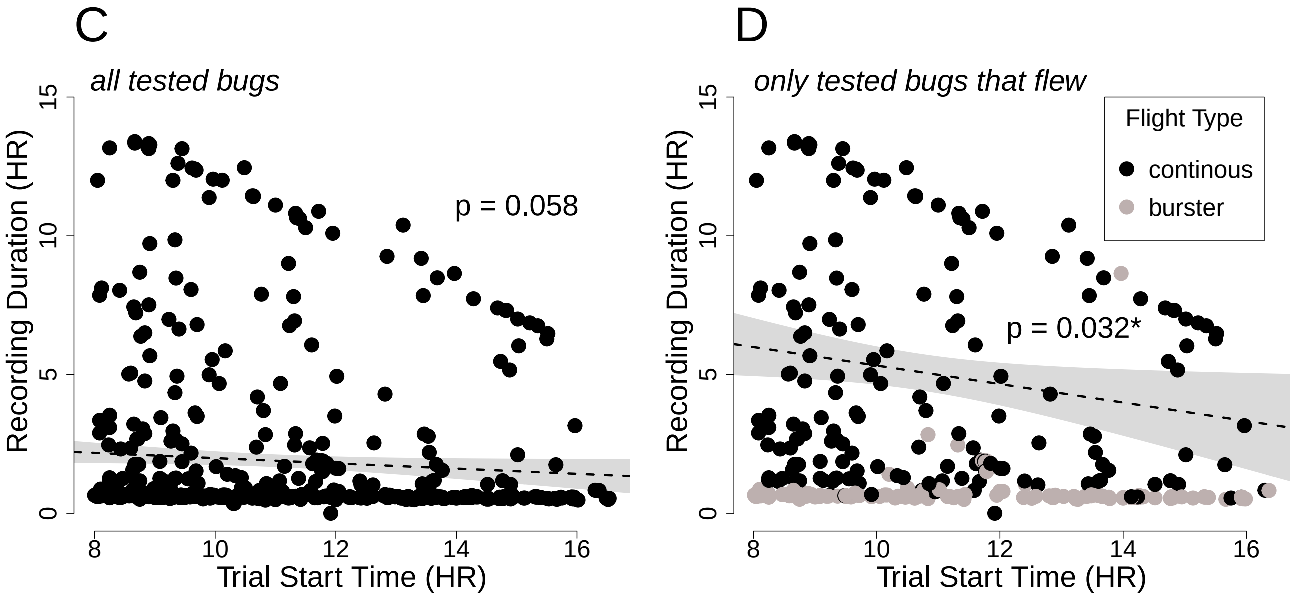

**A & B.** There was a negative effect of day a bug was tested (since the start of trials) on flight probability, but there was a significant effect only when the full dataset is considered. It is not significant for the unique dataset because days from start had to be averaged between trials. This is explored in the next section of the report. **C & D.** There was a negative effect of the trial start time on flight duration but only after removing bugs that did not fly ($p = 0.031$). Continuous flyers are driving this significant relationship (**D**).

#### 2.5 Flight Response Binomial Modeling

To understand SBB flight response, flight response across trials was modeled against sex, host plant, distance from the sympatric zone, wing-to-body ratio, and mass. This was done using the unique dataset.

Because the unique dataset was used, there exist multiple recorded counts of the number of times a SBB flew and did not fly between T1 and T2. For that reason, we used `cbind(num_flew, num_notflew)` when modeling in order to account for all flight successes and failures for each individual.

Finally, we tested whether the data was over-dispersed, which could be resolved using a Quasibinomial:

```
# calculate the confidence interval for the mean of the data (Binomial vs. Quasibinomial)
fit = glm(cbind(num_flew, num_notflew) ~ 1, family = binomial, data = dc)
plogis(confint(fit))
```

```
##      2.5 %    97.5 %
## 0.5191729 0.5976031
```

```
fit_q = glm(cbind(num_flew, num_notflew) ~ 1, family = quasibinomial, data = dc)
plogis(confint(fit_q))
```

```
##      2.5 %    97.5 %
## 0.5108508 0.6056992
```

```
# estimate the dispersion parameter
summary(fit_q)$dispersion
```

```
## [1] 1.464596
```

If the dispersion parameter is close to 1, the data is not over-dispersed, so there is not much of a necessity to apply a Quasibinomial model. Therefore, we selected the family as "binomial".

##### 2.5.1 Average Days Since Start

For the unique dataset, average days since start was computed in order to determine how this experimental factor affected flight response across trials. It proved to not be significant:

```
avg_days_model=glm(cbind(num_flew,num_notflew)~avg_days_c, data=dc, family=binomial)
summary(avg_days_model)
```

```
##
## Call:
## glm(formula = cbind(num_flew, num_notflew) ~ avg_days_c, family = binomial,
##     data = dc)
##
## Deviance Residuals:
##     Min       1Q   Median       3Q      Max
## -1.8429  -1.7825  -0.1593   1.5162   1.5795
##
## Coefficients:
##             Estimate Std. Error z value Pr(>|z|)
## (Intercept)  0.22942    0.08236   2.785  0.00535 **
## avg_days_c   0.01087    0.02354   0.462  0.64430
## ---
## Signif. codes:  0 '***' 0.001 '**' 0.01 '*' 0.05 '.' 0.1 ' ' 1
##
## (Dispersion parameter for binomial family taken to be 1)
##
##     Null deviance: 668.05  on 332  degrees of freedom
## Residual deviance: 667.84  on 331  degrees of freedom
## AIC: 759.17
##
## Number of Fisher Scoring iterations: 3
```

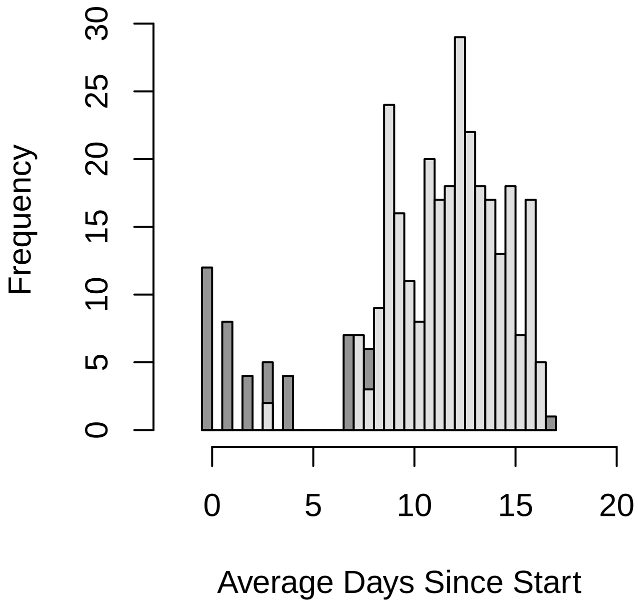

Average days since start accounts for bugs who died before they could be tested twice, which would most likely lead to an early average day value. The rest of the bugs that were tested twice would most likely have a later average day value. This testing regime shapes the bimodal distribution seen in the histogram. Additionally, the advantage of this computed variable is that it controls for the fact that some bugs were tested once late, and some had been tested twice early. In turn, because we randomized test day, when repeated measures for each individual are combined across days, they balance each other out. Thus, average days since start allows the multiple variate models, which control for repeated tests per ID number, to converge, and we can be confident that non-random mortality is not impacting flight response.

##### 2.5.2 Single-Variate Effects sex, mass, wing2body

We used aggregated datasets for single-variate modeling and plotted significant effects below.

```
# tailored variables for plotting
d$mass_block=round(d$avg_mass/0.005)*0.005      # 0.005 g blocks
d$wing2body_block=round(d$wing2body, digits=2)  # 0.01 blocks
d$days_block=round(d$avg_days, digits=0)        # integer blocks
```

```
# aggregated data for plotting
dt=aggregate(flew_prob~sex, data=d, FUN=mean)
dt$trials=c(sum(d$num_flew[d$sex=="F"]+d$num_notflew[d$sex=="F"]),
            sum(d$num_flew[d$sex=="M"]+d$num_notflew[d$sex=="M"]))

ds=aggregate(flew_prob~sex*wing2body_block, data=d, FUN=mean)
ds$n=aggregate(flew_prob~sex*wing2body_block, data=d, FUN=length)$flew_prob

dm=aggregate(flew_prob~sex*mass_block, data=d, FUN=mean)
dm$n=aggregate(flew_prob~sex*mass_block, data=d, FUN=length)$flew_prob

# calculated binomial confidence interval
dt$successes = c(sum(d$num_flew[d$sex=="F"]), sum(d$num_flew[d$sex=="M"]))
dt$CI = binom.confint(dt$successes, dt$trials, methods="exact")
```

```
# sex effect
summary(glm(flew_prob ~ sex, data=ds, family="gaussian"))$coefficients
```

```
##              Estimate Std. Error  t value     Pr(>|t|)
## (Intercept) 0.1858431 0.06354553 2.924566 0.0083810734
## sexM        0.3713285 0.08986695 4.131980 0.0005166983
```

```
# wing-to-body ratio effects split by sex
dsF = ds[ds$sex=="F",] # females
summary(glm(flew_prob ~ wing2body_block, data=dsF, family="gaussian"))$coefficients

dsM = ds[ds$sex=="M",] # males
summary(glm(flew_prob ~ wing2body_block, data=dsM, family="gaussian"))$coefficients
```

```
##                  Estimate Std. Error   t value  Pr(>|t|)
## (Intercept)     -1.015772  0.9521713 -1.066796 0.3138436
## wing2body_block  1.716593  1.3585821  1.263518 0.2381499
##                  Estimate Std. Error   t value     Pr(>|t|)
## (Intercept)     -3.791187  0.7267736 -5.216463 0.0005517947
## wing2body_block  6.140173  1.0250121  5.990342 0.0002049133
```

```
# mass effects split by sex
dmF = dm[dm$sex=="F",] # females
summary(glm(flew_prob ~ mass_block, data=dmF, family="gaussian"))$coefficients

dmM = dm[dm$sex=="M",] # males
summary(glm(flew_prob ~ mass_block, data=dmM, family="gaussian"))$coefficients
```

```
##                 Estimate Std. Error   t value     Pr(>|t|)
## (Intercept)  0.6937214  0.1136749  6.102679 9.138864e-06
## mass_block  -4.6897780  1.1811376 -3.970560 8.967473e-04
##                Estimate Std. Error  t value  Pr(>|t|)
## (Intercept) 0.2997612  0.2465527 1.215810 0.2697179
## mass_block  8.1021284  5.6013019 1.446472 0.1981879
```

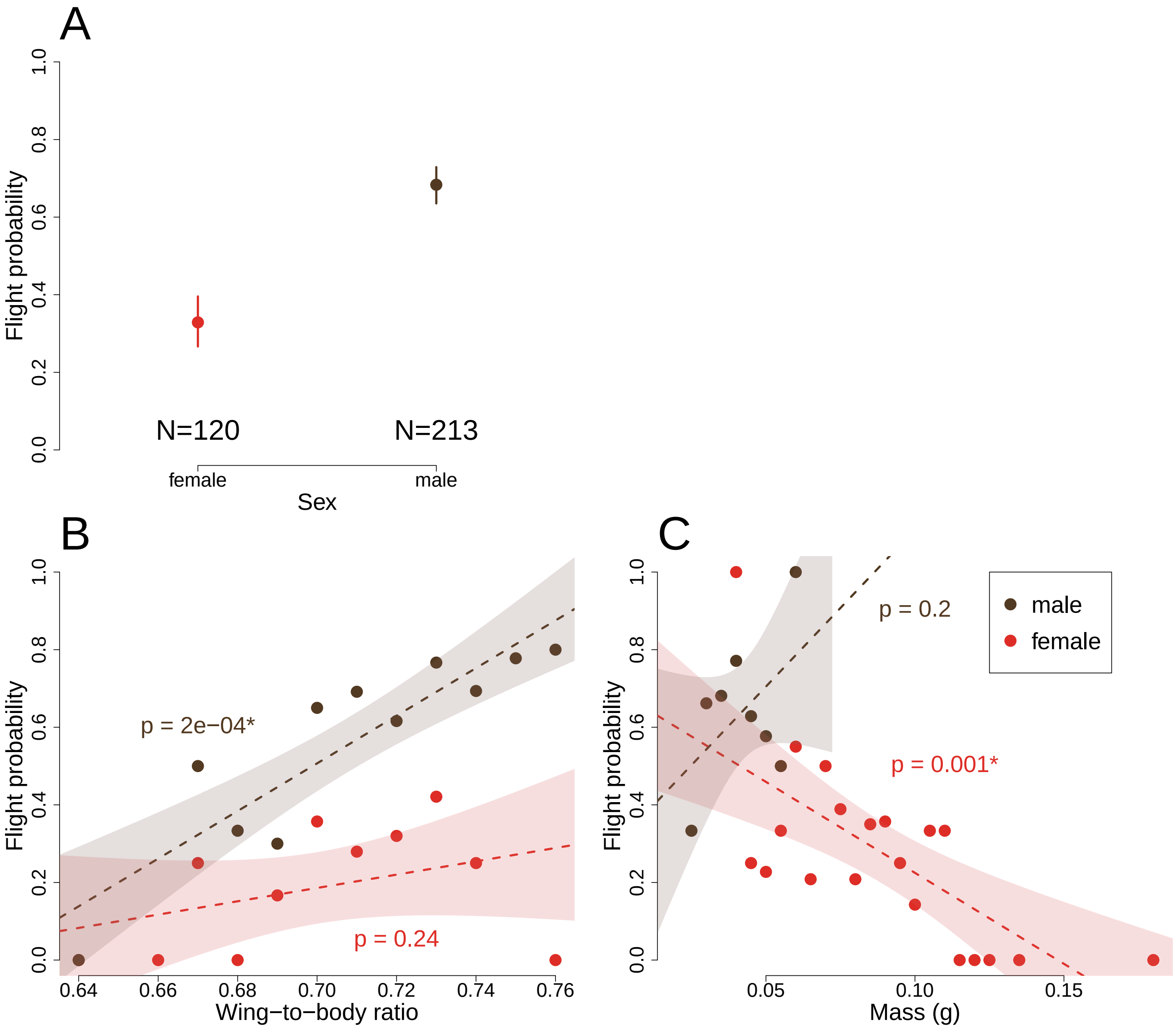

**A.** Males are more than twice as likely to fly than females. **B.** There was a positive effect of wing-to-body ratio for males. **C.** There was a negative effect of mass for females.

##### 2.5.3 Multiple Variate Models
###### sex, mass, wing2body, host plant, distance from sympatric zone

We used the unique dataset for multiple variate modeling. To model flight potential across trials using the unique dataset, flight responses were organized into a matrix. The first column of the matrix counted the number of times a SBB flew across T1 and T2 (e.g. $s$ for 'successes'), and the second column of the matrix counted the number of times a SBB did not fly across T1 and T2 (e.g. $a - s = f$ for 'failures', where $a$ signifies total flight attempts). The matrix was formed using `cbind(s, f)`.

```r
data=data.frame(R1 = dc$num_flew,
                R2 = dc$num_notflew,
                A = dc$host_c,
                B = dc$sex_c,
                C = dc$sym_dist,
                D = dc$avg_mass_logsqrt,
                E = dc$avg_days_c)

model_script = paste0(source_path,"generic models-binomial glm 2R ~ 4-FF + E.R")
model_comparisonsAIC(model_script)
```

```
##         [,1]       [,2]       [,3]
## AICs   683.3791   683.95     684.4483
## models 85         63         50
## probs  0.08873418 0.06669969 0.05198815
##
## m85  glm(formula = cbind(R1, R2) ~ A * D + B * D + C * D + E, family = binomial,
##     data = data)
## m63  glm(formula = cbind(R1, R2) ~ A * D + C * D + B + E, family = binomial,
##     data = data)
## m50  glm(formula = cbind(R1, R2) ~ A * D + B * D + E, family = binomial,
##     data = data)
```

```r
anova(m63, m85, test="Chisq") # Adding B*D does not improve fit
anova(m63, m36, test="Chisq") # Adding C*D does improve fit
```

```
## Analysis of Deviance Table
##
## Model 1: cbind(R1, R2) ~ A * D + C * D + B + E
## Model 2: cbind(R1, R2) ~ A * D + B * D + C * D + E
##   Resid. Df Resid. Dev Df Deviance Pr(>Chi)
## 1       325     580.61
## 2       324     578.04  1   2.5709   0.1088
## Analysis of Deviance Table
##
## Model 1: cbind(R1, R2) ~ A * D + C * D + B + E
## Model 2: cbind(R1, R2) ~ A * D + B + C + E
##   Resid. Df Resid. Dev Df Deviance Pr(>Chi)
## 1       325     580.61
## 2       326     585.11 -1  -4.4988  0.03392 *
## ---
## Signif. codes:  0 '***' 0.001 '**' 0.01 '*' 0.05 '.' 0.1 ' ' 1
```

**Best Fit**

```r
M1 = glm(cbind(num_flew, num_notflew) ~ host_c * avg_mass_logsqrt
          + sym_dist_s * avg_mass_logsqrt + sex_c + avg_days_c, data=dc, family=binomial)
summary(M1)
```

```
##
## Call:
## glm(formula = cbind(num_flew, num_notflew) ~ host_c * avg_mass_logsqrt +
##     sym_dist_s * avg_mass_logsqrt + sex_c + avg_days_c, family = binomial,
##     data = dc)
```

```
##
## Deviance Residuals:
##      Min        1Q    Median        3Q       Max
## -2.54691  -1.08562  -0.03924   1.17713   2.41023
##
## Coefficients:
##                            Estimate Std. Error z value Pr(>|z|)
## (Intercept)                 0.03604    0.11065   0.326  0.74464
## host_c                     -0.14197    0.13044  -1.088  0.27643
## avg_mass_logsqrt           -1.04736    0.88200  -1.187  0.23504
## sym_dist_s                 -0.04098    0.12803  -0.320  0.74890
## sex_c                      -0.46077    0.16797  -2.743  0.00609 **
## avg_days_c                  0.01138    0.02596   0.438  0.66111
## host_c:avg_mass_logsqrt     1.85594    0.59204   3.135  0.00172 **
## avg_mass_logsqrt:sym_dist_s -1.41367    0.68678  -2.058  0.03955 *
## ---
## Signif. codes:  0 '***' 0.001 '**' 0.01 '*' 0.05 '.' 0.1 ' ' 1
##
## (Dispersion parameter for binomial family taken to be 1)
##
##     Null deviance: 668.05  on 332  degrees of freedom
## Residual deviance: 580.61  on 325  degrees of freedom
## AIC: 683.95
##
## Number of Fisher Scoring iterations: 4
```

The best fit model shows two significant interaction terms that interestingly affect SBB flight response across trials: `host_c:avg_mass_logsqrt` and `avg_mass_logsqrt:sym_dist_s`. To explore these terms further and look closer at these island-mainland/native-invasive host plant dynamics, we plotted them. Only part of the code used to generate the plots is shown below.

```
HP = c(1,-1)
SYM = unique(dc$sym_dist_s)
M = seq(min(dc$avg_mass_logsqrt),max(dc$avg_mass_logsqrt), by = 0.05)
c = expand.grid(HP,SYM,M)

eq = function(combo_matrix) {
  effects_col = c()
  for (i in 1:nrow(combo_matrix)) {
    hp=combo_matrix[i,1]
    sym=combo_matrix[i,2]
    ma=combo_matrix[i,3]
    bih = 1.85594
    bis = -1.41367
    total_effect = (bis * sym * ma) + (bih * hp * ma)
    perchange =  (exp(total_effect) - 1) * 100
    effects_col = c(effects_col, perchange)
  }

  return(effects_col)
}
```

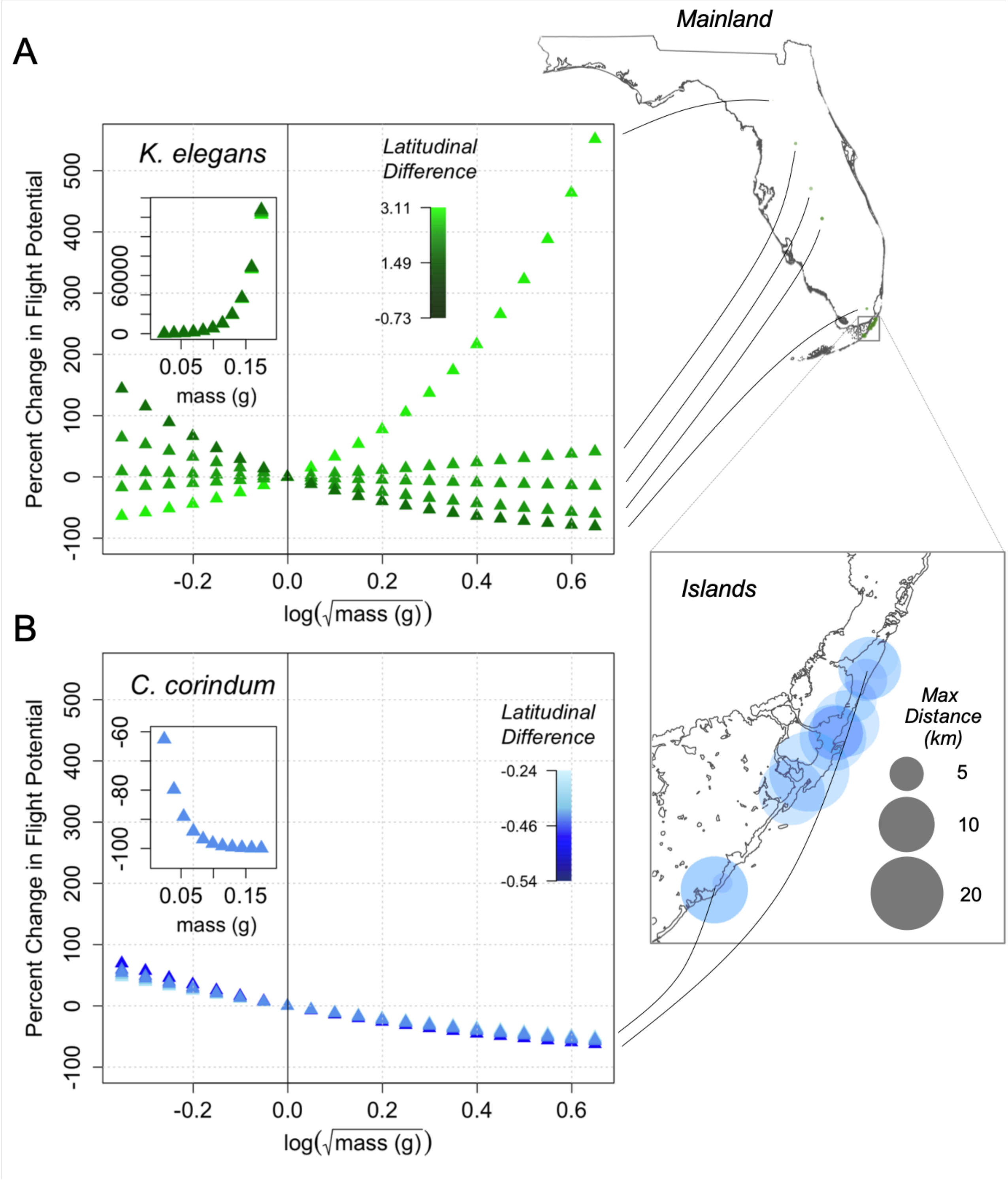

First, we plotted the SBB from golden raintree, the invasive host, which is on the mainland of Florida. Notably, there is a wider range of flight potentials between populations. Distance from the sympatric zone also varies more dramatically with weight changes. The deeper the SBB is in the mainland, then the more likely it will fly if its heaver but if its closer to the islands, then the more likely it will fly if its lighter. Whereas for the islands where the native host plant is located, there is a very narrow range, if any, of flight potential variability between island populations, and there is only a common consistent pattern, where if the SBB from the islands is heavier, then the less likely it will fly, regardless of where it is from on the islands.

This relationship is not only spatially interesting, but it also reveals how weight sensitive SBB can be.

##### 2.5.4 Multiple Variate Models Split By Sex

**Females**

```
data_fem = dc[dc$sex=="F",]
data_fem = center_data(data_fem, is_not_unique_data = FALSE)
```

```
data=data.frame(R1 = data_fem$num_flew,
                R2 = data_fem$num_notflew,
                A = data_fem$host_c,
                B = data_fem$sym_dist,
                C = data_fem$avg_mass_logsqrt,
                D = data_fem$wing2body_logsqrt_i,
                E = data_fem$avg_days_c)

model_script = paste0(source_path,"generic models-binomial glm 2R ~ 4-FF + E.R")
model_comparisonsAIC(model_script)
```

```
##          [,1]       [,2]       [,3]
## AICs   238.8713   239.0635   239.8444
## models 45         25         10
## probs  0.08178418 0.07429069 0.0502761
##
## m45  glm(formula = cbind(R1, R2) ~ A * C + A * D + E, family = binomial,
##     data = data)
## m25  glm(formula = cbind(R1, R2) ~ A * C + D + E, family = binomial,
##     data = data)
## m10  glm(formula = cbind(R1, R2) ~ C + D + E, family = binomial, data = data)
```

```
anova(m25, m45, test='Chisq') #adding A*D does not improve fit
anova(m25, m13, test='Chisq') #adding A*C improves fit
anova(m25, m17, test="Chisq") #adding D improves fit
anova(m25, m45, test="Chisq") #adding D improves fit
```

```
## Analysis of Deviance Table
##
## Model 1: cbind(R1, R2) ~ A * C + D + E
## Model 2: cbind(R1, R2) ~ A * C + A * D + E
##   Resid. Df Resid. Dev Df Deviance Pr(>Chi)
## 1       114     202.11
## 2       113     199.92  1   2.1922   0.1387
## Analysis of Deviance Table
##
## Model 1: cbind(R1, R2) ~ A * C + D + E
## Model 2: cbind(R1, R2) ~ A + C + D + E
##   Resid. Df Resid. Dev Df Deviance Pr(>Chi)
## 1       114     202.11
## 2       115     206.87 -1   -4.764  0.02906 *
## ---
## Signif. codes:  0 '***' 0.001 '**' 0.01 '*' 0.05 '.' 0.1 ' ' 1
## Analysis of Deviance Table
##
## Model 1: cbind(R1, R2) ~ A * C + D + E
## Model 2: cbind(R1, R2) ~ A * C + E
```

```
##   Resid. Df Resid. Dev Df Deviance Pr(>Chi)
## 1       114     202.11
## 2       115     206.35 -1   -4.243  0.03941 *
## ---
## Signif. codes:  0 '***' 0.001 '**' 0.01 '*' 0.05 '.' 0.1 ' ' 1
## Analysis of Deviance Table
##
## Model 1: cbind(R1, R2) ~ A * C + D + E
## Model 2: cbind(R1, R2) ~ A * C + A * D + E
##   Resid. Df Resid. Dev Df Deviance Pr(>Chi)
## 1       114     202.11
## 2       113     199.92  1   2.1922   0.1387
```

##### Best Fit

```
M2 = glm(cbind(num_flew, num_notflew) ~ host_c * avg_mass_logsqrt +  wing2body_logsqrt_i +
           avg_days_c, data=data_fem, family=binomial)
summary(M2)
```

```
##
## Call:
## glm(formula = cbind(num_flew, num_notflew) ~ host_c * avg_mass_logsqrt +
##     wing2body_logsqrt_i + avg_days_c, family = binomial, data = data_fem)
##
## Deviance Residuals:
##     Min       1Q   Median       3Q      Max
## -2.1849  -1.1189  -0.7523   1.1182   2.7357
##
## Coefficients:
##                         Estimate Std. Error z value Pr(>|z|)
## (Intercept)             -0.15913    0.34121  -0.466   0.6409
## host_c                  -0.61037    0.33019  -1.849   0.0645 .
## avg_mass_logsqrt        -2.08700    1.45468  -1.435   0.1514
## wing2body_logsqrt_i     -5.37017    2.66359  -2.016   0.0438 *
## avg_days_c               0.11558    0.04757   2.430   0.0151 *
## host_c:avg_mass_logsqrt  3.02237    1.39976   2.159   0.0308 *
## ---
## Signif. codes:  0 '***' 0.001 '**' 0.01 '*' 0.05 '.' 0.1 ' ' 1
##
## (Dispersion parameter for binomial family taken to be 1)
##
##     Null deviance: 223.66  on 119  degrees of freedom
## Residual deviance: 202.11  on 114  degrees of freedom
## AIC: 239.06
##
## Number of Fisher Scoring iterations: 4
```

The best fit model for female SBB only partially reflects the best fit model for all SBB. Female SBB are sensitive to day changes, which can be a proxi for age. They are also not effected by distance to the local sympatric zone.

##### Males

```
data_male = dc[dc$sex=="M",]
data_male = center_data(data_male, is_not_unique_data = FALSE)
```

```
data=data.frame(R1 = data_male$num_flew,
                R2 = data_male$num_notflew,
                A = data_male$host_c,
                B = data_male$sym_dist,
                C = data_male$avg_mass_logsqrt,
                D = data_male$wing2body_logsqrt_i,
                E = data_male$avg_days_c)

model_script = paste0(source_path,"generic models-binomial glm 2R ~ 4-FF + E.R")
model_comparisonsAIC(model_script)
```

```
##        [,1]       [,2]       [,3]
## AICs   427.3929   427.649    428.1156
## models 105        50         83
## probs  0.08393807 0.07384843 0.05848274
##
## m105     glm(formula = cbind(R1, R2) ~ A * D + B * C + B * D + C * D +
##     E, family = binomial, data = data)
## m50  glm(formula = cbind(R1, R2) ~ A * D + B * D + E, family = binomial,
##     data = data)
## m83  glm(formula = cbind(R1, R2) ~ A * D + B * C + B * D + E, family = binomial,
##     data = data)
```

```
anova(m83, m105, test="Chisq") # adding C*D marginally improves fit
anova(m83, m62, test="Chisq") # adding B*C marginally improves fit
anova(m50, m62, test="Chisq") # adding C does not improve fit
```

```
## Analysis of Deviance Table
##
## Model 1: cbind(R1, R2) ~ A * D + B * C + B * D + E
## Model 2: cbind(R1, R2) ~ A * D + B * C + B * D + C * D + E
##   Resid. Df Resid. Dev Df Deviance Pr(>Chi)
## 1       204     347.73
## 2       203     345.01  1   2.7227  0.09893 .
## ---
## Signif. codes:  0 '***' 0.001 '**' 0.01 '*' 0.05 '.' 0.1 ' ' 1
## Analysis of Deviance Table
##
## Model 1: cbind(R1, R2) ~ A * D + B * C + B * D + E
## Model 2: cbind(R1, R2) ~ A * D + B * D + C + E
##   Resid. Df Resid. Dev Df Deviance Pr(>Chi)
## 1       204     347.73
## 2       205     351.01 -1  -3.2786  0.07019 .
## ---
## Signif. codes:  0 '***' 0.001 '**' 0.01 '*' 0.05 '.' 0.1 ' ' 1
## Analysis of Deviance Table
##
## Model 1: cbind(R1, R2) ~ A * D + B * D + E
## Model 2: cbind(R1, R2) ~ A * D + B * D + C + E
##   Resid. Df Resid. Dev Df Deviance Pr(>Chi)
## 1       206     351.27
```

```
## 2       205     351.01  1  0.25488   0.6137
```

**Best Fit**

```r
M3 = glm(cbind(num_flew, num_notflew)~host_c*wing2body_logsqrt_i +
         sym_dist*wing2body_logsqrt_i + avg_days_c, family=binomial, data=data_male)
summary(M3)
```

```
##
## Call:
## glm(formula = cbind(num_flew, num_notflew) ~ host_c * wing2body_logsqrt_i +
##     sym_dist * wing2body_logsqrt_i + avg_days_c, family = binomial,
##     data = data_male)
##
## Deviance Residuals:
##     Min       1Q   Median       3Q      Max
## -2.6331  -0.7526   0.8309   1.1667   2.0726
##
## Coefficients:
##                              Estimate Std. Error z value Pr(>|z|)
## (Intercept)                   0.46487    0.22478   2.068   0.0386 *
## host_c                       -0.38219    0.18897  -2.023   0.0431 *
## wing2body_logsqrt_i         -15.20652    5.22315  -2.911   0.0036 **
## sym_dist                      0.11229    0.13852   0.811   0.4176
## avg_days_c                   -0.03316    0.03421  -0.969   0.3323
## host_c:wing2body_logsqrt_i   -9.46041    4.27524  -2.213   0.0269 *
## wing2body_logsqrt_i:sym_dist  6.31131    3.02643   2.085   0.0370 *
## ---
## Signif. codes:  0 '***' 0.001 '**' 0.01 '*' 0.05 '.' 0.1 ' ' 1
##
## (Dispersion parameter for binomial family taken to be 1)
##
##     Null deviance: 372.15  on 212  degrees of freedom
## Residual deviance: 351.27  on 206  degrees of freedom
## AIC: 427.65
##
## Number of Fisher Scoring iterations: 4
```

The best fit model for male SBB also only partially reflects the best fit model for all SBB. Host plant and wing-to-body ratio are significant effects while average mass (log-square root transformed) drops off for males.

### 3 Between-Trial Flight Response (T1 vs. T2)

Between-trial flight response analyses determine how differences between trials, such as a SBB's mass or reproductive activity, impact changing flight responses between T1 and T2. Multi-categorical logit modeling was used to analyze changing flight responses, referred to as "flight cases", because the outcomes of the data were no longer binary but instead more than two categories. Flight case is also a nominal response variable, meaning there is no defined order among the response variable categories. See the tables in section 3.4 to read through the encoded categories.

##### 3.1 Read Libraries

The flight case of *J. haematoloma* was analyzed using multi-categorical logit models as implemented in the R package `nnet`. Similar to previous GLM analyses, models were compared using Akaike Information Criterion (AIC), model selection was determined using Akaike weights, and model fit was further evaluated between two models using `anova()`.

Tables were generated using `dplyr` and `kableExtra` and most plots were generated using base R. To generate heatmaps, the `plot.matrix` library was run for easier matrix plotting.

```
library(dplyr)        # data manipulation
library(nnet)         # multinomial modeling
library(kableExtra)   # table formatting
library(plot.matrix)  # enables matrix/heatmap plotting
```

##### 3.2 Read Source Files

The sourced script below aides in creating multi-categorical logit model summary tables, displaying prediction equations of a model, and organizing prediction equation summary matrices for plotting. The tables, after running either `calculate_P2()` or `calculate_P3`, display a model's estimated parameters, standard errors, Wald test statistics, and p values. Those tables can be input into the `get_prediction_eq()` or `get_prediction_eqf()` function to return neatly printed prediction equations. Finally, the summary matrices, after running either `get_significant_models()` or `get_significant_modelsf()`, calculates what `calculate_P2()` or `calculate_P3()` calculates while extracting the p-values of each explanatory variable for each prediction equation of a model. Those p-values are arranged in a matrix and plotted on a heatmap.

The function `model_comparisonsAIC()`, which was run earlier in section 2.2, takes in the path of a generic multi-factor script, but it will now implement the `multinom()` function to build its predictive models. All aforementioned, sourced scripts are located in the **Rscr** folder.

```
script_names = c("multinom_functions.R")  # 6 functions:
                                          # calculate_P2(), calculate_P3(),
                                          # get_prediction_eq()
                                          # get_prediction_eqf()
                                          # get_significant_models(),
                                          # get_significant_modelsf(),

for (script in script_names) {
  path = paste0(source_path, script)
  source(path)
}
```

##### 3.3 Read the Data

```
# this time, only keeping bugs tested twice
d = create_delta_data(data_tested, remove_bugs_tested_once=TRUE)
```

##### 3.4 Encodings & Signs

Below are the categorical encodings and/or signs used for the multi-categorical logit models.

| Flight Case Key | |
|---|---|
| Event | Encoding |
| flew in both trials | 2 |
| flew in T2 only | 1 |
| flew in neither trials | 0 |
| flew in T1 only | -1 |

| Mass Percent Change Key (%) | |
|---|---|
| Event | Sign |
| gained % mass from T1 to T2 | + |
| no % mass change between trails | 0 |
| lost % mass from T1 to T2 | - |

| Host Plant Key | |
|---|---|
| Host | Encoding |
| Golden Rain Tree (GRT) | 1 |
| Balloon Vine (BV) | -1 |

| Sex Key | |
|---|---|
| Sex | Encoding |
| Female | 1 |
| Male | -1 |

##### 3.5 Flight Case Multinomial Modeling

To offer a brief explanation, logit models for nominal response variables pair each category ($j$) with a selected baseline category ($J$). The equation below offers a general prediction equation with a predictor $x$,

$$\log(\frac{\pi_j}{\pi_J}) = \alpha + \beta_j x, j = 1, ..., J - 1$$

In the equation, $\pi$ is the probability of either the category or the baseline, $\alpha$ is the intercept of the model equation, and $\beta$ is the slope, or effect, of the predictor variable. The logit function on the left-hand side of the equation signifies the logarithm of the odds. To calculate the odds of selecting one category over the baseline, each side needs to be exponentiated.

###### 3.5.1 Baseline

The choice of the baseline category is arbitrary. For example, the baseline could be defined as the flight case where a bug flew in neither trial ($Y_i = 0$) and all other categories ($Y_i = 1, -1, \text{or } 2$) would be individually compared to the baseline in the model.

```
# removed any missing values for flight case or mass percent change between trials
df = d[with(d,!is.na(flight_case) & !is.na(mass_per)),]
```

```
# ordered the dataset by ascending mass percent change values
df = df[with(df, order(mass_per)),]

# releveled the flight case factors so as to set 0 as the first level
df$flight_case = relevel(as.factor(df$flight_case), ref = "0")
```

If a new baseline needs to be defined, a prediction equation of a logit model can be rearranged in order to define it. The equation below expresses an arbitrary pair of a and b where a new baseline, b, is being defined.

$$\log(\frac{\pi_a}{\pi_b}) = \log(\frac{\pi_a/pi_J}{\pi_b/pi_J}) = \log(\frac{\pi_a}{\pi_J}) - \log(\frac{\pi_b}{\pi_J})$$
$$= (\alpha_a + \beta_a x) - (\alpha_b + \beta_b x)$$
$$= (\alpha_a - \alpha_b) - (\beta_a x - \beta_b x)$$

Here the new equation for categories a and b has a new intercept parameter $\alpha = (\alpha_a - \alpha_b)$ and slope parameter $\beta = (\beta_a - \beta_b)$.

##### 3.5.2 Compare Models
mass, sex, host plant

```
data = data.frame(R = df$flight_case,
                  A = df$mass_per,
                  B = df$sex_c,
                  C = df$host_c)
model_script = paste0(source_path,"generic multinomial models- multinom 1RF + 3 FF.R")
model_comparisonsAIC(model_script)
```

```
##        [,1]      [,2]      [,3]       [,4]
## AICs   587.5607  591.9016  592.3168   592.4231
## models 4         7         13         12
## probs  0.7141852 0.0815063 0.06622882 0.06280119
##
## m4   multinom(formula = R ~ A + B, data = data, trace = FALSE)
## m7   multinom(formula = R ~ A + B + C, data = data, trace = FALSE)
## m13  multinom(formula = R ~ B * C + A, data = data, trace = FALSE)
## m12  multinom(formula = R ~ A * C + B, data = data, trace = FALSE)
```

```
anova(m4, m7, test="Chisq") # Adding C (host plant) does not improve fit
anova(m4, m8, test="Chisq") # Adding A*B does not improve fit
```

```
## Likelihood ratio tests of Multinomial Models
##
## Response: R
##       Model Resid. df Resid. Dev   Test    Df LR stat.   Pr(Chi)
## 1     A + B       825   569.5607
## 2 A + B + C       822   567.9016 1 vs 2     3 1.659076 0.6460701
## Likelihood ratio tests of Multinomial Models
##
## Response: R
##   Model Resid. df Resid. Dev   Test    Df  LR stat.   Pr(Chi)
## 1 A + B       825   569.5607
## 2 A * B       822   569.4209 1 vs 2     3 0.1398496 0.9866598
```

Here is a potential best fit; however, wing-to-body ratio was not yet considered.

```
M4 = multinom(flight_case ~ mass_per + sex_c, data = df, trace=FALSE)
model_table4 = calculate_P2(M4, "mass_per", "sex_c")
```

```
## AIC:  587.5607
##    (Intercept) mass_per  sex_c DF   SEi   SE1   SE2      zi     z1      z2
## -1      -1.015    0.043 -0.692  9 0.239 0.010 0.203  -4.248  4.390  -3.408
## 1       -6.820   -0.009 -5.626  9 0.183 0.026 0.183 -37.245 -0.348 -30.721
## 2        0.124    0.019 -0.902  9 0.167 0.008 0.159   0.742  2.334  -5.684
##       waldi  wald1   wald2 Pi > |z| P1 > |z| P2 > |z|
## -1   18.049 19.272  11.617    0.000    0.000    0.001
## 1  1387.197  0.121 943.764    0.000    0.728    0.000
## 2     0.551  5.447  32.310    0.458    0.020    0.000
```

Host plant was not a significant predictor, so we reran the model comparisons with wing-to-body ratio included as a predictor with mass percent change and sex.

##### 3.5.3 Compare Models mass, sex, wing2body

```
df$wing2body_c = df$wing2body - mean(df$wing2body) # re-centered the w2b predictor
```

```
data = data.frame(R = df$flight_case,
                  A = df$mass_per,
                  B = df$sex_c,
                  C = df$wing2body_c)
model_script = paste0(source_path,"generic multinomial models- multinom 1RF + 3 FF.R")
model_comparisonsAIC(model_script)
```

```
##        [,1]      [,2]      [,3]
## AICs   582.2678  585.1197  587.133
## models 7         12        13
## probs  0.6671688 0.1603139 0.05858546
##
## m7   multinom(formula = R ~ A + B + C, data = data, trace = FALSE)
## m12  multinom(formula = R ~ A * C + B, data = data, trace = FALSE)
## m13  multinom(formula = R ~ B * C + A, data = data, trace = FALSE)
```

```
anova(m7, m12, test="Chisq") # adding A*C does not improve fit
anova(m7, m13, test="Chisq") # Adding B*C does not improve fit
```

```
## Likelihood ratio tests of Multinomial Models
##
## Response: R
##       Model Resid. df Resid. Dev   Test    Df LR stat.   Pr(Chi)
## 1 A + B + C       822   558.2678
## 2 A * C + B       819   555.1197 1 vs 2     3 3.148182 0.3693379
## Likelihood ratio tests of Multinomial Models
##
## Response: R
```

```
##       Model Resid. df Resid. Dev   Test    Df LR stat.   Pr(Chi)
## 1 A + B + C        822   558.2678
## 2 B * C + A        819   557.1330 1 vs 2     3 1.134887 0.7686596
```

##### 3.5.4 Best Fit

```
M5 = multinom(flight_case ~ mass_per + sex_c + wing2body_c, data = df, trace=FALSE)
model_table5 = calculate_P3(M5)
```

Computer results are rewritten in the table below in order to more legibly show the best fit model's **prediction equations**. Only prediction equations with at least one significant main effect are shown.

| Equation | Effect | Parameter | Estimate | SE | Odds * | Wald | p |
|---|---|---|---|---|---|---|---|
| $P(Yi = flew\ twice)$ | Intercept | $\alpha$ | 1.14 | 0.24 | 3.13 | 22.45 | <0.001 |
| $P(Yi = flew\ in\ T1\ only)$ | $\delta$ Mass % | $\beta_1$ | -0.02 | 0.01 | 0.67 | 7.71 | 0.005 |
| | Sex | $\beta_2\ (F = 1 \vert M = -1)$ | -0.19 | 0.20 | 0.83 | 0.88 | 0.348 |
| | Wing-to-Body | $\beta_3\ (\mu = 0)$ | 4.22 | 10.81 | 1.04 | 0.15 | 0.696 |
| $P(Yi = flew\ twice)$ | Intercept | $\alpha$ | 7.18 | 0.20 | 1312.91 | 1345.71 | <0.001 |
| $P(Yi = flew\ in\ T2\ only)$ | $\delta$ Mass % | $\beta_1$ | 0.02 | 0.02 | 1.49 | 0.92 | 0.336 |
| | Sex | $\beta_2\ (F = 1 \vert M = -1)$ | 5.00 | 0.20 | 148.41 | 642.16 | <0.001 |
| | Wing-to-Body | $\beta_3\ (\mu = 0)$ | 32.82 | 18.71 | 1.39 | 3.08 | 0.079 |
| $P(Yi = flew\ twice)$ | Intercept | $\alpha$ | 0.20 | 0.17 | 1.22 | 1.33 | 0.249 |
| $P(Yi = did\ not\ fly)$ | $\delta$ Mass % | $\beta_1$ | 0.02 | 0.01 | 1.49 | 4.59 | 0.032 |
| | Sex | $\beta_2\ (F = 1 \vert M = -1)$ | -0.76 | 0.17 | 0.47 | 21.07 | <0.001 |
| | Wing-to-Body | $\beta_3\ (\mu = 0)$ | 28.09 | 9.72 | 1.32 | 8.36 | 0.004 |
| $P(Yi = flew\ in\ T1\ only)$ | Intercept | $\alpha$ | -0.94 | 0.24 | 0.39 | 14.96 | <0.001 |
| $P(Yi = did\ not\ fly)$ | $\delta$ Mass % | $\beta_1$ | 0.04 | 0.01 | 2.23 | 18.1 | <0.001 |
| | Sex | $\beta_2\ (F = 1 \vert M = -1)$ | -0.57 | 0.21 | 0.57 | 7.28 | 0.007 |
| | Wing-to-Body | $\beta_3\ (\mu = 0)$ | 23.74 | 12.06 | 1.27 | 3.88 | 0.049 |
| $P(Yi = flew\ in\ T2\ only)$ | Intercept | $\alpha$ | -8.18 | 0.19 | 0.00 | 1919.87 | <0.001 |
| $P(Yi = did\ not\ fly)$ | $\delta$ Mass % | $\beta_1$ | -0.01 | 0.03 | 0.82 | 0.05 | 0.83 |
| | Sex | $\beta_2\ (F = 1 \vert M = -1)$ | -6.95 | 0.19 | 0.00 | 1380.28 | <0.001 |
| | Wing-to-Body | $\beta_3\ (\mu = 0)$ | -6.60 | 18.79 | 0.93 | 0.12 | 0.726 |
| $P(Yi = flew\ in\ T1\ only)$ | Intercept | $\alpha$ | 6.04 | 0.23 | 419.89 | 664.96 | <0.001 |
| $P(Yi = flew\ in\ T2\ only)$ | $\delta$ Mass % | $\beta_1$ | 0.05 | 0.03 | 2.72 | 3.42 | 0.065 |
| | Sex | $\beta_2\ (F = 1 \vert M = -1)$ | 5.19 | 0.22 | 179.47 | 563.3 | <0.001 |
| | Wing-to-Body | $\beta_3\ (\mu = 0)$ | 28.75 | 20.18 | 1.34 | 2.03 | 0.154 |

* Instead of a 1% mass increase, which is relatively small change, the mass percent change estimates were multiplied by 20 before calculating the odds. This transformation better represents experimental observations and offers a more realistic odds.

##### 3.5.5 Visualize Significant Effects in Prediction Equations

From the model summary table above, it appeared that mass, sex, and wing-to-body ratio are significant in most model equations. To better visualize which prediction equations had significant effects, we generated heatmaps for each effect for every model prediction equation.

```r
# defined a run_multinom_model function based on the best fit model
run_multinom_model = function(d) {
  m = multinom(flight_case ~ mass_per + sex_c + wing2body_c, trace=FALSE, data = d)
  model_table = calculate_P3(m, print_table=FALSE)
  return(model_table)
}

# determined which prediction equation effects are significant with a plot
par(mfrow=c(2,2))
mass_per_ML = get_significant_models(19) # % mass
  mtext("Mass", side=3, adj=0, line=0.5, cex=1.4, font=1)
sex_ML = get_significant_models(20) # sex
  mtext("Sex", side=3, adj=0, line=0.5, cex=1.4, font=1)
w2b_ML = get_significant_models(21) # wing2body
  mtext("Wing-to-body", side=3, adj=0, line=0.5, cex=1.4, font=1)
```

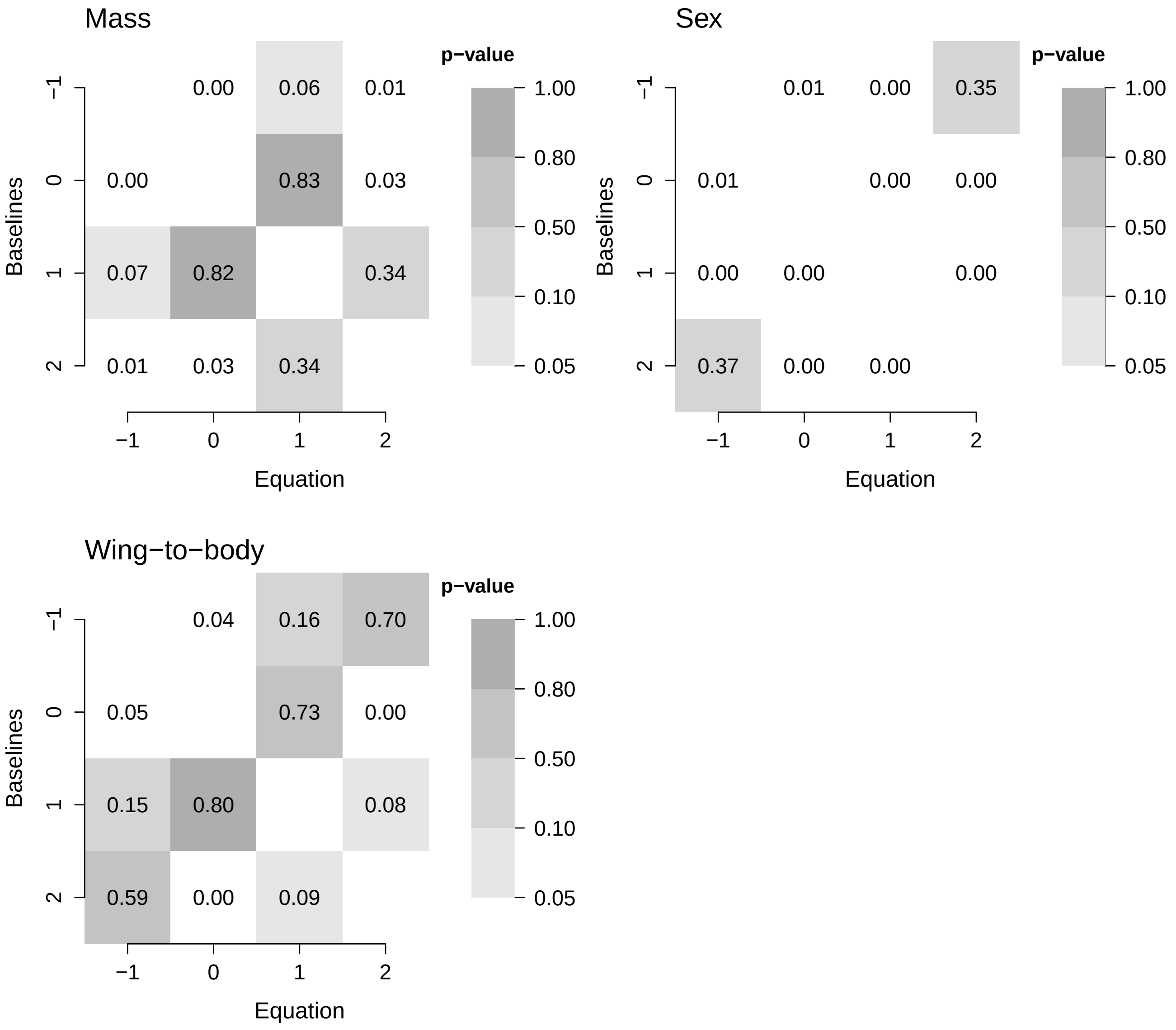

From the heatmap visuals, it becomes clearer to see which effects were significant (i.e. the white panels). Sex, as an effect, was significant in all prediction equations except those comparing the likelihood of flying twice with flying only in T1 only and vice versa. Mass and wing-to-body ratio were less frequently significant but shared significance in common prediction equations such as, those comparing the likelihood of flying twice with not flying at all and the likelihood of flying in T1 only with not flying at all. Mass was also the only significant effect in the prediction equation comparing the likelihood of flying twice with flying only in T1 only and vice versa.

##### 3.5.6 Plot Predicted Probabilities

The predicted probabilities were computed using the `fitted()` function, which extracts fitted values from model objects.

```
pp = fitted(M4) # without wing-to-body ratio
```

```
pp = fitted(M5) # with wing-to-body ratio
```

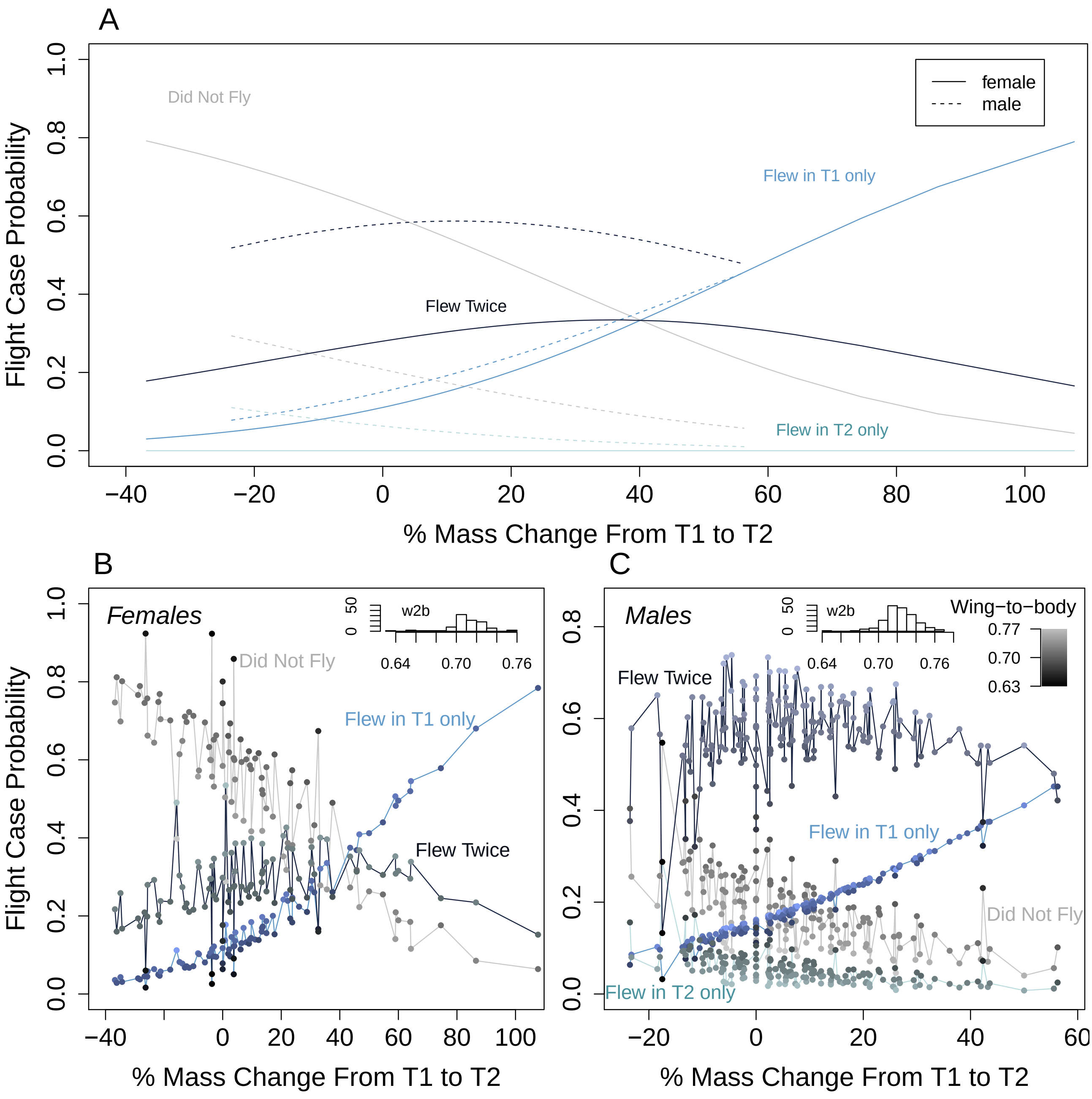

It is noticeable to observe how adding wing-to-body ratio in the model changes the flight case outcome. **B & C.** Each point is colored as a gradient where a low wing-to-body ratio is darker and a higher wing-to-body ratio is lighter. For either sex, there are cases where a really small wing-to-body ratio can supersede any mass changes, making a SBB no longer likely to fly at all in both trials. Meanwhile, a large wing-to-body ratio can do the inverse where the chances of flying twice can spike up and make a SBB most likely to fly twice even at extreme mass changes. Additionally, across each sex, there are flight cases where the stochasticity varies for each case. The blue line and green lines for only flying once do not oscillate much compared to the red and orange lines for flying twice or not flying at all.

##### 3.6 Flight Case Multinomial Modeling (Females Only)

Multi-categorical logit modeling was used to analyze the flight case for females only because females were laying viable and/or inviable eggs during flight trials. These eggs were collected and counted. Additionally, on a female's trial day, we recorded whether we saw or did not see eggs in her bug home. This set of observations was termed as the "egg case", which is encoded below.

###### 3.6.1 Encodings

In addition to the encodings and/or signs mentioned in section 3.4, below are the egg case encodings used for the multi-categorical logit models that explain flight case selection for female SBB only.

Delta Egg Response Key

| Event | Encoding |
|---|---|
| laid eggs in both trials | 2 |
| laid eggs in T2 only | 1 |
| laid eggs in neither trials | 0 |
| laid eggs in T1 only | -1 |

###### 3.6.2 Baseline

```
# filtered for females and removed missing values
df = d[with(d,!is.na(flight_case) & !is.na(mass_per) & !is.na(egg_case) & sex=="F"),]

# ordered the dataset by ascending mass percent change values
df = df[with(df, order(mass_per)),]

# releveled the flight case factors so as to set 0 as the first level.
df$flight_case = relevel(as.factor(df$flight_case), ref = "0")

# no female bug only flew in T2, so dropped factor "1"
df$flight_case = droplevels(df$flight_case)
```

###### 3.6.3 Compare Models

**mass, egg case, host plant**

```
data = data.frame(R = df$flight_case,
                  A = df$egg_case,
                  B = df$mass_per,
                  C = df$host_c)
model_script = paste0(source_path,"generic multinomial models- multinom 1RF + 3 FF.R")
model_comparisonsAIC(model_script)
```

```
##        [,1]      [,2]      [,3]      [,4]       [,5]       [,6]
## AICs   164.3817  165.6054  166.336   167.5638   167.9891   168.3593
## models 7         4         13        11         16         12
## probs  0.3761191 0.2039899 0.1415644 0.07661927 0.06194208 0.0514745
##
## m7   multinom(formula = R ~ A + B + C, data = data, trace = FALSE)
```

```
## m4   multinom(formula = R ~ A + B, data = data, trace = FALSE)
## m13  multinom(formula = R ~ B * C + A, data = data, trace = FALSE)
## m11  multinom(formula = R ~ A * B + C, data = data, trace = FALSE)
## m16  multinom(formula = R ~ B * C + A * B, data = data, trace = FALSE)
## m12  multinom(formula = R ~ A * C + B, data = data, trace = FALSE)
```

```
anova(m4, m7, test="Chisq") # Adding C does not improve fit
anova(m7, m13, test="Chisq") # Adding mass_per*host does not improve fit
```

```
## Likelihood ratio tests of Multinomial Models
##
## Response: R
##       Model Resid. df Resid. Dev   Test    Df LR stat.   Pr(Chi)
## 1     A + B       180   153.6054
## 2 A + B + C       178   148.3817 1 vs 2     2 5.223671 0.0733997
## Likelihood ratio tests of Multinomial Models
##
## Response: R
##       Model Resid. df Resid. Dev   Test    Df LR stat.   Pr(Chi)
## 1 A + B + C       178   148.3817
## 2 B * C + A       176   146.3360 1 vs 2     2 2.045698 0.3595691
```

Host plant was not significant for females as well, so we tested with wing-to-body ratio next.

##### 3.6.4 Compare Models
mass, egg case, wing2body

```
data = data.frame(R = df$flight_case,
                  A = df$egg_case,
                  B = df$mass_per,
                  C = df$wing2body)
model_script = paste0(source_path,"generic multinomial models- multinom 1RF + 3 FF.R")
model_comparisonsAIC(model_script)
```

```
##        [,1]      [,2]      [,3]      [,4]
## AICs   164.5293  164.9831  165.6054  167.7955
## models 7         13        4         12
## probs  0.3174096 0.2529723 0.1853291 0.06199495
##
## m7   multinom(formula = R ~ A + B + C, data = data, trace = FALSE)
## m13  multinom(formula = R ~ B * C + A, data = data, trace = FALSE)
## m4   multinom(formula = R ~ A + B, data = data, trace = FALSE)
## m12  multinom(formula = R ~ A * C + B, data = data, trace = FALSE)
```

```
anova(m4, m7, test="Chisq") # adding wing2body does not improve fit
anova(m7, m13, test="Chisq") # Adding A*C does not improve fit
anova(m7, m12, test="Chisq") # Adding B*C does not improve fit
```

```
## Likelihood ratio tests of Multinomial Models
##
## Response: R
```

```
##        Model Resid. df Resid. Dev   Test    Df LR stat.    Pr(Chi)
## 1      A + B       180   153.6054
## 2 A + B + C       178   148.5293 1 vs 2     2  5.07612 0.07901956
## Likelihood ratio tests of Multinomial Models
##
## Response: R
##        Model Resid. df Resid. Dev   Test    Df LR stat.  Pr(Chi)
## 1 A + B + C       178   148.5293
## 2 B * C + A       176   144.9831 1 vs 2     2 3.546174 0.169808
## Likelihood ratio tests of Multinomial Models
##
## Response: R
##        Model Resid. df Resid. Dev   Test    Df  LR stat.   Pr(Chi)
## 1 A + B + C       178   148.5293
## 2 A * C + B       176   147.7955 1 vs 2     2 0.7337197 0.6929067
```

##### 3.6.5 Best Fit

```r
# same best fit model as the set of model comparisons in section 3.6.3
M6 = multinom(flight_case ~ mass_per + egg_case, data = df, trace=FALSE)
model_table6 = calculate_P2(M6, "mass_per", "egg_case")
```

Computer results are rewritten in the table below in order to more legibly show the best fit model's **prediction equations**. Only prediction equations with at least one significant main effect are shown.

| Equation | Effect | Parameter | Estimate | SE | Odds * | Wald | p |
|---|---|---|---|---|---|---|---|
| $P(Yi = flew\ twice)$ | Intercept | $\alpha$ | 0.41 | 0.42 | 1.51 | 0.92 | 0.338 |
| $P(Yi = did\ not\ fly)$ | $\delta$ Mass % | $\beta_1$ | 0.02 | 0.01 | 1.49 | 4.15 | 0.042 |
| | Egg Case | $\beta_2$ | -1.10 | 0.30 | 0.33 | 13.69 | <0.001 |
| $P(Yi = flew\ in\ T1\ only)$ | Intercept | $\alpha$ | -0.95 | 0.62 | 0.39 | 2.37 | 0.124 |
| $P(Yi = did\ not\ fly)$ | $\delta$ Mass % | $\beta_1$ | 0.04 | 0.01 | 2.23 | 11.49 | 0.001 |
| | Egg Case | $\beta_2$ | -0.53 | 0.38 | 0.59 | 1.97 | 0.161 |

* Instead of a 1% mass increase, which is relatively small change, the mass percent change estimates were multiplied by 20 before calculating the odds. This transformation better represents experimental observations and offers a more realistic odds.

##### 3.6.6 Visualize Significant Effects in Prediction Equations

```r
# defined the run_multinom_model function based on the best fit model
run_multinom_model = function(d) {
  m = multinom(flight_case ~ mass_per + egg_case, trace=FALSE, data = d)
  model_table = calculate_P2(m, "mass_per", "egg_case", print_table=FALSE)
  return(model_table)
}

# visuals of significant effects
par(mfrow=c(1,2))
mass_per_ML = get_significant_modelsf(15) # mass_per
  mtext("Mass", side=3, adj=0, line=0.5, cex=1.5, font=1)
egg_case_ML = get_significant_modelsf(16) # egg_case
  mtext("Egg Case", side=3, adj=0, line=0.3, cex=1.5, font=1)
```

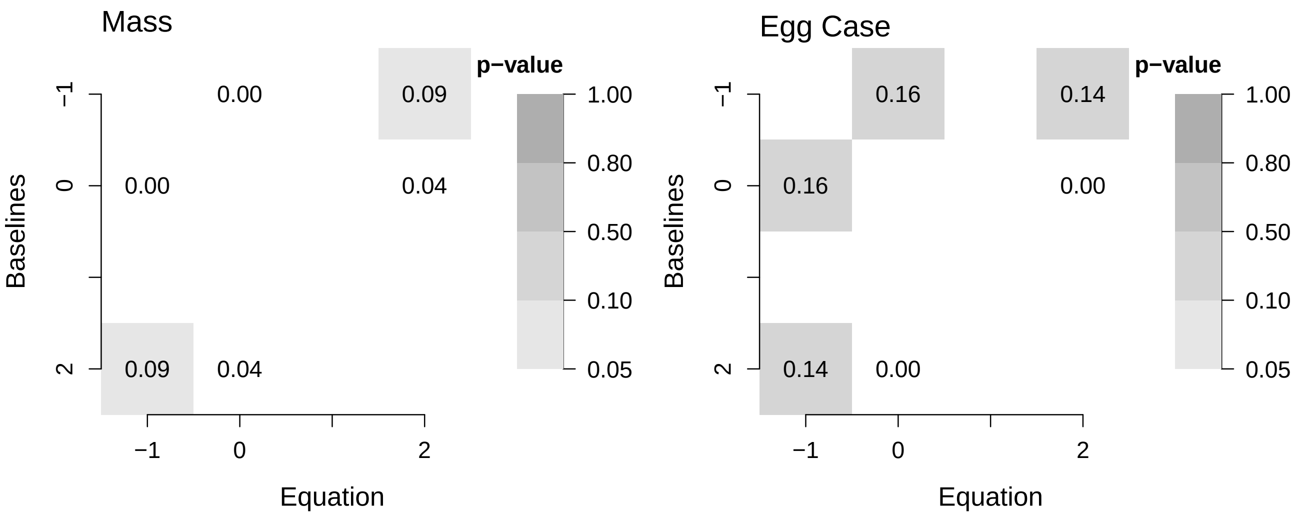

From the heatmap visuals, it becomes clearer to see which effects were significant (i.e. the white panels). Mass and egg case were both significant in the prediction equation that compared the likelihood flying twice with flying in neither trial and vice versa. Mass was also significant in the prediction equation that compared the likelihood of flying only in T1 with flying in neither trial and vice versa. The empty labeled tick mark on the axes signify the missing flight case where a bug flew only in T2. No female flew only in T2 for the Winter 2020 trials.

##### 3.6.7 Plot Predicted Probabilities

The predicted probabilities were computed using the `fitted()` function, which extracts fitted values from model objects.

```
pp = fitted(M6)
```

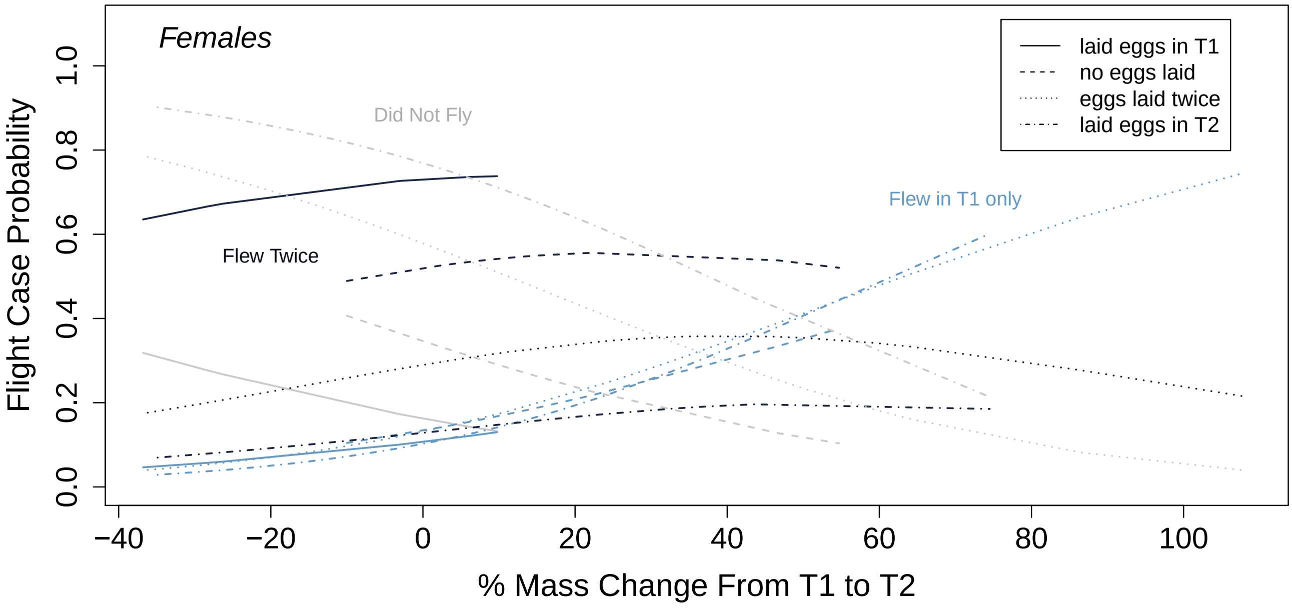

There are several ways to read this graph. First, there are the blue lines, clustered together, that represent the case where a SBB flew only in T1. Whether a SBB flies only once seems to be irregardless of the egg case, which reflects the heatmap plotted in section 3.6.6. However, for flight cases where a SBB flew twice or did not fly at all, the red and black lines are mirror of each other, where egg case does significantly change the flight case outcome. The only situation in which a female SBB is most likely to fly twice would then be the small window where she would have laid no eggs but would have also gained around 40-60% of her original body mass. Additional analyses can be made from this approach.

Another way to read the graph is with the help of an interactive version of this plot available at https://rpubs.com/avbernat/729789. Here, users can select and deselect by egg case, so as to see how each egg case impacts flight case probability. For example, if all egg cases except "no eggs" are deselected, then the user is left to see how a female SBB is most likely to fly twice irrespective of mass changes, just like a male SBB's flight case probability plot.

### Fall 2019 Flight Trials

### 4 Flight Case Predictions

Best fit multinomial models generated for the Winter 2020 flight trials were used to predict the flight case of a SBB during Fall 20019 trials. Actual flight cases were then compared to predicted flight cases in order to assess model accuracy. In turn, we could hypothesize which factors, when considered, could have improved model predictions.

#### 4.1 Read Libraries

All plots were generated using base R and supplemented with the `cvms` package to display confusion matrices.

```
library(cvms)  # cross-validating regressions
```

#### 4.2 Read Source Files

Each sourced script below aides in data cleaning (`clean_flight_data.Fall()`, `create_delta_data.Fall()`) or calculating model accuracy (`calculate_accuracy()`, `get_confusion_matrix()`). All sourced scripts are located in the **Rscr** folder.

```
script_names = c("clean_flight_data-Fall.R",  # 1 function: clean_flight_data.Fall()
                 "unique_flight_data-Fall.R", # 1 function: create_delta_data.Fall()
                 "prediction_accuracy.R",     # 1 function: calculate_accuracy()
                 "confusion_matrix.R")        # 1 function: get_confusion_matrix()

for (script in script_names) {
  path = paste0(source_path, script)
  source(path)
}
```

#### 4.3 Read the Data

The `clean_flight_data.Fall()` function standardizes data types and centers values of the flight response, sex, host plant, wing morph, egg-laying response, average mass, and distance from the sympatric zone. Then, what is returned is a full dataset (n=574) that includes all bugs collected, flight tested, and measured for their morphology during Fall 2020. The full dataset is then filtered to contain SBBs whose masses were measured and who were tested in flight sets 72 to 76 because their experimental design was continuous like the Winter 2020 trial sets. Finally, the `create_delta_data.Fall()` function generates the unique dataset by grouping by ID (n=45). The function also computes trial differences, percent differences, and averages for variables of interest such as mass, flight response, and egg-laying response. Then, the unique data variables are centered.

```
data_path = paste0(dir,"/Dispersal/Winter_2020/stats/data/full_data-Fall2019.csv")
dataFall = clean_flight_data.Fall(data_path)

# extracted sets with an experimental design similar to the Winter tests
```

```r
ongoing_data = dataFall[with(dataFall,!is.na(mass) & set_number > 71),]

# created unique data and sorted by % mass
d = create_delta_data.Fall(ongoing_data)
d = d[with(d, order(mass_per)),]
```

##### 4.4 Plot Predicted Probabilities

The predicted probabilities were calculated using an alternative expression of the multicategory logit model that was represented in section 3.5.1.

$$\pi_j = \frac{e^{\alpha_j + \beta_j x}}{\sum_J e^{\alpha_J + \beta_J x}}, j = 1, ..., J$$

The numerators for each probability $\pi$ varies according to the given flight case $j$, and the probabilities all sum to 1. Meanwhile, the denominator is the same for each flight case.

```r
# stored the best fit model summary table in a new variable
mt = model_table5

# initiated vectors to store predicted probabilities of each flight case
none_pred = c()
T1_vs_none_pred = c()
T2_vs_none_pred = c()
both_vs_none_pred = c()

for (i in 1:nrow(d)) {
  m = d$mass_per[[i]]
  s = d$sex_c[[i]]
  w = d$wing2body_c[i]
  # extracted effects from the best fit model and exponentiated
  top0 = exp(0) # none; equals 1 because it is the baseline
  top1 = exp(mt[1,1] + mt[1,2]*m + mt[1,3]*s + mt[1,4]*w) # T1 rather than none
  top2 = exp(mt[2,1] + mt[2,2]*m + mt[2,3]*s + mt[2,4]*w) # T2 rather than none
  top3 = exp(mt[3,1] + mt[3,2]*m + mt[3,3]*s + mt[3,4]*w) # both rather than none
  bottom = top0 + top1 + top2 + top3
  # calculated predicted probabilities
  none_pred = c(none_pred, top0/bottom)
  T1_vs_none_pred = c(T1_vs_none_pred, top1/bottom)
  T2_vs_none_pred = c(T2_vs_none_pred, top2/bottom)
  both_vs_none_pred = c(both_vs_none_pred, top3/bottom)
  }
```

From the Fall 2019 continuous flight trials, two differences are noticeable in the plots below: 1) the mass percent changes are narrower and 2) there is less stochasticity, which could both be artifacts of fewer bugs having been tested. What becomes more important then is understanding how well the Winter 2020 models do at predicting the Fall 2019 results, which follows in the next section.

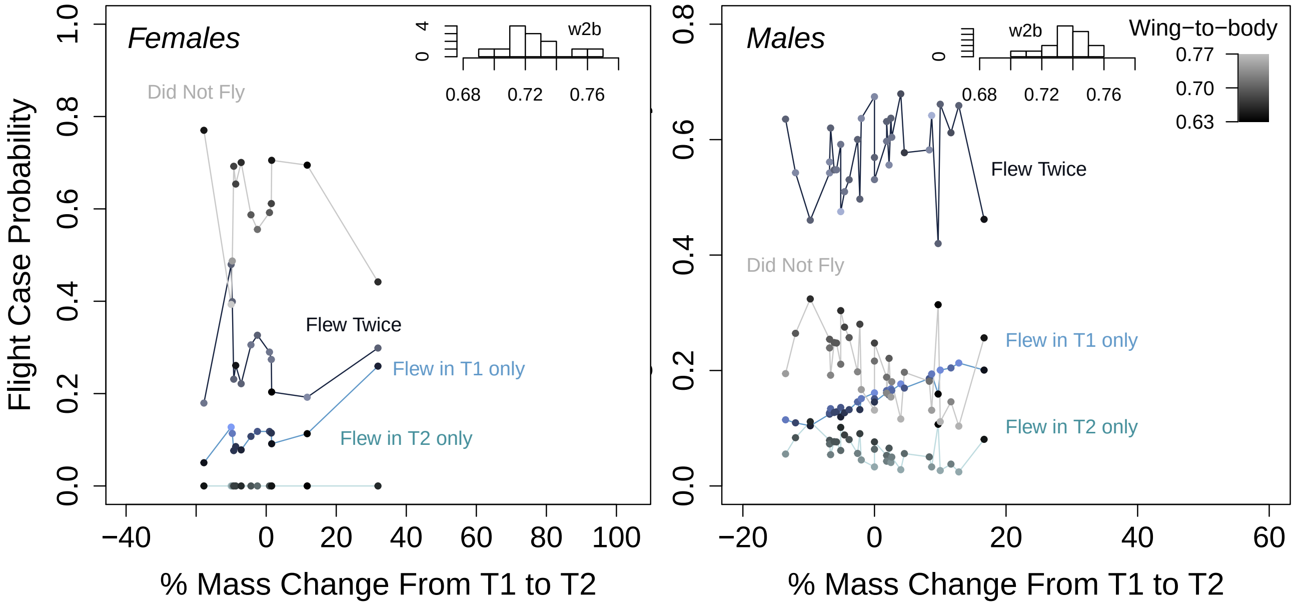

##### 4.5 Overall and Grouped Accuracies

```
probs = round(cbind(none_pred, T1_vs_none_pred, T2_vs_none_pred, both_vs_none_pred),2)

summary_probs = cbind(as.character(d$flight_case), as.character(d$sex), probs)
colnames(summary_probs) = c("event", "sex", "none", "T1", "T2", "both")
df_probs = as.data.frame(summary_probs)

# overall
acc = calculate_accuracy(df_probs,3,6)
paste("Overall prediction accuracy, ", round(acc,2))

# by sex
femdata = df_probs[df_probs$sex=="F",]
maledata = df_probs[df_probs$sex=="M",]

accF = calculate_accuracy(femdata,3,6)
paste("Female prediction accuracy, ", round(accF,2))
accM = calculate_accuracy(maledata,3,6)
paste("Male prediction accuracy, ", round(accM,2))
```

```
## [1] "Overall prediction accuracy,  0.6"
## [1] "Female prediction accuracy,  0.38"
## [1] "Male prediction accuracy,  0.69"
```

Additional, accuracy scores can be measured with the help of the `evaluate()` function embedded in the `get_confusion_matrix` and available through the `cvms` library. For example, sensitivity scores measure true positive frequency and specificity scores measure true negative frequency.

```
acc_table = get_confusion_matrix(df_probs,3,6)
acc_table[,4:5]
```

```
## # A tibble: 1 x 2
##   Sensitivity Specificity
##         <dbl>       <dbl>
## 1       0.293       0.784
```

#### 4.6 Confusion Matrix

The final important performance metric used to determine how well the Winter 2020 models did at predicting the Fall 2019 results is a confusion matrix. A confusion matrix will compare model predictions to actual outcomes in order to provide rates of false positives, false negatives, true positives and true negatives. In this case, the best fit multinomial model for all SBB was one that considered sex, mass percent change, and wing-to-body ratio.

```
confusion_matrix = acc_table$'Confusion Matrix'[[1]]
plot_confusion_matrix(confusion_matrix, add_sums=TRUE,
                      sums_settings = sum_tile_settings(
                        palette = "Oranges",
                        label = "total"),
                      palette="Greys", place_x_axis_above=FALSE,
                      add_zero_shading = FALSE)
```

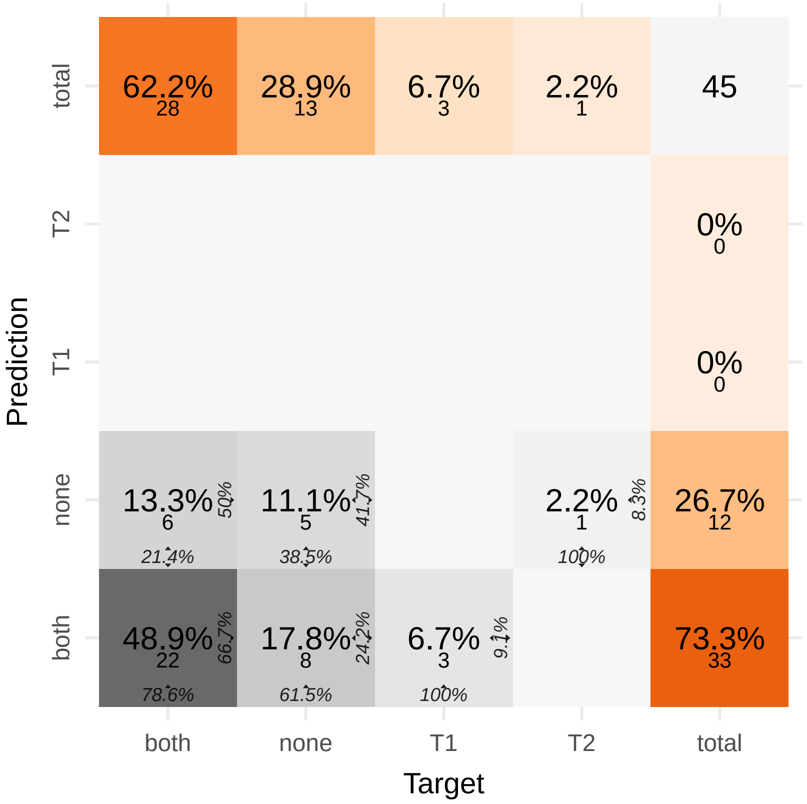

Target cases are the observed cases during Fall trials and prediction cases are the flight cases predicted by the Winter models. Each grey box describes the percentages of false positives, false negatives, true positives, and true negatives for predicting flight cases observed during continuous flight trials in Fall 2019. Each orange box describes the percentages of overall cases that were either observed during Fall trials or were predicted by the model. Based on the confusion matrix ($n = 45$), the Winter 2020 model appears to be overestimating the flight case where a SBB flew twice and underestimating flight cases where a SBB would fly only once.

```
dfem = d[d$sex=="F",]
dfem = dfem[with(dfem, order(mass_per)),]
```

```
mt = model_table6

neither = c()
T1_rather_than_none = c()
both_rather_than_none = c()
```

```
for (i in 1:nrow(dfem)) {
  M = dfem$mass_per[[i]]
  EC = dfem$egg_diff[[i]]
  top0 = exp(0)
  top1 = exp(mt[1,1] + mt[1,2]*M + mt[1,3]*EC)
  top2 = exp(mt[2,1] + mt[2,2]*M + mt[2,3]*EC)
  bottom = top0 + top1 + top2
  neither = c(neither, top0/bottom)
  T1_rather_than_none = c(T1_rather_than_none, top1/bottom)
  both_rather_than_none = c(both_rather_than_none, top2/bottom)
}
```

##### 4.6.1 Plot Predicted Probabilities

The predicted probabilities were calculated using an alternative expression of the multicategory logit model as described in section 4.4.

```
probs = round(cbind(neither, T1_rather_than_none, both_rather_than_none),2)

summary_probs = cbind(as.character(dfem$flight_case), as.character(dfem$egg_diff), probs)
colnames(summary_probs) = c("event", "egg_diff", "none", "T1", "both")

egg2 = c(1,2,3,5,6,7,9,10,11,13)
noegg = c(4,8,12)

dataframe = as.data.frame(summary_probs)
dataframe$egg_cat = c(2,2,2,0,2,2,2,0,2,2,2,0,2)
```

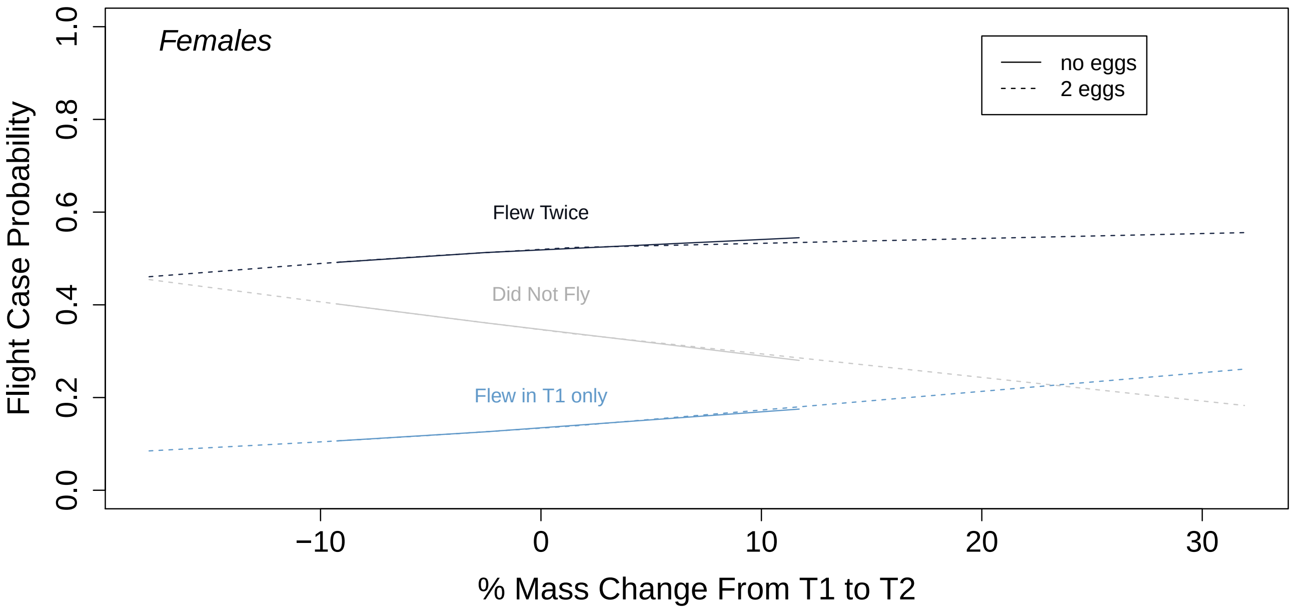

From the Fall 2019 continuous flight trials, two differences are noticeable in the plots below: 1) there were only two egg cases (laid twice or no eggs) and 2) egg cases seem to make no noticeable differences within a given flight case.

Because mass but not egg case seems to be driving flight case outcome for Fall 2019 female SBB, this can already signal that our Winter 2020 best fit model would not necessarily be a reliable predictor of flight case. This is confirmed in the next sections using performance metrics.

###### 4.6.2 Overall and Grouped Accuracies

```
accF_eggs = calculate_accuracy(dataframe,3,5)
paste("Female prediction accuracy for mass diff and egg model, ", round(accF_eggs,2))
```

```
## [1] "Female prediction accuracy for mass diff and egg model,  0.46"
```

```
acc_table = get_confusion_matrix(dataframe,3,5)
acc_table[,4:5]
```

```
## # A tibble: 1 x 2
##   Sensitivity Specificity
##         <dbl>       <dbl>
## 1       0.333       0.667
```

###### 4.6.3 Confusion Matrix

```
confusion_matrix = acc_table$'Confusion Matrix'[[1]]
plot_confusion_matrix(confusion_matrix, add_sums=TRUE,
                      sums_settings = sum_tile_settings(
                        palette = "Oranges",
                        label = "total"),
                      palette="Greys", place_x_axis_above=FALSE,
                      add_zero_shading = FALSE)
```

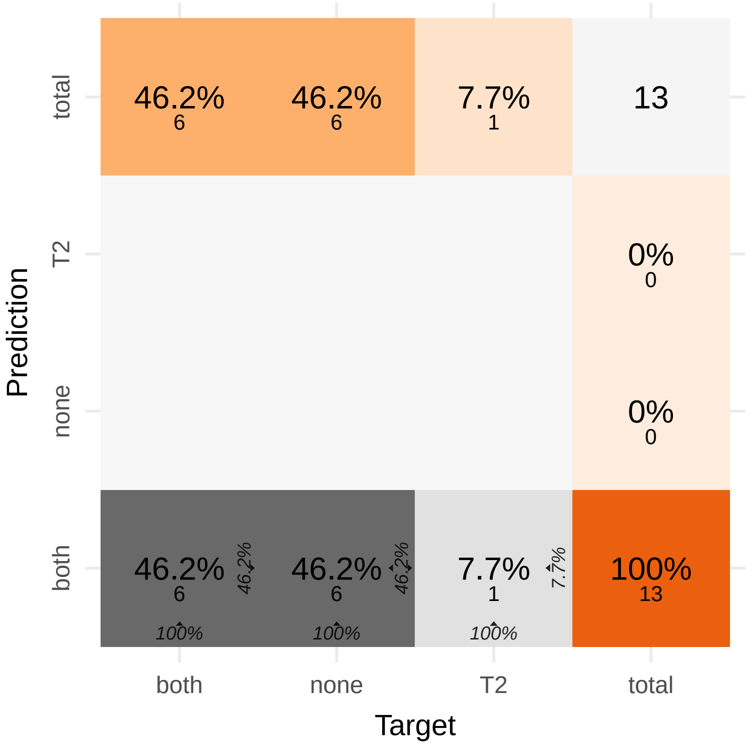

The best fit model ($n = 13$) for female SBB tested in Winter 2020 included mass percent change and egg case. This model, based on the confusion matrix, also largely overestimated the flight case where females flew twice and largely underestimated the remaining flight cases. Here, it becomes evident that our best fit models would need more factors to be more accurate and sensitive to new data. It is possible that other, unmeasured and untested factors could better explain a flight case outcome, such as age or thorax muscle mass weight.
